# Representations of context and context-dependent values in vmPFC compete for guiding behavior

**DOI:** 10.1101/2021.03.17.435844

**Authors:** Nir Moneta, Mona M. Garvert, Hauke R. Heekeren, Nicolas W. Schuck

## Abstract

Value representations in ventromedial prefrontal-cortex (vmPFC) are known to guide the choice between options. But the value of an option can be different in different task contexts. Goal-directed behavior therefore requires to know the current context and associated values of options, and to flexibly switch between value representations in a task-dependent manner. We tested whether task-relevant and -irrelevant values influence behavior and asked whether both values are represented together with context signals in vmPFC. Thirty-five participants alternated between tasks in which stimulus color or motion predicted rewards. As expected, neural activity in vmPFC and choices were largely driven by task-relevant values. Yet, behavioral and neural analyses indicate that participants also retrieved the values of irrelevant features, and computed which option would have been best in the alternative context. Investigating the probability distributions over values and contexts encoded in multivariate fMRI signals, we find that vmPFC maintains representations of the current context, i.e. task state, the value associated with it, and the hypothetical value of the alternative task state. Crucially, we show that evidence for irrelevant value signals in vmPFC relates to behavior on multiple levels, competes with expected value signals, and interacts with task state representations. Our results thus suggest that different value representations are represented in parallel and imply a link between neural representations of task states, their associated values and their influence on behavior. This sheds new light on vmPFC’s role in decision making, bridging between a hypothesized role in mapping observations onto the task states of a mental map, and computing value expectations for alternative states.

## Introduction

Decisions are always made within the context of a given task. Even a simple choice between two apples will depend on whether the task is to find a snack, or to buy ingredients for a cake, for which different apples might be best. In other words, the same objects can yield different outcomes in different task contexts. This could complicate the computations underlying the retrieval of learned values during a decision, since outcome expectations from the wrong context might exert influence on the neural value representation of the available options.

Much work has studied how the reward a choice will yield *in a given task context* is at the core of decisions [e.g. 1]. Most prominently, previous studies have shown in a variety of species that the ventromedial prefrontal cortex (vmPFC) represents this so-called expected value (EV) [2–7], and thereby plays a crucial role in determining choices [8]. It is also known that the brain’s attentional control network enhances the processing of features that are relevant given the current task context or goal [9, 10], and that this helps to shape which features influence EV representations in vmPFC [11–15]. Moreover, the vmPFC seems to also represent the EV of different features in a common currency [16, 17]; and is involved in integrating the expectations from different reward predicting features of the same object [18–21]. It remains unclear, however, how context-*irrelevant* value expectations of available features, i.e. rewards that would be obtained in a different task-context, might affect neural representations in vmPFC, and whether such “undue” influence of irrelevant value expectations can lead to wrong choices. Notably, even when relevant value information dominates choices and vmPFC activity, irrelevant values could still lead to subtle effects on vmPFC activation patterns and behavior.

This is particularly relevant because we often have to do more than one task within the same environment, such as shopping in the same supermarket for different purposes. Thus we have to switch between the values that are relevant in the different contexts. This can lead to less than perfect separation between task contexts/goals and result in processing of task-irrelevant aspects. In line with this idea, several studies have shown that decisions are influenced by contextually-irrelevant information, and traces of the distracting features have been found in several cortical regions, for instance areas responsible for task execution [22–26]. Similarly, task-irrelevant valuation has been shown to influence attentional selection [27] as well as activity in posterior parietal [28] or ventromedial prefrontal cortex [29]. This raises the possibility that in addition to its well known role in signaling values, vmPFC could also represent different values that occur in different task contexts during choice.

If that is the case, the neural representation of context might play a major role in gating context-dependent values in vmPFC. We therefore hypothesised – in line with previous work [30–33] – that vmPFC would also encode the task context, and that a stronger activation of the relevant task-context will enhance the representation of task-relevant values. To test this idea, we investigated whether vmPFC activation is influenced by multiple task-dependent values during choice, and studied how these representations influence decisions, interact with the encoding of the relevant task-context, and with each other. Such a multifaceted representation of multiple values and task contexts within the same region would reconcile work that emphasizes the role of choice value representations in vmPFC and OFC with work which emphasizes the encoding of other aspects of the current task [34–38], in particular of so-called task states [30–33], within the same region [see also, 39, 40].

Note that knowing the current context alone will not immediately resolve which value of two presented options should be represented, similar to how knowing what you are shopping for (cake or snack) will not answer which of the available apples you should pick. We therefore propose that context/task state representations influence value computations in the vmPFC, such that a state representation triggers a comparison between the values of options as they would be expected in the represented state/context. In consequence, the value of the option that *would be best in the activated state* will become represented, and partial co-activation of different possible states could therefore lead to value representations that can refer to different choices (the value of the apple best for snacking and the value of the apple best for baking, even if those are different apples). Moreover, this assumes that context-specific value codes will relate to the strength of the respective state representations within the same region. An alternative view in which state representations do not impact value computations would assume that activated values would always refer to the choice one is going to make in the present context (how valuable the apple chosen for snaking would be for baking).

We investigated these questions using a multi-feature choice task in which different features of the same stimulus predicted different outcomes and a task-context cue modulated which feature was relevant. Based on the above reviewed evidence of neural processing of irrelevant features and values [e.g., 24, 29], we hypothesized that values arising from relevant and irrelevant contexts would influence the vmPFC representation, specifically the expected values of each context. Moreover, we tested whether different possible EVs were integrated into a single value representation or processed in parallel. The former would support a role of the vmPFC for representing *only* the EV of choice, whereas the latter would indicate that the vmPFC encodes several aspects of a complex task structure, including the expected value of one’s choice in the currently relevant context, but also the hypothetical value in the presently irrelevant context.

## Results

### Behavioral results

Thirty-five right-handed young adults (18 women, *μ_age_* = 27.6, *σ_age_* = 3.35, see Methods for exclusions) were asked to judge either the color (context 1) or motion direction (context 2) of moving dots on a screen (random dot motion kinematogramms, [e.g. 41]). Four different colors and motion directions were used. Before entering the MRI scanner, participants performed a stair-casing task in which participants first received a cue that instructed them which feature (a color or direction) will be the target of the current trial. Then participants had select the matching stimulus from two random dot motion stimuli (see Fig. S1c). Motion-coherence and the speed which dots changed from grey to a target color were adjusted such that the different stimulus features could be discriminated equally fast, both within and between contexts (i.e. Color / Motion). As intended, this led to significantly reduced differences in reaction times (RTs) between the eight stimulus features (*t*_(34)_ = 7.29, *p* < .001, Fig.1a), also when tested for each button separately (*t*_(34)_ =Left: 6.52, Right: 7.70, *p*s< .001, Fig. S1d).

**Figure 1:**
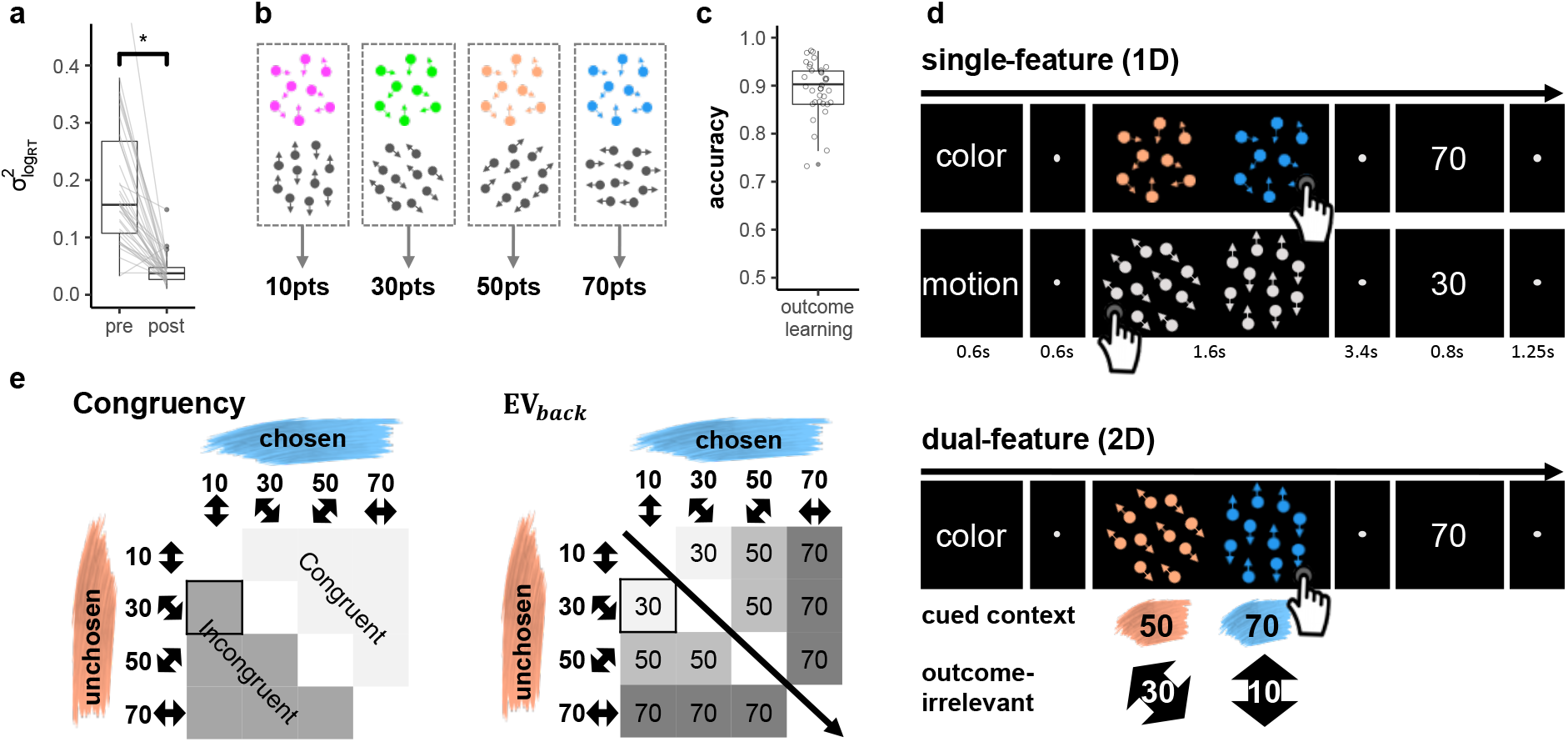
Task and Design. **a.** Staircasing procedure reduced differences in detection speed between features. Depicted is the variance of reaction times (RTs) across different color and motion features (y axis). While participants’ RTs were markedly different for different features before staircasing (pre), a significant reduction in RT differences was observed after the procedure (post). The staircasing procedure was performed before value learning. RT-variance was computed by summing the squared difference of each feature’s RT and the general mean RT per participant. Center line in each box represents the mean and the box limits the first and third quartiles. *N* = 35, *p* < .001. **b.** The task included eight features, four color and four motion directions. After the stair-casing procedure, a specific reward was assigned to each motion and each color, such that one feature from each of the contexts had the same value as it was associated with the same reward. Feature values were counterbalanced across participants. **c.** Participants were trained on feature values shown in (b) and achieved near ceiling accuracy in choosing the highest valued feature afterwards (*μ* = .89, *σ* = .06). Center line in each box represents the mean and the box limits the first and third quartiles. **d.** Single- and dual-feature trials (1D, 2D, respectively). Each trial started with a cue of the relevant context (Color or Motion, 0.6s), followed by a short fixation circle (0.6s). Participants were then presented with a choice between two clouds (1.6s). Each cloud had only one feature in 1D trials (colored dots, but random motion, *or* directed motion, but gray dots, top) and two features for 2D trials (motion *and* color, bottom). Participants were instructed to make a decision between the two clouds based on the cued context and ignore the other. Choices were followed by a fixation period (3.4s) and the value associated with the chosen cloud’s feature of the cued context (0.8s). After another short fixation (1.25s) the next trial started. **e.** Variations in values irrelevant in the present task context of a 2D trial. For each feature pair (e.g. blue and orange), all possible context-irrelevant feature-combinations were included in the task, except the same feature on both sides. Congruency (left): trials were separated into those in which the irrelevant features favored the same choice as the relevant features (congruent trials), or not (incongruent trials). EV_back_ (right): based on this factor, the trials were characterized by different hypothetically expected values of the contextually-irrelevant features, i.e. the maximum value of both irrelevant features. Crucially, EV, EV_back_ and Congruency were orthogonal by design. The example trial presented in (d, bottom) is highlighted.

Only then, participants learned to associate each color and motion feature with a fixed number of points (10, 30, 50 or 70 points), whereby one motion direction and one color each led to the same reward (counterbalanced across participants, Fig.1b). To this end, participants had to make a choice between clouds that had only one feature-type, while the other feature type was absent or ambiguous (clouds were grey in motion-only clouds and moved randomly in color clouds). To encourage mapping of all features on a unitary value scale, choices in this part (and only here) also had to be made between contexts (e.g. between a green and a horizontal-moving cloud). At the end of the learning phase, participants achieved near-ceiling accuracy in choosing the cloud with the highest valued feature (*μ* = .89, *σ* = 0.06, t-test against chance: *t*_(34)_ = 41.8, *p* < .001, Fig. 1c), also when tested separately for color, motion and across context (*μ* = .88, .87, .83, *σ* = .09, .1, .1, t-test against chance: *t*_(34)_ = 23.9, 20.4,19.9, *p*s< .001, respectively, Fig. S1e). Once inside the MRI scanner, one additional training block ensured changes in presentation mode did not induce feature-specific RT changes (*F*_(7,202)_ = 1.06, *p* = 0.392). These procedures made sure that participants began the main experiment inside the MRI scanner with firm knowledge of feature values; and that RT differences would not reflect perceptual differences, but could be attributed to the associated values. Additional information about the pre-scanning phase can be found in Online Methods and in Fig.S1.

During the main task, participants had to select one of two dot-motion clouds. In each trial, participants were first cued whether a decision should be made based on color or motion features, and then had to choose the cloud that would lead to the largest number of points. Following their choice, participants received the points corresponding to the value associated with the chosen cloud’s relevant feature. To reduce complexity, the two features of the *cued task-context* always had a value difference of 20, i.e. the choices on the cued context were only between values of 10 vs. 30, 30 vs. 50 or 50 vs. 70. One third of the trials consisted of a choice between single-feature clouds of the same context (henceforth: 1D trials, Fig.1d, top). All other trials were dual-feature trials, i.e. each cloud had a color *and* a motion direction at the same time (henceforth: 2D trials, Fig.1d bottom), but the context indicated by the cue mattered. Thus, while 2D trials involved four features in total (two clouds with two features each), only the two color *or* two motion features were relevant for determining the outcome. The cued context stayed the same for a minimum of four and a maximum of seven trials. Importantly, for each comparison of relevant features, we varied which values were associated with the features of the *irrelevant* context, such that each relevant value was paired with all possible irrelevant values (Fig.1e). Consider, for instance, a color trial in which the color shown on the left side led to 50 points and the color on the right side led to 70 points. While motion directions in this trial did not have any impact on the outcome, they might nevertheless influence behavior. Specifically, they could favor the same side as the colors or not (Congruent vs Incongruent trials, see Fig.1e left), and have larger or smaller values compared to the color features (Fig.1e right).

We investigated the impact of these factors on RTs in correct 2D trials, where the extensive training ensured near-ceiling performance throughout the main task (*μ* = 0.91, *σ* = 0.05, t-test against chance: *t*_(34)_ = 48.48, *p* < .0001, Fig.2a). RTs were log transformed to approximate normality and analysed using mixed effects models with nuisance regressors for choice side (left/right), time on task (trial number), differences between attentional contexts (color/motion) and number of trials since the last context switch (all nuisance regressors had a significant effect on RTs in the baseline model, all *p*s< 0.03). We used a hierarchical model comparison approach to assess the effects of (1) the objective value of the chosen option (or: EV), i.e. points associated with the features on the cued context; (2) the maximum points that could have been obtained if the irrelevant features were the relevant ones (the expected value of the background, henceforth: EV_back_, Fig 1e left), and (3) whether the irrelevant features favored the same side as the relevant ones or not (Congruency, Fig. 1e right). Any effect of the latter two factors would indicate that outcome associations that were irrelevant in the current context nevertheless influence behavior, and therefore could be represented in vmPFC.

**Figure 2:**
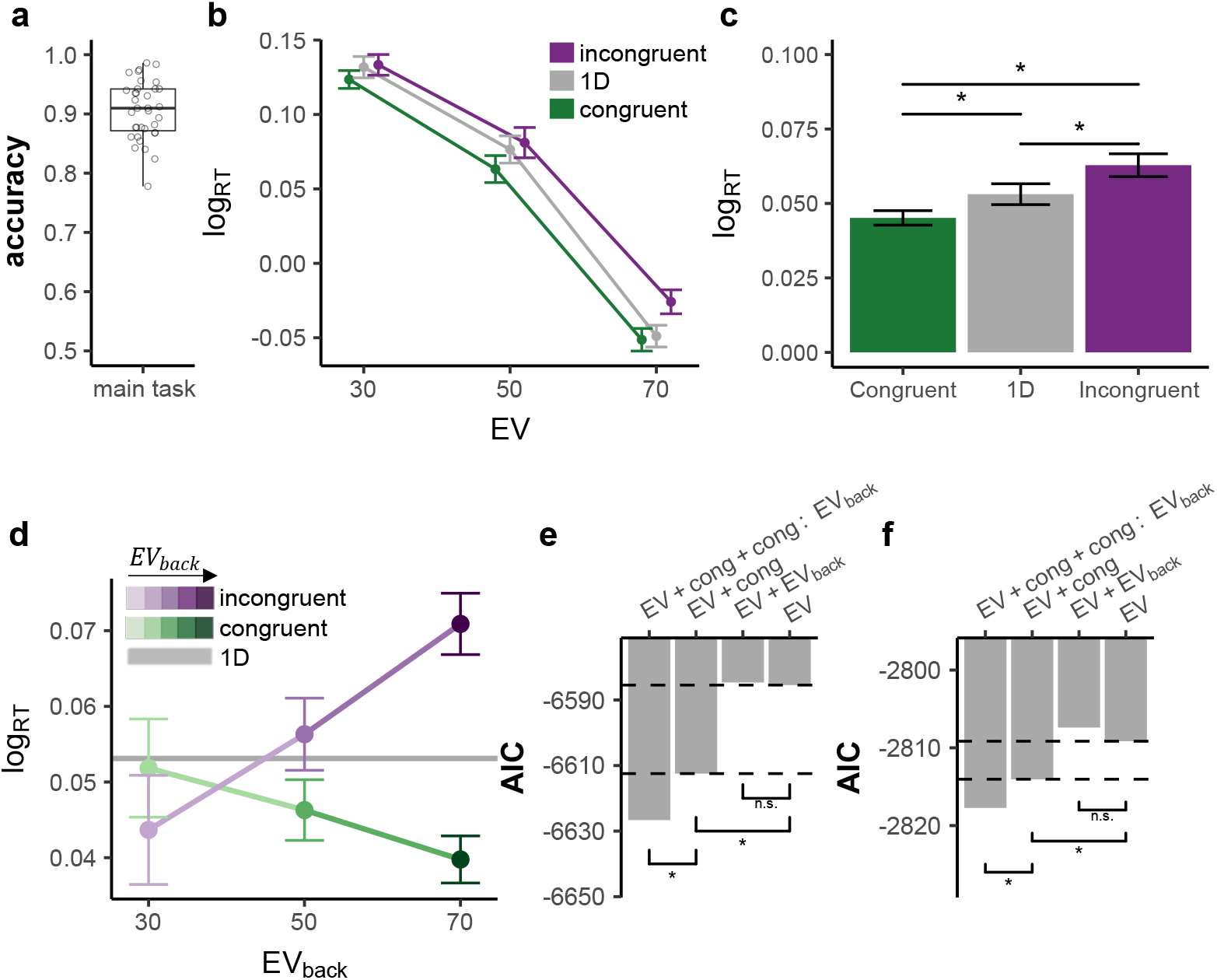
Behavioral results. **a.** Participants were at near-ceiling performance throughout the main task, *μ* = 0.905, *σ* = 0.05. Center line in each box represents the mean and the box limits the first and third quartiles. **b.** Participants reacted faster the higher the EV (x-axis) and slower to incongruent (purple) compared to congruent (green) trials. An interaction of EV × Congruency indicated stronger Congruency effect for higher EV (*p* = .037), but did not replicate in the replication sample (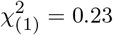, *p* = .63). RT for 1D trials is plotted in gray, see formal tests in panel c. Error bars represent corrected within subject SEMs [42, 43]. **c.** Participants reacted slower to incongruent compared to 1D trials (p = .008) and faster to congruent compared to either 1D (*p* = .017) or incongruent trials (*p* < .001). **d.** The Congruency effect was modulated by EV_back_, i.e. the more participants could expect to receive from the ignored context, the slower they were when the contexts disagreed and respectively faster when contexts agreed (x axis, shades of colours). Gray horizontal line depicts the average RT for 1D trials across subjects and EV. Error bars represent corrected within subject SEMs [42, 43]. **e.** Hierarchical model comparison on 2D trials for the main sample showed that including Congruency (*p* < .001), yet not EV_back_ (*p* = .27), improved model fit. Including then an additional interaction of Congruency × EV_back_ improved the fit even more (*p* < .001). **f.** We replicated the behavioral results in an independent sample of 21 participants outside of the MRI scanner. Including Congruency (*p* = .009), yet not EV_back_ (*p* = .63), improved model fit. Including an additional interaction of Congruency × EV_back_ explained the data best (*p* = .017).

A baseline model including only the factor EV indicated that participants reacted faster in trials that yielded larger rewards 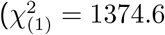, *p* < .001, Fig. 2b), in line with previous literature [44–46]. Adding Congruency to the model, we found that Congruency also affected RTs, i.e. participants reacted slower to incongruent compared to congruent trials (t-test: *t*_(34)_ = 5.38, *p*< .001, Fig. 2c, likelihood ratio test to asses improved model fit: 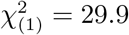, *p* < .001, Fig. 2b). Note that compared to 1D trials (Fig. 2b-c) participants were slower to respond to incongruent trials (t-test: *t*_(34)_ = −2.79, *p* = .008) and faster to respond to congruent trials (t-test: *t*_(34)_ = 2.5, *p* = .017). These effects on RT shows that even when participants accurately chose based on the relevant context, the additional information provided from the irrelevant context was not completely filtered out, affecting the speed with which choices could be made. Neither adding a main effect for EV_back_ nor the interaction of EV × EV_back_ improved model fit (LR-test with added terms: 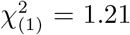, *p* = .27 and 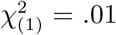, *p* = 0.9 respectively), meaning neither larger irrelevant values, nor their similarity to the objective value influenced participants’ behavior.

In a second step, we investigated if the congruency effect interacted with the expected value of the other context, i.e the points associated with the most valuable irrelevant stimulus feature (EV_back_). Indeed, we found that the higher EV_back_ was, the faster participants were on congruent trials. In incongruent trials, however, higher EV_back_ had the opposite effect (Fig. 2d, LR-test of model with added interaction: 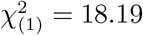, *p* < .001). In contrast, the lower valued irrelevant feature did not show comparable effects (LR-test to baseline model: 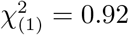, *p* = .336), and did not interact with Congruency 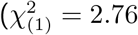, *p* = .251). This means that the expected value of a ‘counterfactual’ choice resulting from consideration of the irrelevant features mattered, i.e. that the outcome such a choice could have led to, also influenced reaction times. All major effects reported above hold when running the models nested across the levels of EV (as well as Block and Context, see Fig. S2), and replicated in an additional sample of 21 participants (15 women, *μ_age_* = 27.1, *σ_age_* = 4.91) that were tested outside of the MRI scanner (LR-tests: Congruency, 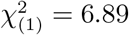, *p* = .009, EV_back_, 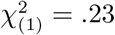, *p* = .63, Congruency × EV_back_, 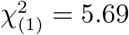, *p* = .017, Fig.2e). Details of other significant effects and alternative regression models considering for instance within-cloud or between-context value differences can be found in Fig.S3 and Fig. S4 respectively.

We took a similar hierarchical approach to model accuracy of participants in 2D trials, using mixed effects models with the same nuisance regressors as in the RT analysis. This revealed a main effect of EV (baseline model: 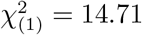, *p* < .001), indicating higher accuracy for higher EV. Introducing Congruency and then an interaction of Congruency × EV_back_ further improved model fit (LR-tests: 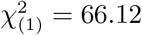, *p* < .001, 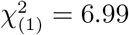, *p* = .03, respectively), reflecting decreased performance on incongruent trials, with higher error rates occurring on trials with higher EV_back_. Unlike RT, error rates were not modulated by the interaction of EV and Congruency (LR-test with EV × Congruency: 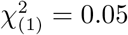, *p* = .825). Out of all nuisance regressors, only switch had an influence on accuracy (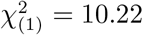, *p* = .001, in the baseline model) indicating increasing accuracy with increasing trials since the last switch trial.

In summary, these results indicated that participants did not merely perform a value-based choice among features on the currently relevant context. Rather, both reaction times and accuracy indicated that participants also retrieved the values of irrelevant features and computed the resulting counterfactual choice. We next turned to test if the neural code of the vmPFC would also incorporate such counterfactual choices, and if so, how the representation of the relevant and irrelevant contexts and their associated values might interact.

### fMRI results

#### Multivariate value and context signals co-exist within the the vmPFC

Our fMRI analyses focused on understanding the representations of expected values in vmPFC. We therefore first sought to identify a value-sensitive region of interest (ROI) that reflected expected values in 1D and 2D trials, following common procedures in the literature [e.g. 4]. Specifically, we analyzed the fMRI data using general linear models (GLMs) with separate onsets and EV parametric modulators for 1D and 2D trials (at stimulus presentation, see online methods for full model). The union of the EV modulators for 1D and 2D trials defined a functional ROI for value representations that encompassed 998 voxels, centered on the vmPFC (Fig. 3a, *p* < .0005, smoothing: 4mm, to match the multivariate analysis), which was transformed to individual subject space for further analyses (mean number of voxels: 768.14, see online methods). In the rest of the analyses we focused on the multivariate fMRI activation patterns acquired approximately 5 seconds after stimulus onset in the above-defined functional ROI.

**Figure 3:**
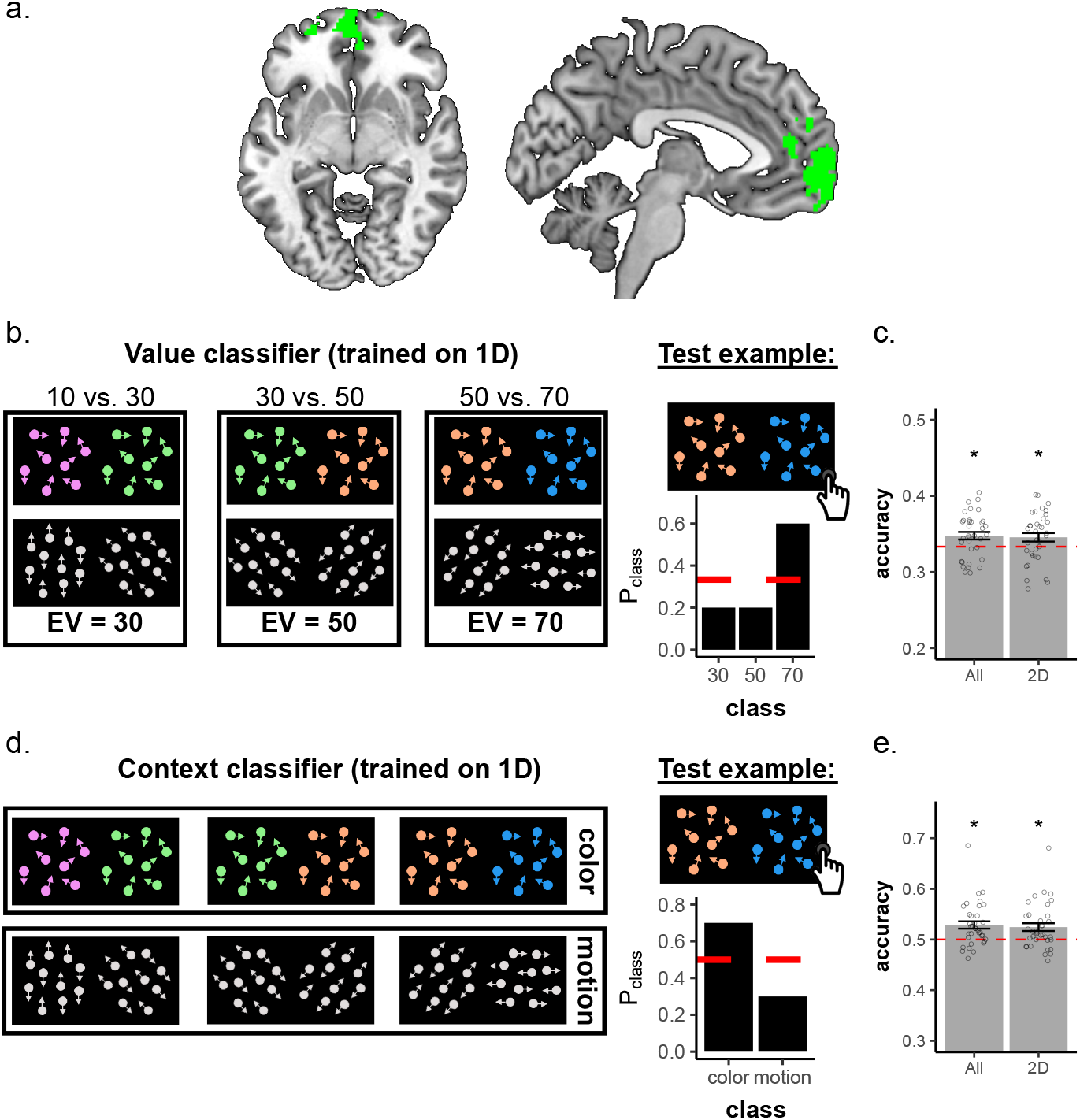
Multivariate analyses revealed expected value and context signals co-reside within the vmPFC. **a.** The union of the EV parametric modulator allowed us to isolate a cluster in the vmPFC. Displayed coordinates in the figure: x=−6, z=−6. **b.** the Value classifier was trained on behaviorally accurate 1D trials on patterns within the functionally-defined vmPFC ROI (left). This classifier was tasked with identifying the correct EV class (i.e. 30, 50 or 70). The classifier yielded for each testing example one probability for each class (right). **c.** The classifier assigned the highest probability to the correct class (objective EV) significantly above chance for all trials (*p* = .007), also when tested on generalizing to 2D trials alone (*p* = .039). Error bars represent corrected within subject SEMs [42, 43]. **d.** We trained the same classifier on the same behaviorally accurate data within the functionally-defined vmPFC ROI, only this time we split the training set to classes corresponding to the two possible contexts: Color (top) or Motion (bottom), irrespective of the EV, though we kept the training sets balanced for EV (see online methods). The classifier yielded for each testing example one probability for each class (right). **e.** The classifier assigned the highest probability to the correct class (objective Context) significantly above chance for all trials (*p* < .001), also when tested on generalizing to 2D trials alone (*p* = .002). Error bars represent corrected within subject SEMs [42, 43].

As previously mentioned, we were most interested in how the neural value representation of EV interacts with EV_back_ and its neural representation. For this purpose we trained a single multivariate multinomial logistic regression classifier to identify the EV on behaviorally accurate 1D trials, where no irrelevant values were present (henceforth: Value classifier, Fig. 3b, left; leave-one-run-out training; see online methods for details). For each testing example, the multinomial classifier assigned the probability of each class given the data (classes are the expected outcomes, i.e. ‘30’,’50’ and ‘70’, and probabilities sum up to 1, Fig. 3b, right). Crucially, it had no information about the task context of each given trial (training sets were up-sampled to balance the color/motion contexts within each set, see online methods). Because the ROI was constructed such as to contain significant information about EVs, it is not surprising that the class with the maximum probability corresponded to the objective outcome significantly more often than chance when tested on all remaining trials (*μ_all_* = .35, *σ_all_* = .029, *t*_(34)_ = 2.89, *p* = .007, Fig. 3c) as well as when tested separately to generalize from 1D to the 2D trials (*μ*_2*D*_ = .35, *σ*_2*D*_ = .033, *t*_(34)_ = 2.20, *p* = .034, Fig. 3c).

Importantly, which value expectation was relevant depended on the task context. We therefore hypothesized that, in line with previous work, vmPFC would also encode the task context, although this is not directly value-related (the average values of both contexts were identical). We thus turned to see if we can decode the trial’s context from the same ROI that was sensitive to EV. For this analysis, we trained a multinomial classifier on accurate 1D trials as before, but this time it was trained to identify if the trial was ‘Color’ or ‘Motion’ (Fig. 3d, left). Crucially, the classifier had no information as to what was the EV of each given trial, and training sets were up-sampled to balance the EVs within each set (see online methods). As expected, the classifier was above chance for decoding the correct context (*t*_(34)_ = 3.93, *p* < .001,Fig. 3e) also when tested separately to generalize to 2D trials (*t*_(34)_ = 3.2, *p* = .003, Fig. 3e). Additionally, the context is decodable also when only testing on 2D trials in which value difference in both contexts was the same (i.e. when keeping the value difference of the background 20, since the value difference of the relevant context was always 20, *t*_(34)_ = 2.73, *p* = .01).

The following analyses model directly the class probabilities estimated by the value and the context classifiers. Probabilities were modelled with beta regression mixed effects models [47]. For technical reasons, we averaged across nuisance regressors used in behavioral analyses. An exploratory analysis of raw data including nuisance variables showed that they had no influence and confirmed all model comparison results reported below (see Fig S6 and S8).

#### Multivariate neural value codes reflect value similarities and are negatively affected by contextually-irrelevant value information

We first focused on the Value classifier and asked whether EVs affected not only the probability of the corresponding class, but also influenced the full probability distribution predicted by the Value classifier. We reasoned that if the classifier is decoding the neural code of values, then similarity between the values assigned to the classes will yield similarity in probabilities associated to those classes. Specifically, we expected not only that the probability associated with the correct class be highest (e.g. ‘70’), but also that the probability associated with the closest class (e.g. ‘50’) would be higher than the probability with the least similar class (e.g. ‘30’, Fig. 4a, note that this difference also reflects which options where displayed vs not in a given trial, but see below). To test our hypothesis, we modelled the probabilities in each trial as a function of the absolute difference between the objective EV of the trial and the class (|EV-class|, i.e. in the above example with a correct class of 70, the probability for the class 50 will be modelled as condition 70-50=20 and the probability of 30 as 70-30=40). This analysis indeed revealed such a value similarity effect (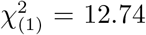, *p* < .001) also when tested separately on 1D and 2D trials (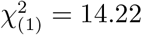, *p* < .001, 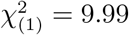, *p* = .002, respectively, Fig. 4b). Note that the difference between |EV-class| = 20 and |EV-class| = 40 also reflects which options where displayed vs. not in a given trial. Careful analysis of perceptual overlap, however, indicated that this could not explain our results (see below and SI).

**Figure 4:**
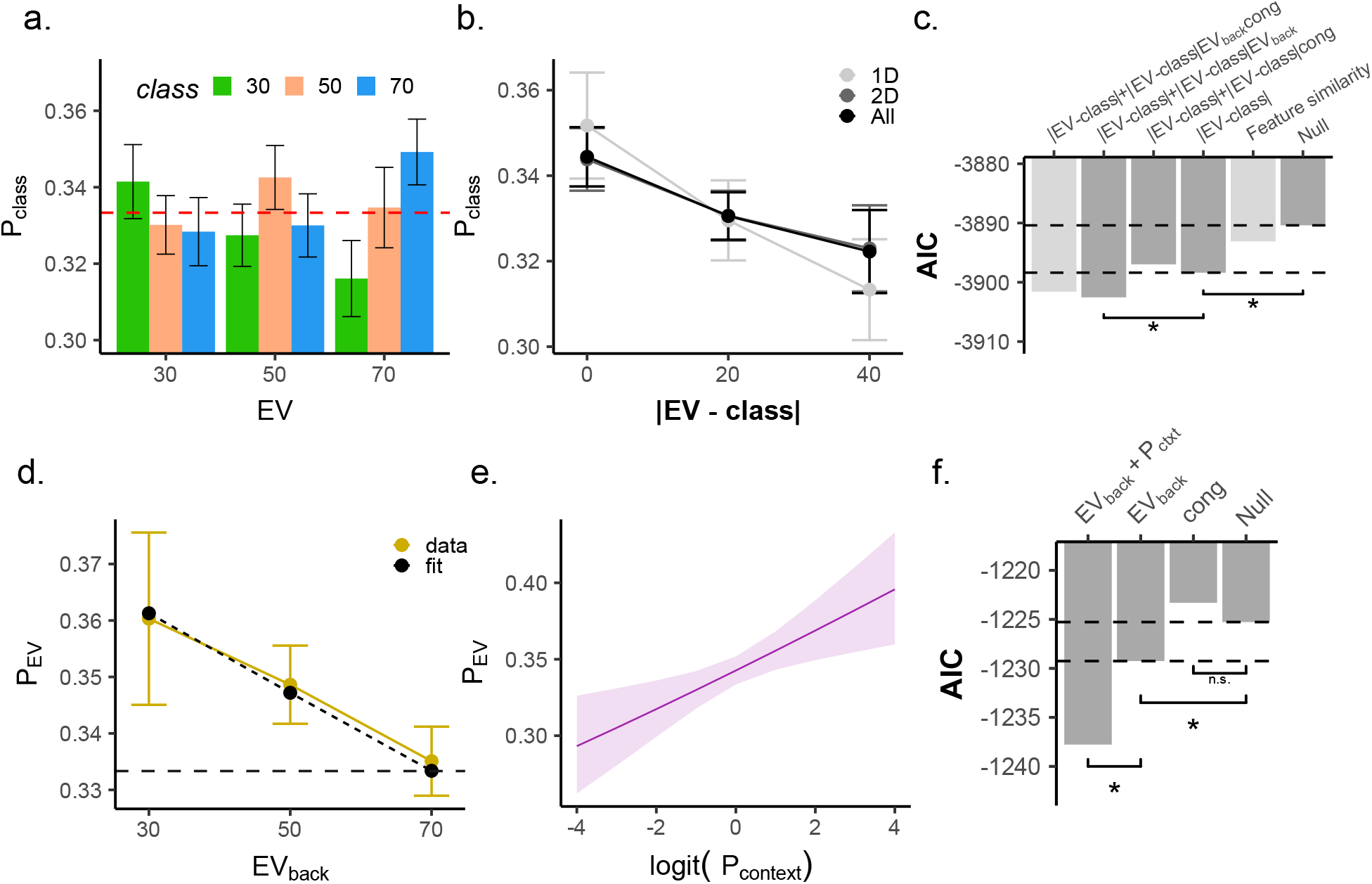
Irrelevant value expectations and neural representation of Context have independent opposite effects on EV representation in vmPFC. **a.** Analyses of all probabilities by the Value classifier revealed gradual value similarities. The y-axis represents the probability assigned to each class, colors indicate the classifier class and the x-axis represents the trial type (the objective EV of the trial). As can be seen, the highest probability was assigned to the class corresponding to the objective EV of the trial (i.e. when the color label matched the X axis label). Error bars represent corrected within subject SEMs [42, 43] **b.** Larger difference between the decoded class and the objective EV of the trial (x axis) was related to a lower probability assigned to that class (y axis) when tested in 1D, 2D or all trials (all *p* < .002, grey shades). Hence, the multivariate classifier reflected gradual value similarities. Note that when |EV – class|=0, P_class_ is the probability assigned to the objective EV of the trial (see panel d-e). Error bars represent corrected within subject SEMs [42, 43]**c.** AIC values of competing models of value probabilities classified from vmPFC. Hierarchical model comparison of 2D trials revealed not only the differences between decoded class and objective EV (|EV-class|) improved model fit (*p* < .002), but rather that EV_back_ modulated this effect (*p* = .013). Crucially, Congruency did not directly modulate the value similarity (*p* = .446). Light gray bars represent models outside the hierarchical comparison. Including a 3-way interaction (with both EV_back_ and Congruency) did not provide better AIC score (−3902.5,−3901.6, respectively). A perceptual model encoding the feature similarity between each testing trial and the training classes (irrespective of values) did not provide a better AIC score than the value similarity model (|EV-class|), see Fig S7 for details. **d.** Modeling directly only the probability assigned to the EV class (*P*_EV_). Higher EV_back_ was related to a decreased decodability of EV (*p* = .015) in behaviorally accurate trials. Yellow line reflects data, dashed line model fit from mixed effects models described in text. Error bars represent corrected within subject SEMs [42, 43]. **e.** The objective outcome was strongly represented (P_EV_), the more the context was decodable from the vmPFC (*p* = .001, x-axis, modeled as logit-transformed probability assigned to the trial-context of the trial). **f.** Hierarchical model comparisons revealed an effect of EV_back_ (*p* = .015) and no main effect of Congruency (*p* = .852). Adding an trial-context decodability effect improved prediction of the objective outcome probability, beyond the EV_back_ (*p* = .001).

Our main hypothesis was that context-irrelevant values might directly influence neural codes of expected value in the vmPFC. The experimentally manipulated background values in our task should therefore interact with the EV probabilities decoded from vmPFC. We thus asked whether the above described value similarity effect was influenced by EV_back_ and/ or Congruency in 2D trials. Analogous to our RT analyses, we used a hierarchical model comparison approach and tested if the interaction of value similarity with these factors improved model fit. We found that EV_back_, but not Congruency, modulated the value similarity effect (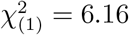, *p* = .013, 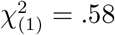, *p* = .446, respectively, Fig. 4c). This effect indicated that the higher the EV_back_ was, the less steep was the value similarity effect. These results also hold when running the models nested within the levels of EV (Fig.S6, see online methods).Additional control analyses included perceptual models that merely encoded the amount of perceptual overlap between each training class and 2D testing as well as the presence of the perceptual feature corresponding to EV_back_ in the training class. These analyses indicated that our classifier was indeed sensitive to values and not only to the perceptual features the values were associated with, see S7 for details.

#### Irrelevant values and vmPFC context signals influence expected value representations

Modelling the full probability distribution over values offers important insights, but it only indirectly sheds light on how the relevant EV representation is affected by irrelevant values in behaviorally accurate trials. We next focused on modelling the probability associated with the class corresponding to the objective EV of each 2D trial (henceforth: *P*_EV_). This also resolved the statistical issues arising from the dependency of the three classes (i.e. for each trial they sum to 1). As can be inferred by Fig 4a above, the median probability of the objective EV on 2D trials was higher than the the average of the other non-EV probabilities (*t*_(34)_ = 2.50, *p* = .017). In line with the findings reported above, we found that EV_back_ had a negative effect on *P*_EV_ (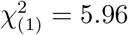, *p* = .015, Fig. 4d), meaning that higher EV_back_ trials were associated with a lower probability of the objective EV, *P*_EV_. This confirms that EV_back_ specifically decreases the decodability of the objective EV.

Next we hypothesized that if vmPFC is involved in signaling the trial context as well as the values, then the strength of context signal might relate to the strength of the contextually relevant value. We found that P_*context*_ had a positive effect on the decodability of EV and that adding this term in addition to EV_back_ to the P_EV_ model improved model fit (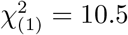, *p* = .001, Fig. 5e). In other words, the better we could decode the context, the higher was the probability assigned to the correct EV class. The effect of EV_back_ also holds when running the model nested inside the levels of EV (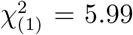, *p* = 0.014, Fig.S8b), and cannot be attributed to perceptual effects, since replacing EV_back_ with a regressor indicating the presence of its corresponding perceptual feature did not provide a better model fit (AICs: −1229.2,−1223.3, respectively). We found no evidence for an interaction of EV_back_ and P_*context*_ (LR-test with added term: 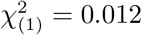, *p* = .91).

**Figure 5:**
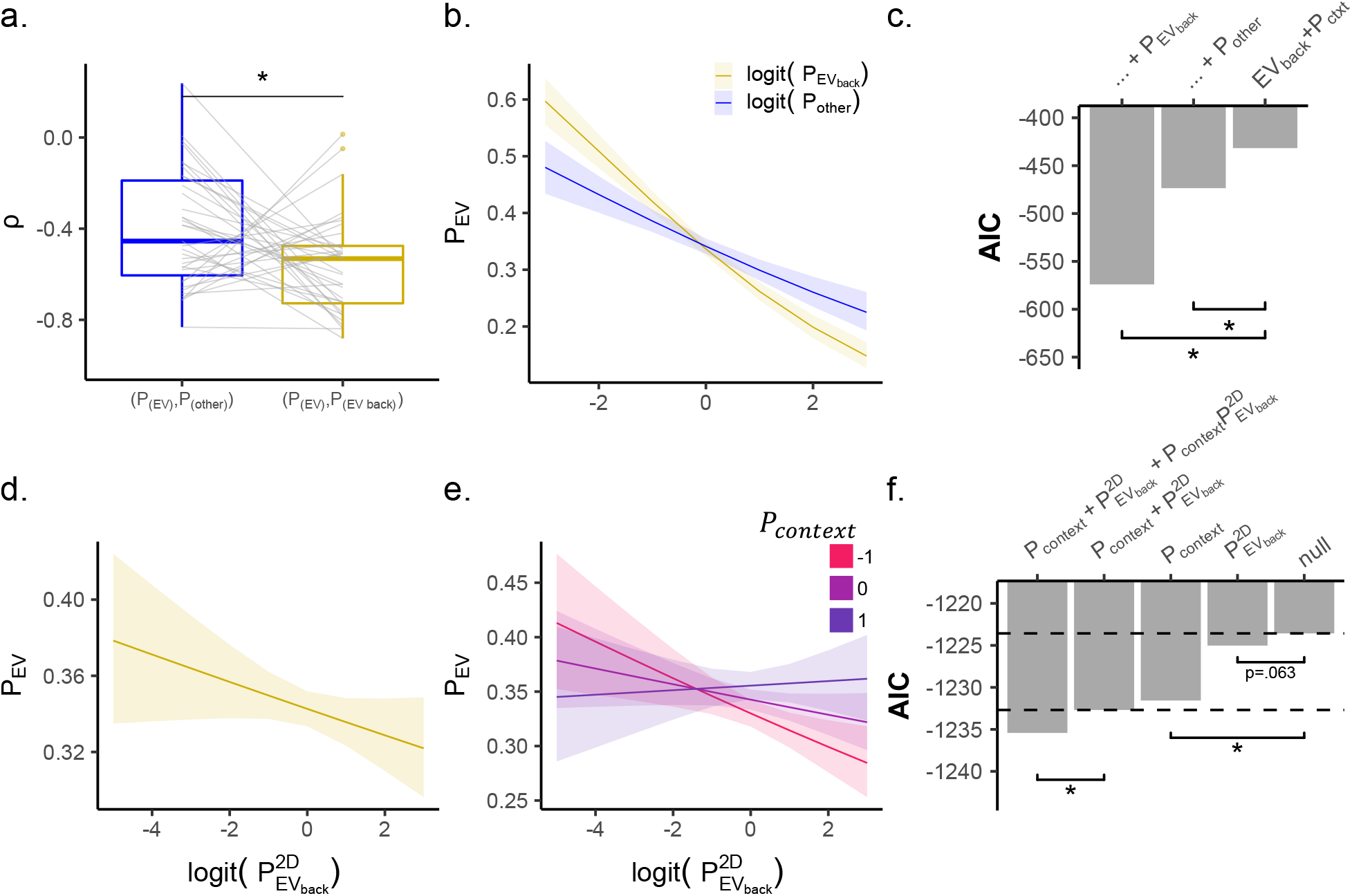
Neural representation of EV_back_ in vmPFC directly influence EV representation and its relation to the context signal, in behaviorally accurate trials. **a.** Testing trials in which EV≠EV_back_ revealed that both the probability the value classifier assigned to the class corresponding to EV_back_ (P_EV_back__, yellow line) as well as to the third other class (P_Other_, blue line) had a strong negative correlation with the probability assigned to P_EV_. However, the correlation of P_EV_ and P_EV_back__ (yellow) was stronger than with P_Other_ (blue, *p* = .017). **b.** In trials where EV≠EV_back_, the effect of P_EV_ was stronger than P_Other_ (x-axis, modeled as multinomial-logit-transformed probability assigned to the trial-context of the trial, see online methods for details). **c.** Hierarchical model comparisons revealed that adding an effect of either P_EV_back__ or P_Other_ increased model fit. However, adding P_EV_back__ provided a better (i.e. lower) AIC score (−574, −473, respectively). **d.** Stronger representation of the irrelevant EV (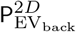, x-axis, from a classifier trained on 2D trial to directly detect EV_back_, modeled as multinomial-logit-transformed probability) slightly decreased the representation of the objective outcome (P_EV_, y-axis, nested in the levels of EV_back_, *p* = .063, yet see AIC scores in panel f.). Plotted are model predictions. **e.** The higher P_*context*_ was, the weaker was the effect of 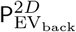 on P_EV_. In other words, stronger representation of the Context weakened the effect the representation of EV_back_ had on EV representation (*p* = .022). Plotted are model predictions. **f.** Comparing models of P_EV_ revealed that adding either P_*context*_ or 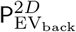 (nested within the levels of EV_back_) improved AIC scores (however only the former was significant according to LR test: *p* = .002 and *p* = .063 respectively). Adding an interaction term of 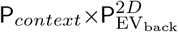 improved model fit compared to only P_*context*_ (*p* = .022) and also to a model with 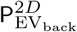 in it as well (*p* = .029). Note that by P_EV_back__ (panels a-b) we indicate the probabilities the Value classifier, trained on 1D trials, assigned the EV_back_ class, whereas 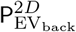 indicated probabilities from a classifier tasked to identify EV_back_ directly (trained on 2D trials)

Interestingly, and unlike in the behavioral models, we found that neither Congruency nor its interaction with EV or with EV_back_ influenced *P*_EV_ (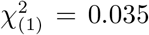, *p* = .852, 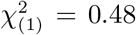, *p* = .787, 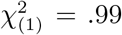, *p* = .317, respectively, Fig. 5f). Additionally, when value expectations of both contexts matched (i.e. when EV=EV_back_) there was neither an increase nor a decrease of *P*_EV_ (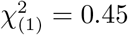, *p* = .502, see online methods for details). Lastly, as in our behavioral analysis, we evaluated alternative models of *P*_EV_ that included a factor reflecting within-option or between-context value differences, or alternatives for EV_back_ (Fig.S8).

In summary, this indicates that the neural code of value in the vmPFC is affected by contextually-irrelevant value expectations, such that larger alternative values disturb neural value codes in vmPFC more than smaller ones. Even though the neural code in vmPFC is mainly influenced by the contextually relevant EV, the representation of the relevant expected value was measurably weakened on trials in which the alternative context would lead to a large expected value. This was the case even though the alternative value expectations were not relevant in the context of the considered trials. The effect occurred irrespective of the agreement or action-conflict between the relevant and irrelevant values (unlike participants’ behaviour). Lastly, we found that the Context is represented within the same region as the EV, and that the strength of its representation is directly linked to the representation of EV. Our finding therefore suggests that the (counterfactual) value of irrelevant features must have been computed and poses the power to influence neural codes of objective EV in vmPFC.

#### Representational conflict between EV and EV_back_ moderated by the Context signal

Our previous analyses indicated that the probability to correctly decode EV from vmPFC activity decreased with increasing EV_back_. This decrease could reflect a general disturbance of the value retrieval process caused by the distraction of competing values. Alternatively, the encoding of EV_back_ could directly compete with the representation of EV – reflecting that the irrelevant values might be represented using similar neural codes used for the objective EV (note that the classifier was trained in the absence of task-irrelevant values, i.e. the objective EV of 1D trials). In order to test this idea, we took the Value classifier (Fig. 3b.) and tested it on trials in which EV ≠ EV_back_, i.e. in which the value expected in the current task context was different from the value that would be expected, would the same trial occur in a different task-context. This allowed us to interpret the class probabilities of our Value classifier as either signifying EV (P_EV_), EV_back_ ( P_EV_back__) or a value that was expected in neither case (P_other_). We then examined the correlation between each pair of classes. To prevent a bias between the classes, we only included trials in which a feature that signified the ‘other’ value appeared on the screen as either a relevant or irrelevant feature.

For each trial, the three class probabilities sum up to 1 and hence are strongly biased to correlate negatively with each other. Not surprisingly, we found such strong negative correlations across participants of both pairs of probabilities, i.e. between P_EV_ and P_EV_back__ (*ρ* = −.56, *σ* = .22) as well as between P_EV_ and P_other_ (*ρ* = −.40, *σ* = .25). However, the former correlation was significantly more negative than the latter (*t*_(34)_ = −2.77, *p* = .017, Fig. 5a), indicating that when the probability assigned to the EV decreased, it was accompanied by a stronger increase in the probability assigned to EV_back_, akin to a competition between both types of expectations. We tested this formally by adding either P_EV_back__ or P_other_ to the model predicting P_EV_ (as multinomial-logit-transformed probability, see online methods). We found that the model including P_EV_back__ resulted in a better (i.e. smaller) AIC (−574), compared to the model with P_other_ as predictor (−473, 5c).

Next, we tested whether vmPFC represents EV_back_ directly by training classifiers for each class of EV_back_ on accurate 2D trials. A balanced accuracy did not surpass chance level (*t*_(34)_ = 0.96, *p* = .171). However, we believe that the reason for that relates to the fact that the number of unique examples for each class of EV_back_ differed drastically (due to our design, see Fig. 1c), and our approach of combining one-vs-rest training with oversampling and sample weights could not fully counteract these imbalances (see online methods). We therefore proceeded to ask if the probability the EV_back_ classifier assigned to the correct class 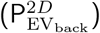 might still relate to encoding of the relevant value as indicated by the Value classifier (i.e., P_EV_). Importantly, both classifiers were trained on independent data (EV_back_ classifier was trained in 2D, the Value classifier on 1D trial), but in both cases on behaviorally accurate trials, i.e. trials where participants choose according to EV, as indicated by the relevant context. A mixed effect model of P_EV_ with random effects nested within levels of EV_back_ confirmed our previous finding that the strength of context encoding affected value encoding (effect of P_context_, LR-test: 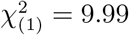, *p* = .002). Notably, we also found that encoding of EV_back_ when measured independently 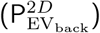 improved the AIC score of the model (−1223.6 to −1225.0, but note that in the LR test 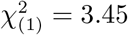, *p* = 0.063, Fig. 5d)). This confirms our previous analysis showing that stronger neural representation of EV_back_ reduced EV decodability. Most remarkably, the effect of Context, P_*context*_, interacted with the effect of expected value of the background, i.e. 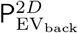 (LR test: 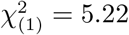, *p* = 0.022, Fig. 5e). In other words, the stronger the contribution of Context to EV representation, the weaker the influence EV_back_ representation had on EV.

In summary, we showed the neural representation of EV was reduced in trials with higher expected value of the background, and weakened EV representations indeed were accompanied by stronger neural representations of such background values in the same vmPFC region on a trial by trial basis. We confirmed this by showing the same relationship in two independent analyses that probed the neural representation of EV_back_ either through the standard Value classifier or a separate classifier trained on different trials and tested nested in the levels of EV_back_. Most strikingly, the negative influence of EV_back_ representation on EV decodability was governed by the Context signal, i.e. when the link between the Context and EV was strongest, the EV_back_ representation was effect diminished. As will be discussed later in detail, we consider this to be evidence for parallel processing of two task aspects in this region, EV and EV_back_, which are governed by the Context signal.

#### Neural representation of EV, EV_back_ and Context guide choice behavior

To conclude the multivariate analysis, we investigated how vmPFC’s representations of EV, EV_back_ and the relevant Context influence participants’ behavior. We first investigated this influence on choice accuracy. Importantly, the two contexts only indicate different choices in incongruent trials, where a wrong choice could be a result of a strong influence of the irrelevant context. Motivated by our behavioral analyses that indicated an influence of the irrelevant context on accuracy, we asked whether P_EV_back__ was different on behaviorally wrong or incongruent trials. We found an interaction of accuracy × Congruency (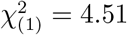, *p* = .034, Fig. 6a) that indicated increases in P_EV_back__ in accurate congruent trials and decreases in wrong incongruent trials. Hence, on trials in which participants erroneously chose the option with higher valued irrelevant features, P_EV_back__ was increased. Focusing only on behaviorally accurate trials, we found no effect of EV nor Congruency on P_EV_back__ (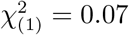, *p* = .794, 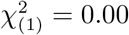, *p* = .987 respectively).

**Figure 6:**
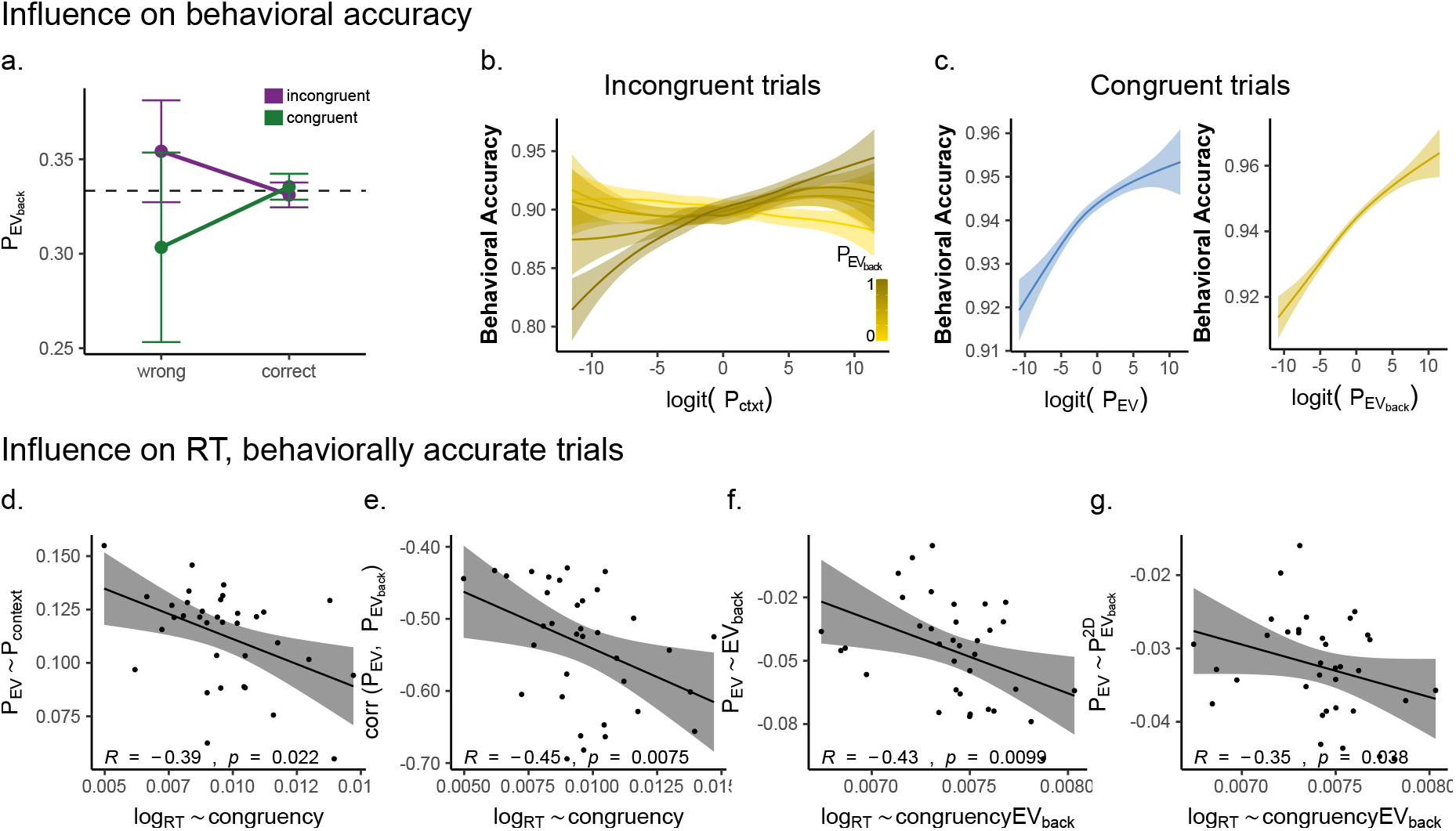
Neural representations of context and value in vmPFC jointly guide behavior. **a.** Representation of EV_back_ relates to behavioral accuracy. The probability the Value classifier assigned to EV_back_ (P_EV_back__, y-axis) was increased when participants chose the option based on EV_back_. Specifically, in incongruent trials (purple), high P_EV_back__ was associated a wrong choice, whereas in Congruent trials (green) it was associated with correct choices. This effect is preserved when modeling only wrong trials (main effect of Congruency: 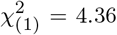, *p* = .037). Error bars represent corrected within subject SEMs [42, 43]. **b.** Lower context decodability of the relevant context (Context classifier, x axis) was associated with less behavioral accuracy (y-axis) in incongruent trials (*p* = .051). This effect was modulated by the representation of EV_back_ in vmPFC (*p* = .012, shades of gold, Value classifier), i.e. it was stronger in trials where EV_back_ was strongly decoded from the vmPFC (shades of gold, plotted in 5 quantiles). Shown are fitted slopes from analysis models reported in the text. **c.** Decodability from the Value classifier of both EV (*p* = .058, blue, left) and EV_back_ (*p* = .009, gold, right) labels had a positive relation to behavioral accuracy (y axis) in congruent trials. Shown are fitted slopes from analysis models reported in the text. **d.** When focusing on behaviorally accurate trials, participants that had a weaker effect of Context decodability on EV decodability (y-axis, Fig 4e), had a stronger effect of Congruency on RT (x-axis, larger values indicate a stronger RT decrease in incongruent compared to congruent trials, equivalent to the distance between the purple and green lines in Fig.2b.) **e.** When focusing on behaviorally accurate trials, participants who had a stronger effect of EV_back_ on the EV decodability (y-axis, more negative values indicate stronger decrease of *P*_EV_ as a result of high EV_back_, see Fig.4d) also had a stronger effect of Congruency on their RT (x-axis, same as panel d.). **f.** When focusing on behaviorally accurate trials, participants that had a stronger (negative) correlation of P_EV_ and P_EV_back__ (y-axis, more negative values indicate stronger negative relationship, see Fig.5a.) also had a stronger modulation of EV_back_ on the effect of Congruency on their RT (x-axis, more positive values indicate stronger influence on the slow incongruent and fast congruent trials. Equivalent to the distance between the purple and green lines in Fig 2d). **g.** When focusing on behaviorally accurate trials, participants that had a stronger (negative) effect of 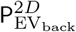 on P_EV_ (y-axis, more negative values indicate stronger decrease of *P*_EV_ as a result of 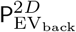, see Fig.5d), also had a stronger modulation of EV_back_ on the effect of Congruency on their RT (x-axis, same as panel f.).

Motivated by the different predictions for congruent and incongruent trials, we next turned to model these trial-types separately. When focusing on incongruent trials (Fig. 6b) we found that a weaker representation of the relevant context was marginally associated with an increased error rate (negative effect of P_*context*_) on accuracy, LR-test with P_*context*_): 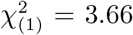, *p* = .055). Moreover, if stronger representation of the wrong context (i.e. 1-P_*context*_)) decreases accuracy, than stronger representation of the value associated with this context (EV_back_) should strengthen that influence. Indeed, we found that adding a P_*context*_ × P_EV_back__ term to the model explaining error rates improved model fit (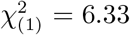, *p* = .012, Fig. 6b). However, neither the representation of EV nor EV_back_ directly influence behavioral accuracy (P_EV_: 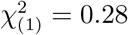, *p* = .599, P_EV_back__: 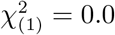, *p* = .957). Contrary to incongruent trials, in congruent trials choosing the wrong choice is unlikely a result of wrong context encoding, since both contexts lead to the same choice. Indeed, when focusing on Congruent trials (Fig. 6c) there was no influence of P_*context*_) on accuracy (LR-test: 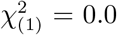, *p* = .922). However, strong representation of either relevant or irrelevant EV should lead to a correct choice. Indeed, we found that both an increase in P_EV_back__ and (marginally) in P_EV_ had a positive relation to behavioral accuracy (P_EV_back__: 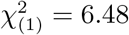, *p* = .011, P_EV_: 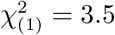, *p* = .061, Fig. 6c).

Finally, if the EV representation in vmPFC does guide behavior, then any influence on it should not be restricted to choice-accuracy and should extend to RT of behaviorally accurate trials, i.e. trials in which participants choose according to the relevant context. In line with this idea, we found that participants who had a weaker influence of the Context representation on the EV representation, had a stronger Congruency effect on their RT (*r* = −.39, *p* = .022 Fig 6d). In other words, the less influence the Context signal had on enhancing the relevant EV signal, the bigger was the influence the value of a counterfactual choice had on participants’ RTs. Next, we hypothesized that if vmPFC represents both EV and EV_back_ simultaneously, than increasing conflict between the representations of the two should directly influence participant’s RT. Strikingly, we found that all three main findings of conflict between EV and EV_back_ correlated with the Congruency-related RT effect: Participants who showed more negative correlation between P_EV_ and P_EV_back__ (taken from the 1D trained value classifier) had a stronger Congruency effect on their RTs (*r* = −.45, *p* = .008, Fig. 6e); Participants who had a stronger negative effect of EV_back_ on EV representation, had a stronger modulation of EV_back_ on the RT Congruency effect (*r* = .43, *p* = .01, Fig. 6f); Finally, the same was true when considering the strength of the effect of the neural representation of EV_back_ 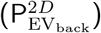 on the neural EV signal in relation to the above behavioral marker (*r* = 35, *p* = .004, Fig. 6g). In other words, we saw that both high valued EV_back_ and stronger EV_back_ representation were related to the behavioral modulation effects EV_back_ had on Congruency (i.e. stronger influence on the slow incongruent and fast congruent trials).

In summary, behavioral accuracy seemed to be influenced by context representation and its associated EV only in incongruent trials (i.e. when it mattered), whereas both neural representation of EV and EV_back_, but not the context, contributed to choice-accuracy in congruent trials. When focusing on accurate trials only, participants who exhibited a larger association between the decodability of EV and of Context, had a smaller influence of the counterfactual choice on their behavior. Lastly, an increase in any effect of conflict between the representations of EV and EV_back_ directly resulted in an increase of the RT effect of conflict between the two EVs. Brought together these findings show that the representations of EV, EV_back_ and Context in the vmPFC don’t only interact with each other, but directly guide choice behavior as reflected in accuracy as well as RT in behaviorally accurate trials

#### No evidence for univariate modulation of contextually irrelevant information on expected value signals in vmPFC

The above analyses indicated that multiple value expectations are represented in parallel within vmPFC. Lastly, we asked whether whole-brain univariate analyses could also uncover evidence for processing of multiple value representations. Detailed description of the univariate analysis can be found in Fig. S9. Notably, unlike the multivariate analysis, no univaraite modulation effect of neither Congruency, EV_back_ nor their interaction was observed in any frontal region (but a negative effect of EV_back_ in the Superior Temporal Gyrus, *p* < .001, Fig. S9c). We also found no region for the univariate effect of Congruency × EV_2D_ interaction (even at *p* < .005). However, we found a negative univariate effect of Congruency × EV_back_ in the primary motor cortex at a liberal threshold, which indicated that the difference between Incongruent and Congruent trials increased with higher EV_back_, akin to a response conflict (*p* < .005, Fig. S9d). These findings contrast with the idea that competing values would have been integrated into a single EV representation in the vmPFC, because this account would have predicted a higher signal for Congruent compared to Incongruent trials.

## Discussion

In this study, we investigated how contextually-irrelevant value expectations influence behavior and neural activation patterns in vmPFC. We asked participants to make choices between options that had different expected values in different task-contexts. Participants reacted slower when the expected values in the irrelevant context favored a different choice, compared to trials in which relevant and irrelevant contexts favored the same choice. This Congruency effect increased with increasing reward associated with the hypothetical choice in the irrelevant context (EV_back_). We then identified a functional ROI that is univariately sensitive to the objective, i.e. relevant, expected values (EV).

We first showed that both EV and the Context could be decoded from vmPFC activity in behaviorally accurate 2D trials, i.e. trials where participants choose according to the highest value of the relevant context. Multivariate analysis then focused on the probability distribution of different values in vmPFC and found that higher EV_back_ was associated with a degraded representation of the objective EV (P_EV_). This decrease in decodability of the value in the relevant context was associated with an increase in the value that would be obtained in the other task-context (P_EV_back__), akin to a conflict of the two value representations. Although we could not find clear group-level evidence for direct EV_back_ decoding, we show that fluctuations in the decodability of the EV_back_ across trials 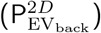 were related to a reduced EV representation in the same vmPFC ROI. Importantly, increased representation of context (P_*context*_) was associated with increase in value retrieval, but also mediated the relationship between the two EVs. Specifically, when the Context signal was strong, the negative effect of 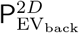 on EV was diminished. We also found that the above-mentioned multifaceted value and context representations in vmPFC were linked to participants choice accuracy as well as RT of accurate trials. Increased representation of EV_back_ in vmPFC during stimuli presentation was associated with an increased chance of choosing accordingly, irrespective of its agreement with the relevant context. Moreover, when the irrelevant context pointed to the wrong choice in incongruent trials, stronger vmPFC representation of the alternative (wrong) context and its corresponding value were related to higher error rates. However, when both contexts agreed on the action to be made, stronger representation of either of their EVs were strongly related to making a correct choice. Even when only looking in behaviorally accurate trials, the impact of EV_back_, and its neural representation, on relevant value representations was associated with how strongly RTs were influenced by the value of counterfactual choices (note that the neural effects occurred irrespective of choice congruency). Lastly, the link between Context and EV signals was also related to choice congruency RT effects. These data suggest that information within the vmPFC is organized into a complex multi-faceted representation, in which multiple values of the same choice under different task-contexts are co-represented and compete in guiding behavior, while the Context signal might act as a moderator of this so-called competition.

Behavioral analyses showed that hypothetical, context-irrelevant, values can still influence choice behavior. In our experiment the relevant features were cued explicitly and the rewards were never influenced by the irrelevant features. Nevertheless, participants’ reactions were influenced not only by the contextually relevant outcome, but also by the (irrelevant) values a counterfactual choice in a different context would yield. These results raise the question how internal value expectation(s) of the choice are shaped by the possible contexts. One hypothesis could be that rewards expected in both contexts integrate into a single EV for a choice, which in turn guides behavior. This perspective suggests that the expected value of choices that are associated with high rewards in both contexts will increase, resulting in an increase in vmPFC signal. An alternative hypothesis would be that both values are kept separate, and will be processed in parallel. In this case, EV representations in vmPFC would not be expected to increase for choices valuable in both contexts. Rather, the specific EV_back_ should be represented in addition to the EV, and possibly compete with it. Moreover, how strongly the two competing value representations influence choices would then depend on the representational strength of the context, while conflicts between incongruent motor commands might be resolved outside of vmPFC.

To differentiate these possibilities, we focused our analysis on the vmPFC, where we could distinguish between a single integrated value and simultaneously co-occurring representations. Notably, the representation of the current task context, which might influence the interaction of values, is known to be represented in the same region and the overlapping orbitofrontal cortex [e.g., 30, 32, 33, 48]. It therefore seemed to be a good candidate region to help illuminate how values stemming from different contexts, as well as information about the contexts themselves, might interact in the brain.

Contradictory to the integration hypothesis, we found no effect of EV_back_ on univariate vmPFC signals. We also did not find any Congruency effect in vmPFC, eliminating a congruency-dependent integration. The latter would predict an increased signal for congruent compared to incongruent trials. Even when the relevant and irrelevant expected values were the same (EV = EV_back_), classifier evidence for EV did not increase. This suggests some differences in the underlying representations of relevant and irrelevant values. At the same time, our analysis showed that the value classifier *was* sensitive to the expected value of the irrelevant context in 2D trials, even though it was trained on 1D trials during which irrelevant values were not present. This could suggest that within the vmPFC ‘conventional’ expected values and counterfactual values are encoded using partially, but not completely, similar patterns.

This interpretation would also be supported by our findings that the negative effect EV_back_ had on EV representations could be reconciled with participants’ behavior, where a large or stronger EV_back_ either impaired or improved performance, depending on congruency. In the first case, when choices for the two contexts differ, competing EV and EV_back_ led to performance decrements; in the second case, when choices are the same, both of the independently contributing representations supported the same reaction and therefore benefited performance. Crucially, even in trials where participants choose accurately by the relevant context, we found the same relationship, namely that participants that had a stronger influence of EV_back_ and its representation on EV signals, also had an increase in the congruency RT effect. This shows that even in those trials the counterfactual choice was still present within the vmPFC and influenced RTs. Our results therefore are in line with the interpretation that both relevant and irrelevant values are retrieved, represented in parallel within the vmPFC and influence behavior.

Univariate analyses revealed a weak negative modulation of primary motor cortex activity by Congruency. Akin to a response conflict, this corresponds to recent findings that distracting information can be traced to areas involved in task execution cortex in humans and monkeys [24, 25]. Crucially however, unlike in previous studies, the modulation found in our study was dependent on the specific expected value of the alternative context. This could suggest that conflicts between incongruent actions based on parallel value representations in the vmPFC are resolved in motor cortex. This would also be in line with our interpretation that the vmPFC does not integrate both tasks into a single EV representation that drives choice.

One important implication of our study concerns the nature of neural representations in the vmPFC/mOFC. A pure perceptual representation should be equally influenced by all four features on the screen. Yet, our decoding results could not have been driven by the perceptual properties of the chosen feature, and effects of background values could also not be explained by perceptual features of the ignored context (Fig. 3 and Fig. S7). Rather, we find that in addition to (expected) values, vmPFC/mOFC represents task-states, which help to identify relevant information if information is partially observable, as suggested by previous work [30, 48]. Note that the task context, which we decode from vmPFC activity in the present paper, could be considered as a superset of the more fine grained task states that reflect the individual motion directions/colors involved in a comparison. Any area sensitive to these states would therefore also show decoding of context as defined here. These findings are in line with work that has found that EV could be one additional aspect of OFC activity [39], which is multiplexed with other task-related information. Crucially, the idea of task-state as integration of task-relevant information [35, 49] could explain why this region was found crucial for integrating valued features, when all features of an object are relevant for choice [18, 35], although some work suggests that it might even reflect integration of features not carrying any value [36].

To conclude, the main contribution of our study is that we elucidated the relation between task-context and value representations within the vmPFC. By introducing multiple possible values of the same option in different contexts, we were able to reveal a complex representation of task structure in vmPFC, with both task-contexts and their associated expected values activated in parallel. The decodability of both contexts and value(s) independently from vmPFC, and their relation to choice behavior, hints at integrated computation of these in this region. We believe that this bridges between findings of EV representation in this region to the functional role of this region as representing task-states, whereby relevant and counterfactual values can be considered as part of a more encompassing state representation.

## Acknowledgments

NWS was funded by an Independent Max Planck Research Group grant awarded by the Max Planck Society (M.TN.A.BILD0004) and a Starting Grant from the European Union (ERC-2019-StG REPLAY-852669). NM was funded by and is grateful for a scholarship from the Ernst Ludwig Ehrlich Studienwerk (ELES) and Einstein Center for Neuroscience (ECN) Berlin throughout this study. We thank Angela J. Langdon for comments on the manuscript. We thank Gregor Caregnato for help with participant recruitment, Anika Löwe, Lena Maria Krippner, Sonali Beckmann and Nadine Taube for help with data acquisition, all participants for their participation and the Neurocode lab for numerous contributions and help throughout this project.

## Data availability statement

Behavioral and MRI data needed to replicate the findings of this study will be made available upon publication.

## Code availability statement

Custom code for all analyses conducted in this study will be made available upon publication.

## Online Methods

### Participants

Forty right-handed young adults took part in the experiment (18 women, *μ_age_* = 27.6, *σ_age_* = 3.35) in exchange for monetary reimbursement. Participants were recruited using the participant database of Max-Planck-Institute for Human Development. Beyond common MRI-safety related exclusion criteria (e.g. piercings, pregnancy, large or circular tattoos etc.), we also did not admit participants to the study if they reported any history of neurological disorders, tendency for back pain, color perception deficiencies or if they had a head circumference larger than 58 cm (due to the limited size of the 32-channel head-coil). After data acquisition, we excluded five participants from the analysis; one for severe signal drop in the OFC, i.e. more than 15% less voxels in functional data compared to the OFC mask extracted from freesurfer parcellation of the T1 image [50, 51]. One participant was excluded due to excessive motion during fMRI scanning (more than 2mm in any axial direction) and three participants for low performance (less than 75% accuracy in one context in the main task). In the behavioral-replication, 23 young adults took part (15 women, *μ_age_* = 27.1, *σ_age_* = 4.91) and two were excluded for the same accuracy threshold. Due to technical reasons, 3 trials (4 in the replication sample) were excluded since answers were recorded before stimulus was presented and 2 trials (non in the replication) in which RT was faster than 3 SD from the mean (likely premature response). The monetary reimbursement consisted of a base payment of 10 Euro per hour (8.5 for replication sample) plus a performance dependent bonus of 5 Euro on average. The study was approved the the ethics board of the Free University Berlin (Ref. Number: 218/2018).

### Experimental procedures

#### Design

Participants performed a random dot-motion paradigm in two phases, separated by a short break (minimum 15 minutes). In the first phase, psychophysical properties of four colors and four motion directions were first titrated using a *staircasing task*. Then, participants learned the rewards associated with each of these eight features during a *outcome learning task*. The second phase took place in the MRI scanner and consisted mainly of the *main task*, in which participants were asked to make decisions between two random dot kinematograms, each of which had one color and/or one direction from the same set. Note there were two additional mini-blocks of 1D trials only, at the end of first- and at the start of the second phase (during anatomical scan, see below). The replication sample completed the same procedure with the same break length, but without MRI scanning. That is, both phases were completed in a behavioral testing room. Details of each task and the stimuli are described below. Behavioral data was recorded during all experiment phases. MRI data was recorded during phase 2. We additionally collected eye-tracking data (EyeLink 1000; SR Research Ltd.; Ottawa, Canada) both during the staircasing and the main decision making task to ensure continued fixation (data not presented). The overall experiment lasted between 3.5 and 4 hours (including the break between the phases). Additional information about the pre-scanning phase can be found in Fig. S1.

#### Room, Luminance and Apparatus

Behavioral sessions were conducted in a dimly lit room without natural light sources, such that light fluctuations could not influence the perception of the features. A small lamp was stationed in the corner of the room, positioned so it would not cast shadows on the screen. The lamp had a light bulb with 100% color rendering index, i.e. avoiding any influence on color perception. Participants sat on a height adjustable chair at a distance of 60 cm from a 52 cm horizontally wide, Dell monitor (resolution: 1920 x 1200, refresh rate 1/60 frames per second). Distance from the monitor was fixed using a chin-rest with a head-bar. Stimuli were presented using psychtoolbox version 3.0.11 [52–54] in MATLAB R2017b [55]In the MRI-scanner room lights were switched off and light sources in the operating room were covered in order to prevent interference with color perception or shadows cast on the screen. Participants lay inside the scanner at distance of 91 cm from a 27 cm horizontally wide screen on which the task was presented a D-ILA JVC projector (D-ILa Projektor SXGA, resolution: 1024×768, refresh rate: 1/60 frames per second). Stimuli were presented using psychtoolbox version 3.0.11 [52–54] in MATLAB R2012b [56] on a Dell precision T3500 computer running windows XP version 2002.

#### Stimuli

Each cloud of dots was presented on the screen in a circular array with 7°visual angle in diameter. In all trials involving two clouds, the clouds appeared with 4°visual angle distance between them, including a fixation circle (2°diameter) in the middle, resulting in a total of 18°field of view [following total apparatus size from 41]. Each cloud consisted of 48 square dots of 3×3 pixels. We used four specific motion and four specific color features.

To prevent any bias resulting from the correspondence between response side and dot motion, each of the four motion features was constructed of two angular directions rotated by 180°, such that motion features reflected an axis of motion, rather than a direction. Specifically, we used the four combinations: 0°-180° (left-right), 45°-225° (bottom right to upper left), 90°-270° (up-down) and 135°-315° (bottom left – upper right). We used a Brownian motion algorithm [e.g. 41], meaning in each frame a different set of given amount of coherent dots was chosen to move coherently in the designated directions in a fixed speed, while the remaining dots moved in a random direction (Fig. S1). Dots speed was set to 5° per second [i.e. 2/3 of the aperture diameter per second, following 41]. Dots lifetime was not limited. When a dot reached the end of the aperture space, it was sent ‘back to start’, i.e. back to the other end of the aperture. Crucially, the number of coherent dots (henceforth: motion-coherence) was adjusted for each participant throughout the staircasing procedure, starting at 0.7 to ensure high accuracy [see 41]. An additional type of motion-direction was ‘random-motion’ and was used in 1D color clouds. In these clouds, dots were split to 4 groups of 12, each assigned with one of the four motion features and their adjusted-coherence level, resulting in a balanced subject-specific representation of random motion.

In order to keep the luminance fixed, all colors presented in the experiment were taken from the YCbCr color space with a fixed luminance of Y = 0.5. YCbCr is believed to represent human perception in a relatively accurate manner [cf. 57]. In order to generate an adjustable parameter for the purpose of staircasing, we simulated a squared slice of the space for Y = 0.5 (Fig. S1) in which the representation of the dots color moved using a Brownian motion algorithm as well. Specifically, all dots started close to the (gray) middle of the color space, in each frame a different set of 30% of dots was chosen to move coherently towards the target color in a certain speed whereas all the rest were assigned with a random direction. Perceptually, this resulted in all the dots being gray at the start of the trial and slowly taking on the designated color. Starting point for each color was chosen based on pilot studies and was set to a distance of 0.03-0.05 units in color space from the middle. Initial speed in color space (henceforth: color-speed) was set so the dots arrive to their target (23.75% the distance to the corner from the center) by the end of the stimulus presentation (1.6s). i.e. distance to target divided by the number of frames per trial duration. Color-speed was adjusted throughout the staircasing procedure. An additional type of color was ‘no color’ for motion 1D trials for which we used the gray middle of the color space.

#### Staircasing task

In order to ensure RTs mainly depended on associated values and not on other stimulus properties (e.g. salience), we created a staircasing procedure that was conducted *prior to value learning*. In this procedure, motion-coherence and color-speed were adjusted for each participant in order to minimize between-feature detection time differences. As can be seen in Fig. S1, in this perceptual detection task participants were cued (0.5s) with either a small arrow (length 2°) or a small colored circle (0.5°diameter) to indicate which motion-direction or color they should choose in the upcoming decision. After a short gray (middle of YCbCr) fixation circle (1.5s, diameter 0.5°), participants made a decision between the two clouds (1.6s). Clouds in this part could be either both single-feature or both dual-features. In dual feature trials, each stimulus had one color and one motion feature, but the cue indicated either a specific motion or a specific color. After a choice, participants received feedback (0.4s) whether they were (a) correct and faster than 1 second, (b) correct and slower or (c) wrong. After a short fixation (0.4s), another trial started. All timings were fixed in this part. Participants were instructed to always look at the fixation circle in the middle of the screen throughout this and all subsequent tasks. To motivate participants and continued perceptual improvements during the later (reward related) task-stages, participants were told that if they were correct and faster than 1 second in at least 80% of the trials, they will receive an additional monetary bonus of 2 Euros.

The staircasing started after a short training (choosing correct in 8 out of 12 consecutive trials mixed of both contexts) and consisted of two parts: two *adjustment blocks* an two *measurement blocks*. All adjustments of color-speed and motion-coherence followed this formula:

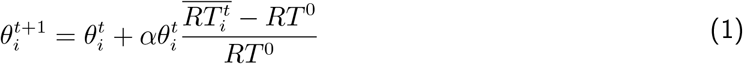

where 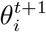 represents the new coherence/speed for motion or color feature *i* during the upcoming time interval/block *t* + 1, 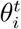 is the level at the time of adjustment, 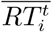 is the mean RT for the specific feature *i* during time interval *t*, *RT*_0_ is the “anchor” RT towards which the adjustment is made and *α* represents a step size of the adjustment, which changed over time as described below.

The basic building block of *adjustment blocks* consisted of 24 cued-feature choices for each context (4 × 3 × 2 = 24, i.e. 4 colors, each discriminated against 3 other colors, on 2 sides of screen). The same feature was not cued more than twice in a row. Due to time constrains, we could not include all possible feature-pairing combinations between the cued and uncued features. We therefore pseudo-randomly choose from all possible background combinations for each feature choice (unlike later stages, this procedure was validated on and therefore included also trials with identical background features). In the first adjustment block, participants completed 72 trials, i.e. 36 color-cued and 36 motion-cued, interleaved in chunks of 4-6 trials in a non-predictive manner. This included, for each context, a mixture of one building block of 2D trials and half a block of 1D trials, balanced to include 3 trials for each cued-feature. 1D or 2D trials did not repeat more than 3 times in a row. At the end of the first adjustment block, the mean RT of the last 48 (accurate) trials was taken as the anchor (*RT*^0^) and each individual feature was adjusted using the above formula with *α* = 1. The second adjustment block started with 24 motion-cued only trials which were used to compute a new anchor. Then, throughout a series of 144 trials (72 motion-cued followed by 72 color-cued trials, all 2D), every three correct answers for the same feature resulted in an adjustment step for that specific feature (Eq. 1) using the average RT of these trials 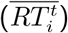 and the motion anchor *RT*^0^ for both contexts. This resulted in a maximum of six adjustment steps per feature, where alpha decreased from 0.6 to 0.1 in steps of 0.1 to prevent over-adjustment.

Next, participants completed two *measurement blocks* identical in structure to the main task (see below) with two exceptions: First, although this was prior to learning the values, they were perceptually cued to chose the feature that later would be assigned with the highest value. Second, to keep the relevance of the feature that later would take the lowest value (i.e. would rarely be chosen), we added 36 additional trials cued to choose that feature (18 motion and 18 color trials per block).

#### Outcome learning task

After the staircasing and prior to the main task, participants learned to associate each feature with a deterministic outcome. Outcomes associated with the four features on each contexts were 10, 30, 50 and 70 credit-points. The value mapping to perceptual features was assigned randomly between participants, such that all possible color- and all possible motion-combinations were used at least once (4! = 24 combinations per context). We excluded motion value-mapping that correspond to clockwise or counter-clockwise ordering. The outcome learning task consisted only of single-feature clouds, i.e. clouds without coherent motion or dots ‘without’ color (gray). Therefore each cloud in this part only represented a single feature. To encourage mapping of the values for each context on similar scales, the two clouds could be either of the same context (e.g. color and color) or from different contexts (e.g. color and motion). Such context-mixed trials did not repeat in other parts of the experiment.

The first block of the outcome learning task had 80 *forced choice* trials (5 repetitions of 16 trials: 4 values × 2 Context × 2 sides of screen), in which only one cloud was presented, but participants still had to choose it to observe its associated reward. These were followed by mixed blocks of 72 trials which included 16 *forced choice* interleaved with 48 *free choice* trials between two 1D clouds (6 value-choices: 10 vs 30/50/70, 30 vs 50/70, 50 vs 70 × 4 context combinations × 2 sides of screen for highest value). To balance the frequencies with which feature-outcome pairs would be chosen, we added 8 forced choice trials in which choosing the lowest value was required. Trials were pseudo-randomized so no value would repeat more than 3 times on the same side and same side would not be chosen more the three consecutive times. Mixed blocks repeated until participants reached at least 85% accuracy of choosing the higher valued cloud in a block, with a minimum of two and a maximum of four blocks. Since all clouds were 1D and choice could be between contexts, these trials started without a cue, directly with the presentation of two 1D clouds (1.6s). Participants then made a choice, and after short fixation (0.2s) were presented with the value of both chosen and unchosen clouds (0.4s, with value of choice marked with a square around it, see Fig. S1). After another short fixation (0.4s) the next trial started. Participants did not collect reward points in this stage, but were told that better learning of the associations will result in more points, and therefore more money later. Specifically, in the MRI experiment participants were instructed that credit points during the main task will be converted into a monetary bonus such that every 600 points they will receive 1 Euro at the end. The behavioral replication cohort received 1 Euro for every 850 points.

#### Main task preparation

In preparation of the main task, participants performed one block of 1D trials at the end of phase 1 and then at the start of the MRI session during the anatomical scan. These blocks were included to validate that changing presentation mediums between phases (computer screen versus projector) did not introduce a perceptual bias to any features and as a final correction for post value-learning RT differences between contexts. Each block consisted of 30 color and 30 motion 1D trials interleaved in chunks of 4-7 trials in a non-predictive manner. The value difference between the clouds was fixed to 20 points (10 repetitions of 3 value comparisons × 2 contexts). Trials were pseudo-randomized so no target value was repeated more than once within context (i.e. not more than twice all in all) and was not presented on the same side of screen more than 3 consecutive trials within context and 4 in total. In each trial, they were first presented with a contextual cue (0.6s) for the trial, followed by short fixation (0.5s) and the presentation of two single-feature clouds of the cued context (1.6s) and had to choose the highest valued cloud. After a short fixation (0.4s), participants were presented with the chosen cloud’s outcome (0.4s). The timing of the trials was fixed and shorter than in the remaining main task because no functional MRI data was acquired during these blocks. Participants were instructed that from the first preparation block they started to collect the rewards. Data from these 1D block were used to inspect and adjust for potential differences between the MRI and the behavior setup. First, participants reacted generally slower in the scanner (*t*(239) = −9.415, *p* < .001, paired t-test per subject per feature). Importantly, however, we confirmed that this slowing was uniform across features, i.e. no evidence was found for a specific feature having more RT increase than the rest (ANOVA test on the difference between the phases, *F*(7,232) = 1.007, *p* = .427). Second, because pilot data indicated increased RT differences between contexts after the outcome learning task we took the mean RT difference between color and motion trials in the second mini-block in units of frames (RT difference divided by the refresh rate), and moved the starting point of each color relative to their target color, the number of frames × its speed. Crucially, the direction of the move (closer/further to target) was the same for all colors, thus ensuring not to induce within-context RT differences.

#### Main task

Finally, participants began with the main experiment inside the scanner. Participants were asked to choose the higher-valued of two simultaneously presented random dot kinematograms, based on the previously learned feature-outcome associations. As described in the main text, each trial started with a cue that indicated the current task context (color or motion). In addition, both clouds could either have two features (each a color and a motion, *2D trials*) or one feature only from the cued context (e.g., colored, but randomly moving dots).

The main task consisted of four blocks in which 1D and 2D trial were intermixed. Each block contained 36 1D trials (3 EV × 2 Contexts × 6 repetitions) and 72 2D trials (3 EV × 2 Contexts × 12 feature-combinations, see fig1c). Since this task took part in the MRI, the duration of the fixation circles were drawn from an truncated exponential distribution with a mean of *μ*=0.6s (range 0.5s-2.5s) for the interval between cue and stimulus, a mean of *μ*=3.4s (1.5s-9s) for the interval between stimulus and outcome and a mean of *μ*=1.25s (0.7s-6s) for the interval between outcome and the cue of the next trial. The cue, stimulus and outcome were presented for 0.6s, 1.6sand 0.8s, respectively. Timing was optimized using VIF-calculations of trial-wise regression models (see Classification procedure section below).

The order of trials within blocks was controlled as follows: the cued context stayed the same for 4-7 trials (in a non-predictive manner), to prevent context confusion caused by frequent switching. No more than 3 repetitions of 1D or 2D trials within each context could occur, and no more than 5 repetition overall. The target did not appear on the same side of the screen on more than 4 consecutive trials. Congruent or incongruent trials did not repeat more than 3 times in a row. In order to avoid repetition suppression, i.e. a decrease in the fMRI signal due to a repetition of information [e.g. 58, 59], no target feature was repeated two trials in a row, meaning the EV could repeat maximum once (i.e. one color and one motion). As an additional control over repetition, we generated 1000 designs according the above-mentioned rules and choose the designs in which the target value was repeated in no more than 10% of trials across trial types, as well as when considering congruent, incongruent or 1D trials separately.

### Behavioral analysis

RT data was analyzed in R (R version 3.6.3 [60], RStudio version 1.3.959 [61]) using linear mixed effect models (lmer in lme4 1.1-21: [62]). When describing main effects of models, the *χ*^2^ represents Type II Wald *χ*^2^ tests, whereas when describing model comparison, the *χ*^2^ represents the log-likelihood ratio test. Model comparison throughout the paper was done using the ‘anova’ function. Regressors were scaled prior to fitting the models for all analyses. The behavioral model that we found to fit the behavioral RT data best was:

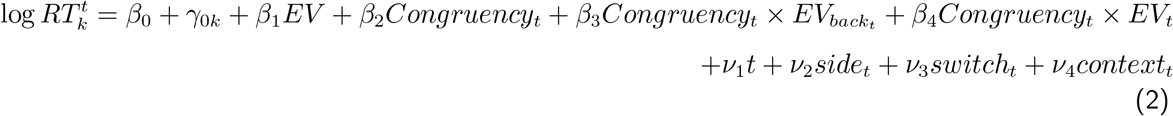

where 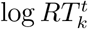 is the log reaction time of subject *k* in trial *t*, *β*_0_ and *γ*_0*k*_ represent global and subject-specific intercepts, *ν*-coefficients reflect nuisance regressors (*side* of target object, trials since last context *switch* and the current *context*), *β*_1_ to *β*_4_ captured the fixed effect of EV, Congruency, Congruency × EV_*back*_ and Congruency × EV, respectively. The additional models reported in the SI included intercept terms specific for each factor level, nested within subject (for EV, Block and Context, see Fig. S2). An exploratory analysis investigating all possible 2-way interactions with all nuisance regressors can be found in Fig. S4.

Investigations of alternative parametrizations of the values can be found in Fig. S3.

Accuracy data was analyzed in R (R version 3.6.3 [60], RStudio version 1.3.959 [61]) using generalized linear mixed effect models (glmer in lme4 1.1-21: [62]) employing a binomial distribution family with a ‘logit’ link function. Regressors were scaled prior to fitting the models for all analyses. No-answer trials of were excluded from this analysis. The model found to fit the behavioral accuracy data best was almost equivalent to the RT model, except for the fourth term involving Congruency × switch:

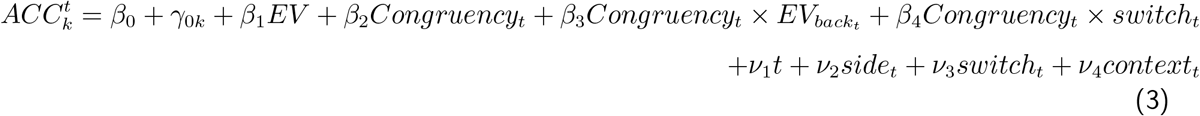

where 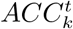 is the accuracy (1 for correct and 0 for incorrect) of subject *k* in trial *t* and all the rest of the regressors are equivalent to Eq. 2. An exploratory analysis investigating all possible 2-way interactions with all nuisance regressors can be found in Fig. S5. We note that the interaction Congruency × switch indicates that participants were more accurate the further they were from a context switch point.

### fMRI data

#### fMRI data acquisition

MRI data was acquired using a 32-channel head coil on a research-dedicated 3-Tesla Siemens Magnetom TrioTim MRI scanner (Siemens, Erlangen, Germany) located at the Max Planck Institute for Human Development in Berlin, Germany. High-resolution T1-weighted (T1w) anatomical Magnetization Prepared Rapid Gradient Echo (MPRAGE) sequences were obtained from each participant to allow registration and brain surface reconstruction (sequence specification: 256 slices; TR = 1900 ms; TE = 2.52 ms; FA = 9 degrees; inversion time (TI) = 900 ms; matrix size = 192 x 256; FOV = 192 x 256 mm; voxel size = 1 x 1 x 1 mm). This was followed with two short acquisitions with six volumes each that were collected using the same sequence parameters as for the functional scans but with varying phase encoding polarities, resulting in pairs of images with distortions going in opposite directions between the two acquisitions (also known as the blip-up / blip-down technique). From these pairs the displacements were estimated and used to correct for geometric distortions due to susceptibility-induced field inhomogeneities as implemented in the the fMRIPrep preprocessing pipeline. In addition, a whole-brain spoiled gradient recalled (GR) field map with dual echo-time images (sequence specification: 36 slices; A-P phase encoding direction; TR = 400 ms; TE1 = 4.92 ms; TE2 = 7.38 ms; FA = 60 degrees; matrix size = 64 x 64; 619 FOV = 192 x 192 mm; voxel size = 3 x 3 x 3.75 mm) was obtained as a potential alternative to the method described above. However, this GR frield map was not used in the preprocessing pipeline. Lastly, four functional runs using a multi-band sequence (sequence specification: 64 slices in interleaved ascending order; anterior-to-posterior (A-P) phase encoding direction; TR = 1250 ms; echo time (TE) = 26 ms; voxel size = 2 x 2 x 2 mm; matrix = 96 x 96; field of view (FOV) = 192 x 192 mm; flip angle (FA) = 71 degrees; distance factor = 0, MB acceleration factor = 4). A tilt angle of 30 degrees from AC-PC was used in order to maximize signal from the orbitofrontal cortex (OFC, see [63]). For each functional run, the task began after the acquisition of the first four volumes (i.e., after 5.00 s) to avoid partial saturation effects and allow for scanner equilibrium. Each run was about 15 minutes in length, including a 20 seconds break in the middle of the block (while the scanner is running) to allow participants a short break. We measured respiration and pulse during each scanning session using pulse oximetry and a pneumatic respiration belt part of the Siemens Physiological Measurement Unit.

#### BIDS conversion and defacing

Data was arranged according to the brain imaging data structure (BIDS) specification [64] using the HeuDiConv tool (version 0.6.0.dev1; freely available from https://github.com/nipy/heudiconv). Dicoms were converted to the NIfTI-1 format using dcm2niix [version 1.0.20190410 GCC6.3.0; [65]]. In order to make identification of study participants highly unlikely, we eliminated facial features from all high-resolution structural images using pydeface (version 2.0; available from https://github.com/poldracklab/pydeface). The data quality of all functional and structural acquisitions were evaluated using the automated quality assessment tool MRIQC [for details, [see 66], and the MRIQC documentation]. The visual group-level reports confirmed that the overall MRI signal quality was consistent across participants and runs.

#### fMRI preprocessing

Data was preprocessed using *fMRIPrep* 1.2.6 ([67]; [68]; RRID:SCR_016216), which is based on *Nipype* 1.1.7 ([69]; [70]; RRID:SCR_002502). Many internal operations of *fMRIPrep* use *Nilearn* 0.5.0 [71, RRID:SCR_001362], mostly within the functional processing workflow.

Specifically, the T1-weighted (T1w) image was corrected for intensity non-uniformity (INU) using `N4BiasFieldCorrection` [72, ANTs 2.2.0], and used as a T1w-reference throughout the workflow. The anatomical image was skull-stripped using `antsBrainExtraction.sh` (ANTs 2.2.0), using OASIS as the target template. Brain surfaces were reconstructed using `recon-all` [FreeSurfer 6.0.1, RRID:SCR_001847, 51], and the brain masks were estimated previously was refined with a custom variation of the method to reconcile ANTs-derived and FreeSurfer-derived segmentations of the cortical gray-matter of Mindboggle [RRID:SCR_002438, 50]. Spatial normalization to the ICBM 152 Nonlinear Asymmetrical template version 2009c [73, RRID:SCR_008796] was performed through nonlinear registration with `antsRegistration` [ANTs 2.2.0, RRID:SCR_004757, 74], using brain-extracted versions of both T1w volume and template. Brain tissue segmentation of cerebrospinal fluid (CSF), white-matter (WM) and gray-matter (GM) was performed on the brain-extracted T1w using `fast` [FSL 5.0.9, RRID:SCR_002823, 75].

To preprocess the functional data, a reference volume for each run and its skull-stripped version were generated using a custom methodology of *fMRIPrep*. A deformation field to correct for susceptibility distortions was estimated based on two echo-planar imaging (EPI) references with opposing phase-encoding directions, using `3dQwarp` [76] (AFNI 20160207). Based on the estimated susceptibility distortion, an unwarped BOLD reference was calculated for a more accurate co-registration with the anatomical reference. The BOLD reference was then co-registered to the T1w reference using `bbregister` (FreeSurfer), which implements boundary-based registration [77]. Co-registration was configured with nine degrees of freedom to account for distortions remaining in the BOLD reference. Head-motion parameters with respect to the BOLD reference (transformation matrices, and six corresponding rotation and translation parameters) are estimated before any spatiotemporal filtering using `mcflirt` [FSL 5.0.9, 78]. BOLD runs were slice-time corrected using `3dTshift` from AFNI 20160207 [76, RRID:SCR_005927] and aligned to the middle of each TR. The BOLD time-series (including slice-timing correction) were resampled onto their original, native space by applying a single, composite transform to correct for head-motion and susceptibility distortions. First, a reference volume and its skull-stripped version were generated using a custom methodology of *fMRIPrep*.

Several confound regressors were calculated during preprocessing: Six head-motion estimates (see above), Framewise displacement, six anatomical component-based noise correction components (aCompCorr) and 18 physiological parameters (8 respiratory, 6 heart rate and 4 of their interaction). The head-motion estimates were calculated during motion correction (see above). Framewise displacement was calculated for each functional run, using the implementations in *Nipype* [following the definitions by 79]. A set of physiological regressors were extracted to allow for component-based noise correction [*CompCor*, 80]. Principal components are estimated after high-pass filtering the BOLD time-series (using a discrete cosine filter with 128s cut-off) for the two *CompCor* variants: temporal (tCompCor, unused) and anatomical (aCompCor). For aCompCor, six components are calculated within the intersection of the aforementioned mask and the union of CSF and WM masks calculated in T1w space, after their projection to the native space of each functional run (using the inverse BOLD-to-T1w transformation). All resamplings can be performed with *a single interpolation step* by composing all the pertinent transformations (i.e. head-motion transform matrices, susceptibility distortion correction, and co-registrations to anatomical and template spaces). Gridded (volumetric) resamplings were performed using `antsApplyTransforms` (ANTs), configured with Lanczos interpolation to minimize the smoothing effects of other kernels [81]. Lastly, for the 18 physiological parameters, correction for physiological noise was performed via RETROICOR [82, 83] using Fourier expansions of different order for the estimated phases of cardiac pulsation (3rd order), respiration (4th order) and cardio-respiratory interactions (1st order) [84]: The corresponding confound regressors were created using the Matlab PhysIO Toolbox ([85], open source code available as part of the TAPAS software collection: https://www.translationalneuromodeling.org/tapas. For more details of the pipeline, and details on other confounds generated but not used in our analyses, see [the section corresponding to workflows in fMRIPrep’s documentation](https://fmriprep.readthedocs.io/en/latest/workflows.html).

For univariate analyses, BOLD time-series were re-sampled to MNI152NLin2009cAsym standard space in the *fMRIPrep* pipeline and then smoothed using SPM [86, SPM12 (7771)] with 8mm FWHM, except for ROI generation, where a 4mm FWHM kernel was used. Multivariate analyses were conducted in native space, and data was smoothed with 4mm FWHM using SPM [86, SPM12 (7771)]. Classification analyses further involved three preprocessing steps of voxel time-series: First, extreme-values more than 8 standard deviations from a voxels mean were corrected by moving them by 50% their distance from the mean towards the mean (this was done to not bias the last z scoring step). Second, the time-series of each voxel was detrended, a high-pass filter at 128 Hz was applied and confounds were regressed out in one action using *Nilearn* 0.6.2 [71]. Lastly, the time-series of each voxel for each block was z scored.

### Univariate fMRI analysis

All GLMs were conducted using SPM12 [86, SPM12 (7771)] in MATLAB [55]. All GLMs consisted of two regressors of interest corresponding to the onsets of the two trial-types (1D/2D, except for one GLM where 2D onsets were split by Congruency) and included one parametric modulator of EV assigned to 1D onset and different combinations of parametric modulators of EV, Congruency, EV_back_ and their interactions (see Fig. S10 for GLM visualization). All parametric modulators were demeaned before entering the GLM, but not orthogonalized. Regressors of no interest reflected cue onsets in Motion and Color trials, stimulus onsets in wrong and no-answer trials, outcome onsets and 31 nuisance regressors (e.g. motion and physiological parameters, see fMRI-preprocessing). The duration of stimulus regressors corresponded to the time the stimuli were on screen. The durations for the rest of the onset regressors were set to 0. Microtime resultion was set to 16 (64 slices / 4 MB factor) and microtime onset was set to the 8 (since slice time correction aligned to middle slice, see fMRI-preprocessing). Data for all univariate analyses were masked with a whole brain mask computed as intercept of each functional run mask generated from fMRIprep [50, 51]. MNI coordinates were translated to their corresponding brain regions using the automated anatomical parcellation toolbox [87–89, AAL3v1] for SPM. We verified the estimability of the design matrices by assessing the Variance Inflation Factor (VIF) for each onset regressor in the HRF-convolved design matrix. Specifically, for each subject, we computed the VIF (assisted by scripts from https://github.com/sjgershm/ccnl-fmri) for each regressor in the HRF-convolved design matrix and averaged the VIFs of corresponding onsets across the blocks. None of the VIFs surpassed a value of 3.5 (a value of 5 is considered a conservative indicator for overly colinear regressors, e.g. [90], see Fig.S10 for details). Detailed descriptions of all GLMs are reported in the main text. Additional GLMs verifying the lack of Congruency in any frontal region can be found in Fig.S10.

#### vmPFC functional ROI

In order to generate a functional ROI corresponding to the vmPFC in a reasonable size, we re-ran the GLM with only EV modulators (i.e. this GLM had no information regarding the contextually irrelevant context) on data that was smoothed at 4mm. We then threshold the EV contrasts for 1D and 2D trials (EV_1D_ + EV_2D_ > 0) at *p* < .0005. The group ROI was generated in MNI space and included 998 voxels. Multivariate analyses were conducted in native space and the ROI was transformed to native space using ANTs and nearest neighbor interpolation [ANTs 2.2.0 74] while keeping only voxels within the union of subject- and run-specific brain masks produced by the fMRIprep pipeline [50, 51]. The resulting subject-specific ROIs therefore had varying number of voxels (*μ* = 768.14, *σ* = 65.62, min = 667, max = 954).

#### Verifying design trial-wise estimability

To verify that the individual trials are estimatable (for the trial-wise multivariate analysis) and as a control over multi-colinearity [90], we convolved a design matrix with the HRF for each subject with one regressor per stimuli (432 regressors with duration equal to the stimulus duration), two regressor across all cues (split by context) and three regressor for all outcomes (one for each EV). We then computed the VIF for each stimulus regressor (i.e. how predictive is each regressor by the other ones). None of the VIFs surpassed 1.57 across all trials and subjects (*μ_VIF_* = 1.42, *σ_VIF_* = .033, min = 1.34). When repeating this analysis with a GLM in which also outcomes were split into trialwise regressors, we found no stimuli VIF larger than 3.09 (*μ_VIF_* = 2.64, *σ_VIF_* = .132, min = 1.9). Note that 1 is the minimum (best) value and 5 is a relatively conservative threshold for colinearity issues ([e.g. 90]). This means that the BOLD responses of individual trials can be modeled separately and should not have colinearity issues with other stimuli nor with the outcome presentation of each trial.

### Multivariate analysis

#### Classification procedure

The training set for Value and Context classifiers consisted of fMRI data from behaviorally accurate 1D trials. For each trial, we took the TR corresponding to approx. 5 seconds after stimulus onset (*round*(*onset* + 5)) to match the peak of the Haemodynamic Response Function (HRF) estimated by SPM [86]. Training of Value and Context classifiers was done using a leave-one-run-out scheme across the four runs with 1D trials. To avoid bias in the training set after sub-setting only to behaviorally accurate trials (i.e. over-representation of some information) we up-sampled each training set to ensure equal number of examples in the training set for each combination of EV (3), Context (2) and Chosen-Side (2). Specifically, if one particular category was less frequent than another (e.g., more value-30, left, color trials than value-50, left-color trials) we up-sampled that example category by randomly selecting a trial from the same category to duplicate in the training set, whilst prioritising block-wise balance (i.e., if one block had 2 trials in the chunk and another block had only 1, we first duplicated the trial from under-represented block etc.). We did not up-sample the testing set. The EV_back_ classifiers were trained on behaviorally accurate 2D trials (5 seconds after stimulus onset) and up-sampled by EV (3), Context (2) and EV_back_ (3) (without Chosen-Side as this resulted in excluding many subjects for lack of trials in some training sets). Due to strong imbalance of unique examples of EV_back_ in the training sets (see below) we trained 3 one-vs-rest classifiers, each tasked with identifying one level of EV_back_. This required to adjust the sample weights in order to account for the higher frequency of the ‘rest’ compared to the ‘one’ label.

Decoding was conducted using multinomial logistic regression as implemented in *scikit-learn* 0.22.2 [91], using a *C* parameter of 1.0, L2 regularization and the lbgfs solver. For each test example (i.e. trial) we obtained the predicted probability per class. To avoid numerical issues in the subsequent modeling of the classifier’s predictions, probabilities were constrained to lie within 0.00001 and 0.99999, rather than 0 and 1. In addition to the probabilities, we obtained the balanced classification accuracy (i.e. is the class with the highest probability also the correct class of the test trial). We separately averaged classification for each participant, test fold and label (this ensured controlling for any label imbalance in the testing set).

Finally, before modelling the probabilities using linear mixed effects models, we averaged the classifiers probabilities across the nuisance effects, i.e. we obtained one average probability for each combination of relevant and irrelevant values. Crossing each level of EV (three levels) with each level of irrelevant value of the chosen side combined with irrelevant value of the non-chosen side (12 level, see Fig. 1), resulted in 36 combinations per participant. Note that the relevant value of the unchosen cloud was always EV – 20 and therefore we did not include this as a parameter of interest. After averaging, we computed for each combination of values the EV_back_, Congruency and alternative parameters (see Fig. S8). The main model comparison, as well as the lack of effects of any nuisance regressor, was confirmed on a dataset with raw, i.e. non-averaged, probabilities (see Fig S6 and S8). Because in the one-vs-rest training of EV_back_ classifiers the three class probabilities for each trial were obtained independently, they sum to 1. We therefore first normalized the probabilities for each testing trial.

Probabilities were analyzed in R (R version 3.6.3 [60], RStudio version 1.3.959 [61]) with Generalized Linear Mixed Models using Template Model Builder (glmmTMB, [92]) models, employing a beta distribution family with a ‘logit’ link function. When describing main effects of models, the *χ*^2^ represents Type II Wald *χ*^2^ tests, whereas when describing model comparison, the *χ*^2^ represents the log-likelihood ratio test. Model comparison throughout the paper was done using the ‘anova’ function. Throughout all the analyses, each regressor was scaled prior to fitting the models. Lastly, for the analysis of behavioral accuracy (Fig. 6) we also included behaviorally wrong trials.

#### Value similarity analyses

asked whether the predicted probabilities reflected the difference from the objective probability class. The model we found to best explain the data was:

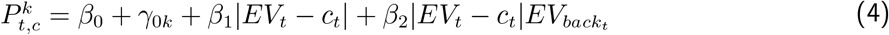

where 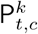, is the probability that the Value classifier assigned to class *c* in trial *t* for subject *k*, *β*_0_ and *γ*_0*k*_ represent global and subject-specific intercepts, |*EV_t_* − *Class_c,t_*| is the absolute difference between the EV of the trial and the class the probability is assigned to and |*EV_t_* − *Class_c,t_*|*EV_back_t__* is the interaction of this absolute difference with EV_back_. For models nested in the levels of EV, we included *ζ*_0*k,EV*_, which is the EV-specific intercept nested within each within each subject level. In these models, testing for main effects of EV_back_ or Congruency was not sensible because both factors don’t discriminate between the classes, but rather assign the same value to all three probabilities from that trial (which sum to 1).

For the feature similarity model we substituted |*EV_t_* − *c_t_*| with a “similarity” parameter that encoded the perceptual similarity between each trial in the test set and the perceptual features that constituted the training examples of each class of the classifier. For 1D trials, this perceptual parameter was identical to the value similarity parameter (|*EV_t_* − *c_t_*|). This was because from the shown pairs of colors, both colors overlapped between training and test if the values were identical; one color overlapped if the values were different by one reward level (e.g. a 30 vs 50 comparison corresponded to two trials that involved pink vs green and green vs orange, i.e. sharing the color green); and no colors overlapped if the values were different by two levels (30 vs 70). On 2D trials however, due to changing background features and their value-difference variation, perceptual similarity of training and test was not identical to value similarity. Even though both the value similarity and the perceptual similarity parameter correlated (*ρ* = .789, *σ* = .005), we found that the value similarity model provided a better AIC score (value similarity AIC: −3898, Feature similarity AIC: −3893, Fig. 4). Detailed description with examples can be found in Fig. S6. Crucially, even when keeping the value difference of the irrelevant features at 20, thus limiting the testing set only to trials with feature-pairs that were included in the training, our value similarity model provided a better AIC (−1959) than the feature similarity model (−1956). To test for a perceptual alternative of EV_back_ we substituted the corresponding parameter from the model with Similarity_back_. This perceptual parameter takes on 1 if the perceptual feature corresponding to the EV_back_ appeared in the 1D training class (as highest or lowest value) and 0 otherwise. As described in the main text, none of the perceptual-similarity encoding alternatives provided a better fit than our models that focused on the expected values the features represented.

#### Modelling the influence of irrelevant values and Context signals on EV representation

The following model of the probability of the objective EV was found to explain the data best:

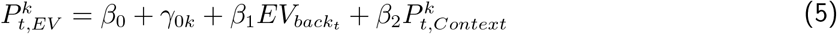

where 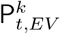 is the probability assigned to the objective class by the Value classifier (corresponding to EV of the trial *t*) for subject *k*, *β*_0_ and *γ*_0*k*_ represent global and subject-specific intercepts, EV_*back*_ is the maximum of the two ignored values (or the EV of the contextually irrelevant context) and 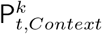 is the probability assigned to the objective class by the Context classifier (logit-transformed, i.e. 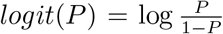, and scaled for each subject). For models nested in the levels of EV, we included *ζ*_0_*k,EV*__ which is EV specific intercept nested within each within each subject level (see Fig. S8). Investigations of alternative parametrizations of the values can be found in Fig. S8. Including an additional regressor that encoded trials in which EV=EV_back_ (or: match) which did not improve model fit, and no evidence for an interaction of the match regressor with the EV_back_ was found (LR test with added terms: 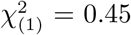, *p* = .502, 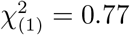, *p* = .379, respectively). This might indicate that when value expectations of both contexts matched, there was neither an increase nor a decrease of *P*_EV_.

To compute the correlations between each pair of classes we transformed the probabilities for each class using a multinomial logit transform. For example, for class 30 we performed probabilities were transformed with 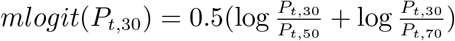. To examine the relationship between EV and EV_back_, we only included 2D trials in which EV ≠ EV_back_. This allowed us to categorize all three probabilities as either EV, EV_back_ or Other, whereby Other reflected the value that was neither the EV, nor the EV_back_. To prevent bias we included only trials in which Other was presented on screen (as relevant or irrelevant value). We then averaged across nuisance regressors (see Classification procedure) and computed the correlation across all trials (Spearman rank correlation). Lastly, we Fisher z-transformed the correlations 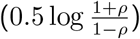 to approximate normality for the t test. To validate these results, we performed an additional model comparison in which we added a term of the logit transformed P*_EV_back__* or of P_*other*_ to Eq. 5 (*β*_2_*mlogit*(*P_t,EV_back__*) or *β*_2_*mlogit*(*P_t,Other_*) respectively). As reported in the main text, adding a term reflecting P_*EV_back_*_ resulted in a smaller (better) AIC score than when we added a term for P_*other*_ (−567,−475, respectively). This was also preserved when running the analysis including nuisance regressors (see *v*s in Eq. 2) on the non-averaged data (AICs: −5913.3,−5813.3). We note that subsetting the data the way we did resulted in a strong negative correlation in the design matrix between EV and EV_back_ (*ρ* = −0.798, averaged across subjects). Although this should not directly influence our interpretation, we validated the results by using alternative models with effects hierarchically nested within the levels of EV *and* EV_back_ (Averaged data AICs: −560, −463, Raw data AICs: −5906.8,−5804.3)

As previously clarified, 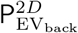 was derived from a classifier trained on 2D trials. We note that the mixed evidence in favor for direct EV_back_ decoding might relate to the fact that the number of unique examples for each class of EV_back_ differed drastically (due to our design, see Fig. 1c) which motivated us to split the decoding of EV_back_ to three classifiers, each trained on a different label (see ‘Classification procedure’). However, our approach of combining one-vs-rest training with oversampling and sample weights could not fully counteract these imbalances and the probabilities each classifier assigned to its corresponding class 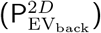 were still biased by class imbalances. Specifically, the correlation of 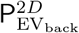 and EV_back_ was *ρ_μ_* = .26, *ρ_σ_* = .07 across subjects, where ‘2D’ indicates the classifier was directly trained on 2D trials, unlike with P_EV_back__ which comes from a classifier trained on EV in 1D trials. Since in this analysis we were mainly interested in the neural representation of EV_back_ regardless of whether EV_back_ was 30, 50 or 70 in given trial, we solved this issue by using mixed effect models and setting a random intercept for each level of EV_back_ (i.e. running the models nested within the levels of EV_back_).

Thus, when testing across the levels of EV_back_, the model that best explained the data was:

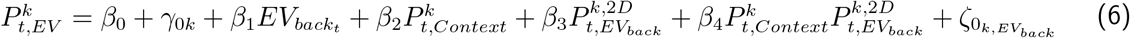

where similar to Eq. 5, 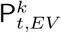, is the probability assigned to the EV class by the Value classifier for trial *t* and subject *k*, *β*_0_ and *γ*_0*k*_ represent global and subject-specific intercepts and 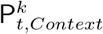 is the logit-transformed probability assigned to Context class. 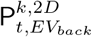 is the probability the EV_back_ classifier assigned the correct class (in main text: 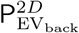, where 2D notes that this classifier was trained on 2D trials) and *ζ*_0_*k,EV_back_*__ is EV_back_ specific intercept nested within each within each subject level.

#### Linking MRI effects to behavior

When modelling the probability of EV_back_ from the Value classifier (P_EV_back__, Fig. 6a.), we did not average across nuisance regressors. Our baseline model was: 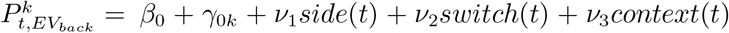. Neither including a main effect nor interactions between EV, EV_back_ and Congruency improved model fit. When including behaviorally wrong trials in the model, we used `drop1` in combination with *χ*^2^-tests from `lmer4` package [62] to test which of the main effects or interactions improves the fit. This resulted in the following model as best explaining the data:

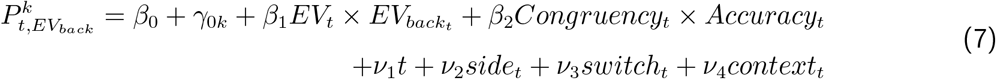

where 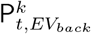 is the probability the Value classifier assigned to the EV_*back*_ class (corresponding to EV_*back*_ of trial *t*) for subject *k, β*_0_ and *γ*_0*k*_ represent global and subject-specific intercepts, EV is the maximum of the two relevant and EV_*back*_ is the maximum of the two ignored values. Congruency reflects whether the actions chosen in the relevant vs. irrelevant context would be the same, and the Accuracy regressor has 1 if participants chose the highest relevant value and 0 otherwise. We note that the interaction EV × EV_*back*_ (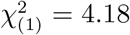, *p* = .041) indicates higher in trials in which EV and EV_*back*_ were more similar, the probability assigned to EV_back_ was higher. However, we find this effect hard to interpret since this corresponds to the value similarity effect we previously reported.

In order to investigate the effect of vmPFC neural representations on behavioral accuracy, we used hierarchical model comparison to directly test the influence of neural representation of EV, EV_back_ and Context on behavioral accuracy separately for congruent and incongruent trials (Fig. 6b-c.). First, we tested if adding *logit*(*P_t,Context_*), *mlogit*(*P_t,EV_*) or *mlogit*(*P_t,EV_back__*) to Eq. 3, would help to explain the behavioral accuracy better. Because the analysis was split for congruent and incongruent trials, we excluded the terms involving a Congruency effect. For incongruent trials, only *logit*(*P_t,Context_*) improved the fit (LR-tests: *logit*(*P_t,context_*): 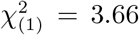, *p* = .055, *mlogit*(*P_t,EV_*): 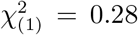, *p* = .599, *mlogit*(*P_t,EV_back__*): 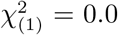, *p* = .957). In a second step we then separately tested the interactions *logit*(*P_t,context_*) × *mlogit*(*P_t,EV_*) or *logit*(*P_t,context_*) × *mlogit*(*P_t,EV_back__*) and found that only the latter had improved the fit 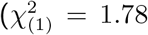, *p* = .183, 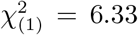, *p* = .012, respectively). For congruent trials, only *mlogit*(*P_t,EV_back__*) and marginally *mlogit*(*P_t,EV_*) improved the fit (LR-tests: *logit*(*P_t,Context_*): 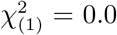, *p* = .922, *mlogit*(*P_t,EV_*): 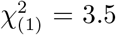, *p* = .061, *mlogit*(*P_t,EV_back__*): 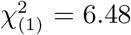, *p* = .011). In a second step we tested separately the interactions *logit*(*P_t,Context_*) × *mlogit*(*P_t,EV_*), *logit*(*P_t,Context_*) × *mlogit*(*P_t,EV_back__*) or *mlogit*(*P_t,EV_back__*) × *mlogit*(*P_t,EV_*) and found none of these improved model fit when adding them to a model that included both main effects from the previous step 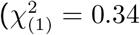, *p* = .560, 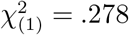, *p* = .598, 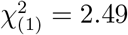, *p* = .115, respectively).

To investigate the effect of vmPFC neural representations on RT in behaviorally accurate trials, we asked whether subjects who had a stronger effect of Context representation (P_context_) on EV representation (P_EV_) or a stronger Spearman rank correlation between P_EV_ and P_EV_back__ (taken from the Value classifier) also had a stronger effect of Congruency on their RT. Additionally, we asked whether subjects who had a stronger effect of EV_back_ on P_EV_ and or a stronger effect of 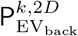 on P_EV_ also had a stronger modulation of EV_back_ on the Congruency RT effect. To obtain subject specific effect of Congruency on RT we added *γ*_1*k*_*Congruency* and *γ*_2*k*_*CongruencyEV_back_t__* to the RT model (Eq. 2), representing subject-specific slopes of Congruency for subject *k* and for the interaction of Congruency and EV_back_, respectively. The subject-specific correlation of P_*EV*_ and P_*EV_back_*_ was estimated by using only trials in which EV ≠ EV_back_. Probabilities were multinomial logit transformed and correlations were Fisher z-transformed (see above) before averaging across trials to achieve one correlation value per subject. In the main text and in Fig 5 we did not average the data to achieve maximum sensitivity to trial-wise variations. The results reported in the main text replicate when running the same procedure while averaging the data across nuisance regressors following the multinomial logit transformation (*R* = .38, *p* = .023). To extract subject-specific slopes for the effect of EV_back_ on P_EV_ we included a term for this effect (*γ*_1*k*_*EV_back_t__*) in Eq. 5, but due to convergence issues during model fitting, we had to drop the subject-specific intercept (*γ*_0*k*_) in that model. Similarly, to extract subject-specific slopes for the effect of 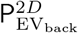 on P_EV_ we included a term for this effect 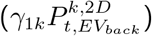 in Eq. 6.

## Supplementary Information

**Figure S1:**
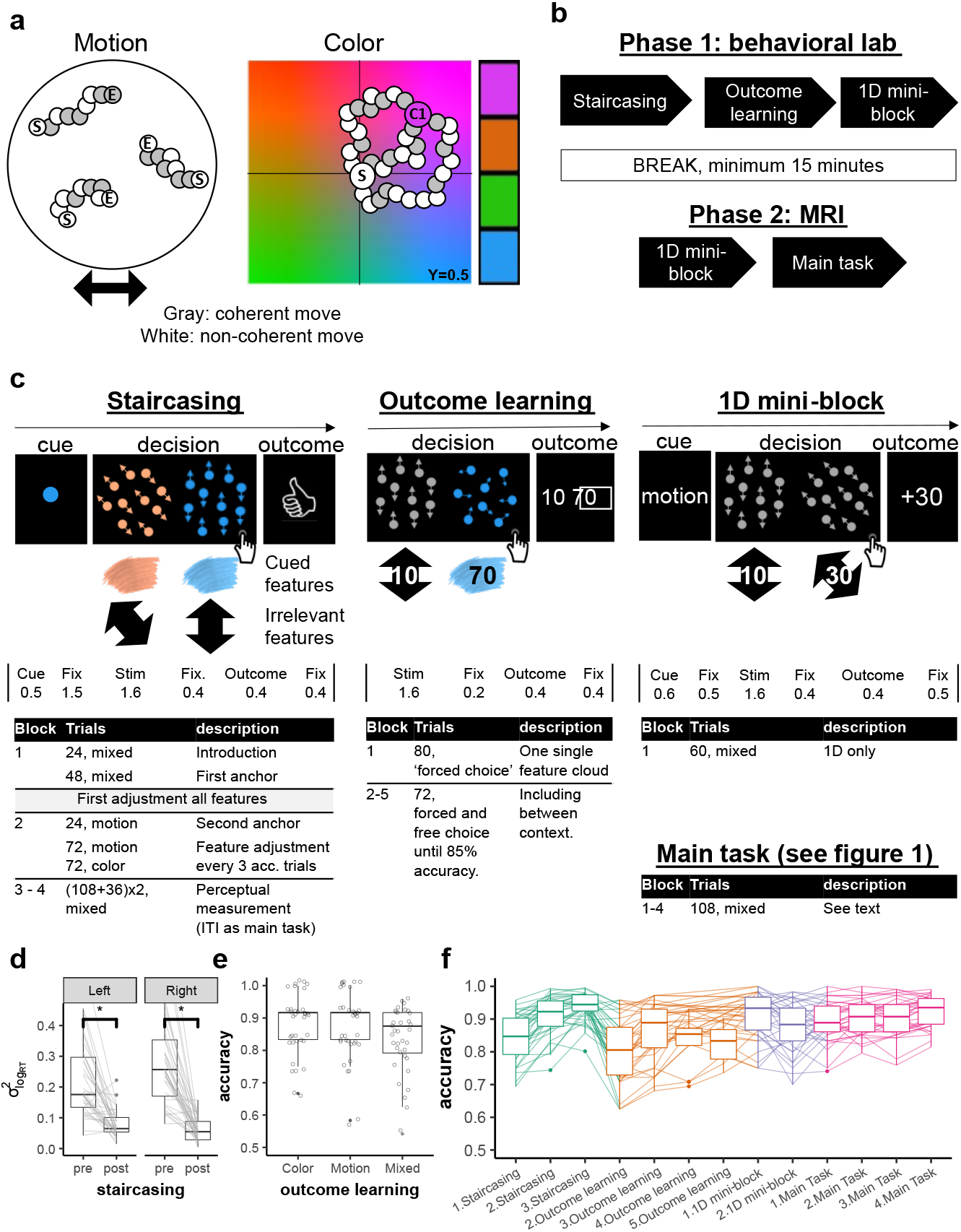
Full procedure and experimental design for all phases, related to Fig 1. **a.** Brownian algorithm for color and motion. Each illustration shows the course of 3 example dots; ‘S’ and ‘E’ marked dots reflect Start and End positions, respectively. Remaining dots represent location in space for different frames. Left panel: Horizontal motion trial. Shown are framewise dot positions between start and end. In each frame, a different set of dots moved coherently in the designated direction (gray) with a fixed speed; remaining dots moved in a random direction [conceptually taken from 41]. Right panel: Example of a pink color trial. We simulated the YCbCr color space that is believed to represent the human perception in a relative accurate way [cf. 57]. A fixed luminance of Y = 0.5 was used. For technical reasons we sliced the X-axis by 0.1 on each side and the Y-axis by 0.2 from the bottom of the space to ensure the middle of the space remained gray given the chosen luminance. In each frame, a different set of dots (always 30% of the dots) moved coherently towards the target color in a certain speed whereas the rest were assigned with a random direction. All target colors were offset by 23.75%from the center towards each corner. Right bar illustrates the used target colors. **b.** Full procedure. The experiment consisted of two phases, the first one took place in the behavioral lab and included Staircasig, Outcome-learning and the first 1D mini-block. The second took place inside the MRI scanner and consisted of the second 1D mini-block and the main task. **c.** Example trial procedures and timing of the different tasks. Timing of each trial is depicted below illustrations. **Staircasing (left)** Each trial started with a cue of the relevant feature. Each cloud had one or two features (motion and/or color) and participants had to detect the cued feature. Participants’ task was to choose the cued feature (here: blue). After a choice, participants received feedback if they were correct and faster than 1 second, correct and slower, or wrong. **Outcome learning (middle)** Participants were presented with either one or two single-feature clouds and asked to chose the highest valued feature. Following their choice, they were presented with the values of both clouds, with the chosen cloud’s associated value marked with a square around it. The pair of shown stimuli included across contexts comparisons, e.g. between up/right and blue, as shown. **1D mini block (right)** At the end of the first phase and beginning of the second phase participants completed a mini-block of 60 1D trials during the anatomical scan (30 color-only, 30 motion-only, interleaved). Participants were again asked to make a value-based two alternative forced choice choice decision. In each trial, they were first presented with a contextual cue (color/motion), followed by the presentation of two single-feature clouds of the cued context. After a choice, they were presented with the chosen-cloud’s value. No BOLD response was measured during these blocks and timing of the trials was fixed and shorter than in the main task (see Main task preparation in online methods) **Main task (bottom)** This part included 4 blocks, each consisting of 36 1D and 72 2D trials trials presented in an interleaved fashion (see online method and Fig. 1). **d.** Button specific reduction in RT variance following the staircasing. We verified that the staircasing procedure also reduced differences in detection speed between features when testing each button separately. Depicted is the variance of reaction times (RTs) across different color and motion features (y axis). While participants’ RTs were markedly different for different features before staircasing (pre), a significant reduction in RT differences was observed after the procedure (post, *p* < .001.) **e.** Choice accuracy in outcome learning trials. Participants achieved near ceiling accuracy in choosing the highest valued feature in the outcome learning task, also when testing for color, motion and mixed trials separately (*p*s< .001). Mixed trials only appeared in this part of the experiment to encourage mapping of the values on similar scales. **f.** Accuracy throughout the experiment, plotted for each block of each part of the experiment. *In the staircasing (left)* High accuracy for the adjustment and measurement blocks (2-3) ensured that there were no difficulties in perceptual detection of the features. *In Outcome learning* a clear increase in accuracy throughout this task indicated learning of feature-outcome associations. Note that Block 5 of this part was only included for those who did not achieve 85% accuracy beforehand. Starting the *1D mini blocks* (middle) and throughout *themain task* (right) until the end of the experiment high accuracy. *μ* and *σ* from left to right: Staircasing: .84,.07;.91,.06;.94,.04; Outcome Learning: .81,.1;.86,.09;.83,.08;.82,.06; 1D mini blocks: .91,.07;.88,.08; Main task: .89,.06;.91,.05;.9,.06;.92,.05.

**Figure S2:**
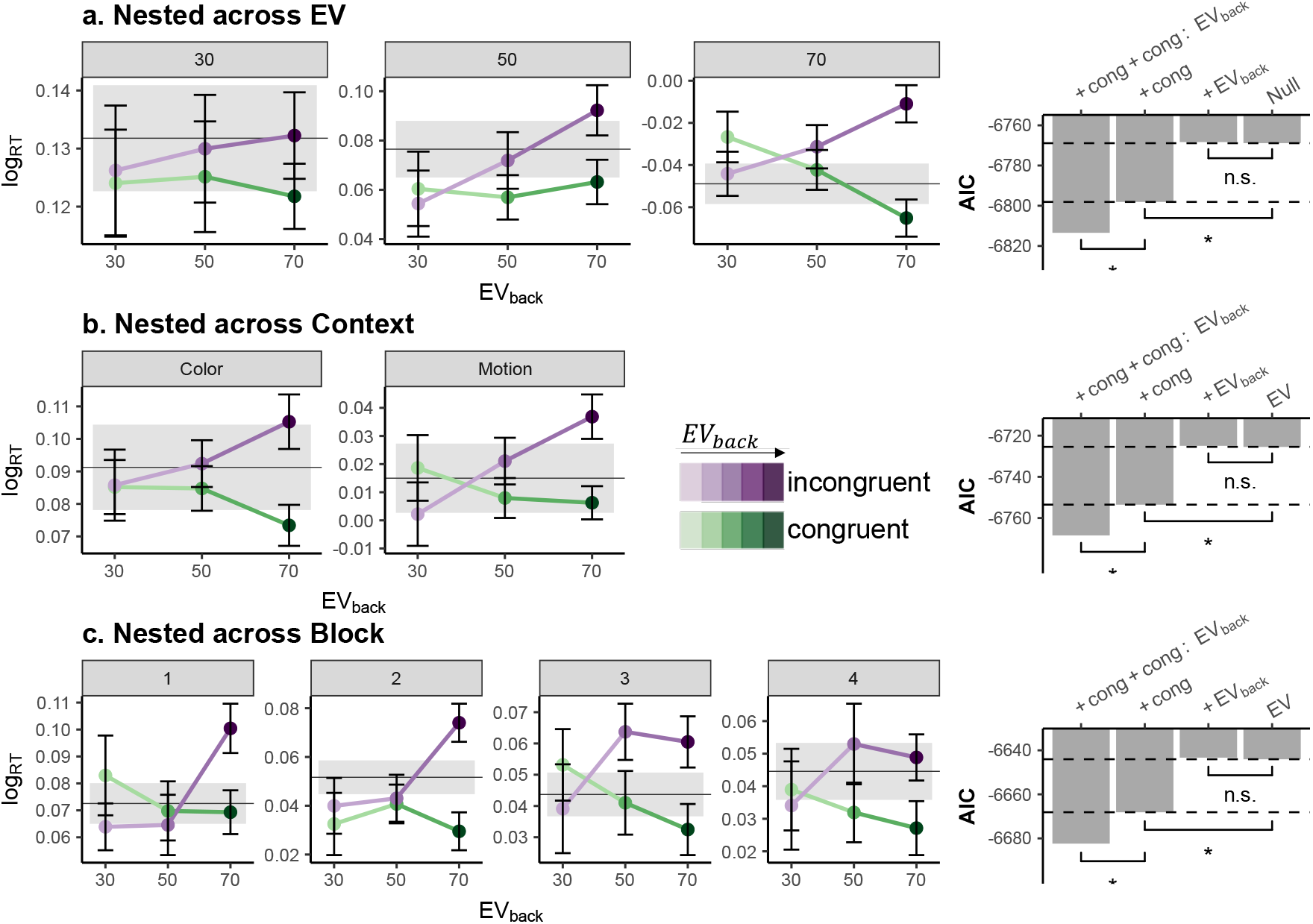
Nested RT models, related to Fig 2. **a-c. Nested models within Factors.** Each row represents one congruency analysis, done separately for each level of expected value (top row), context (middle) or block (bottom). The RT effect of Congruency × EV_back_ is shown on the left, corresponding AICs for mixed effect models with nested factors are shown on the right. Error bars represent corrected within subject SEMs [42, 43]. Mean RT (line) and SEM (shades) for the corresponding 1D trials is plotted in gray for each panel (e.g. mean across all 1D trials where EV=30 are on top left panel). Null models shown on the right are identical to Eq. 2, albeit included *ζ*_0_*k_v_*__, which is the factor-specific (*v*) intercept nested within each within each subject level (see online methods). Likelihood ratio tests were performed to asses improved model fit when adding (1) Congruency or (2) EV_back_ terms to the Null model and when adding (3) Congruency × EV_back_) in addition to Congruency. Stars represent p values less than .05. For nested within EV, the Null model did not include a main effect for EV and the LR test was: (1) 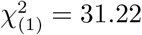, *p* < .001; (2) 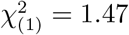, *p* = .226; (3) 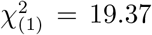, *p* < .001; For models nested within Context the LR test was: (1) 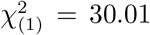, *p* < .001; (2) 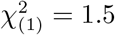, *p* = .22; (3) 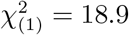, *p* < .001; and for Block: (1) 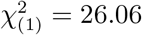, *p* < .001; (2) 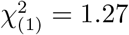, *p* = .26; (3) 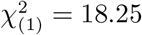, *p* < .001; In the first row (nested across EV) the interaction with EV is visible, i.e. the higher the EV, the stronger our effects of interests were.

**Figure S3:**
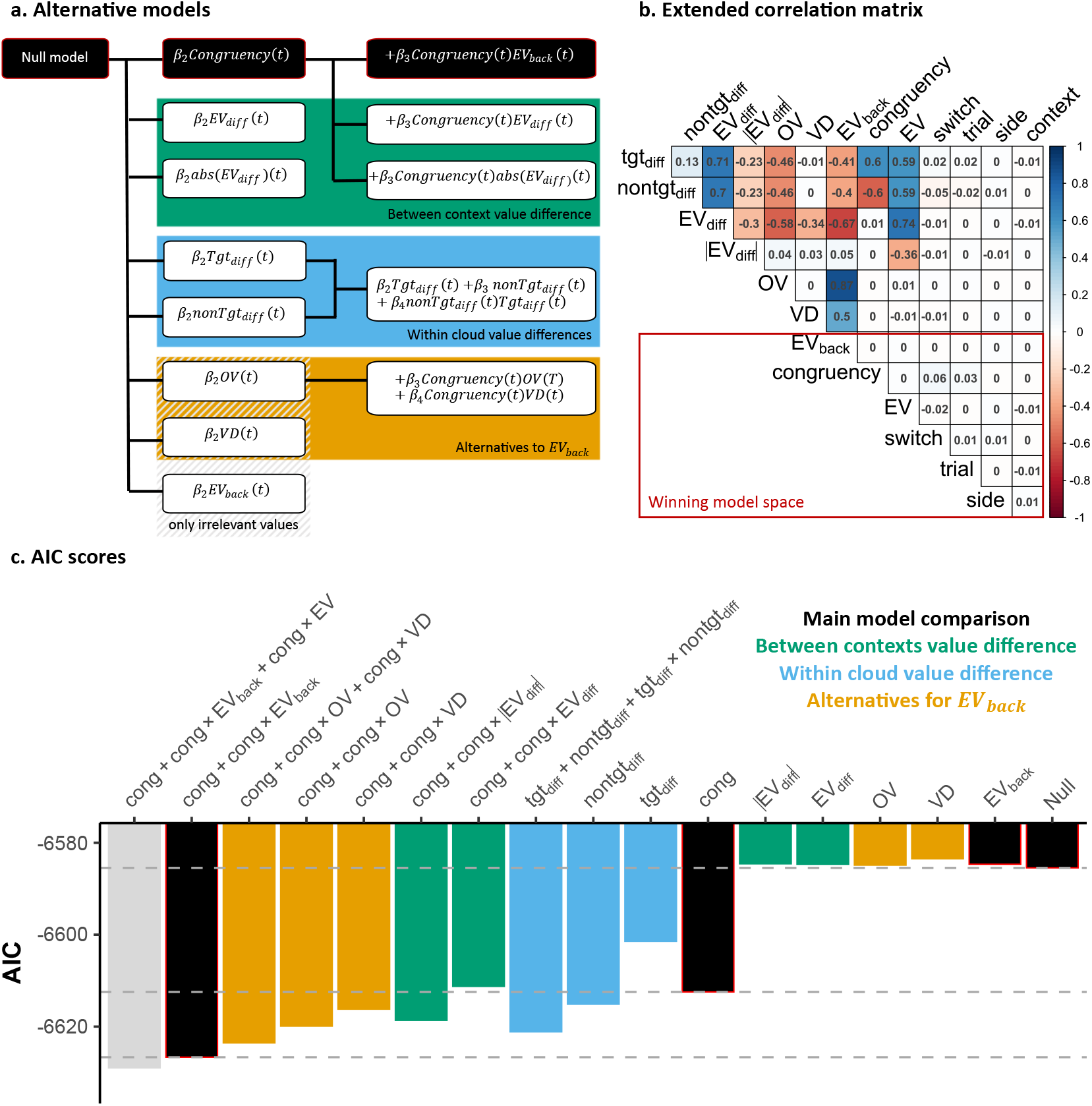
Alternative RT models, extended RT model comparisons and correlation matrix of all regressors, related to Fig 2. **a.** Alternative mixed effect models, each represented as a row which lists main factors of interest. We clustered different alternative models into three classes: *Green models* included factors that reflected the difference between the expected values of both contexts (EV – EV_back_, including unsigned EV factors); *blue* models include instead factor that reflect the value-difference between context within each cloud where ‘tgt’ (target) is the chosen cloud with the highest value according to the relevant context and *orange* models included two alternative parameterization of values in the non-relevant context: irrelevant features’ Value Difference (VD) and Overall Value (OV), which are also orthogonal to Congruency (Cong), and to each other. *In black* is the main model comparison as presented in the main text. **b. Extended correlation matrix**. Averaged correlation across subjects of all scaled regressors for accurate 2D trials (models’ input). Marked in red rectangle are main factors of the experiment which are orthogonal by design and used for the model comparison reported in the Main Text. **c. AIC scores.** We tested different alternatives shown in (a) in a stepwise hierarchical model comparison, as in the main text. Each bar represents the AIC (y-axis) of a different model (x-axis) where the labels on the x-axis depict the added terms to the Null model for that specific model. The Null model included nuisance regressors and the main effect of EV (see *ν* and *β*_1_ in Eq. 2). The models described in the main text are shown in black. The gray model includes the additional term for Congruency × EV. Dashed lines correspond to the AIC values of the models used in the main text. Importantly, no main effect representing only the contextually irrelevant values (VD, OV, EV_back_) nor the difference between the EVs (EV_diff_,|EV_diff_|, also when excluding EV from the null model, not presented) improved model fit over the Null model. This supports our finding that neither large irrelevant values, nor their similarity to the objective EV, influenced participants’ behavior. Similar to EV_back_, factors from the green and orange clusters are also orthogonal to Congruency, which allowed us to test their interaction. Factors from the blue cluster highly correlate with both Congruency (and EV_back_) and therefore were tested separately. Non of the alternatives provided a better AIC score (y axis, lower is better).

**Figure S4:**
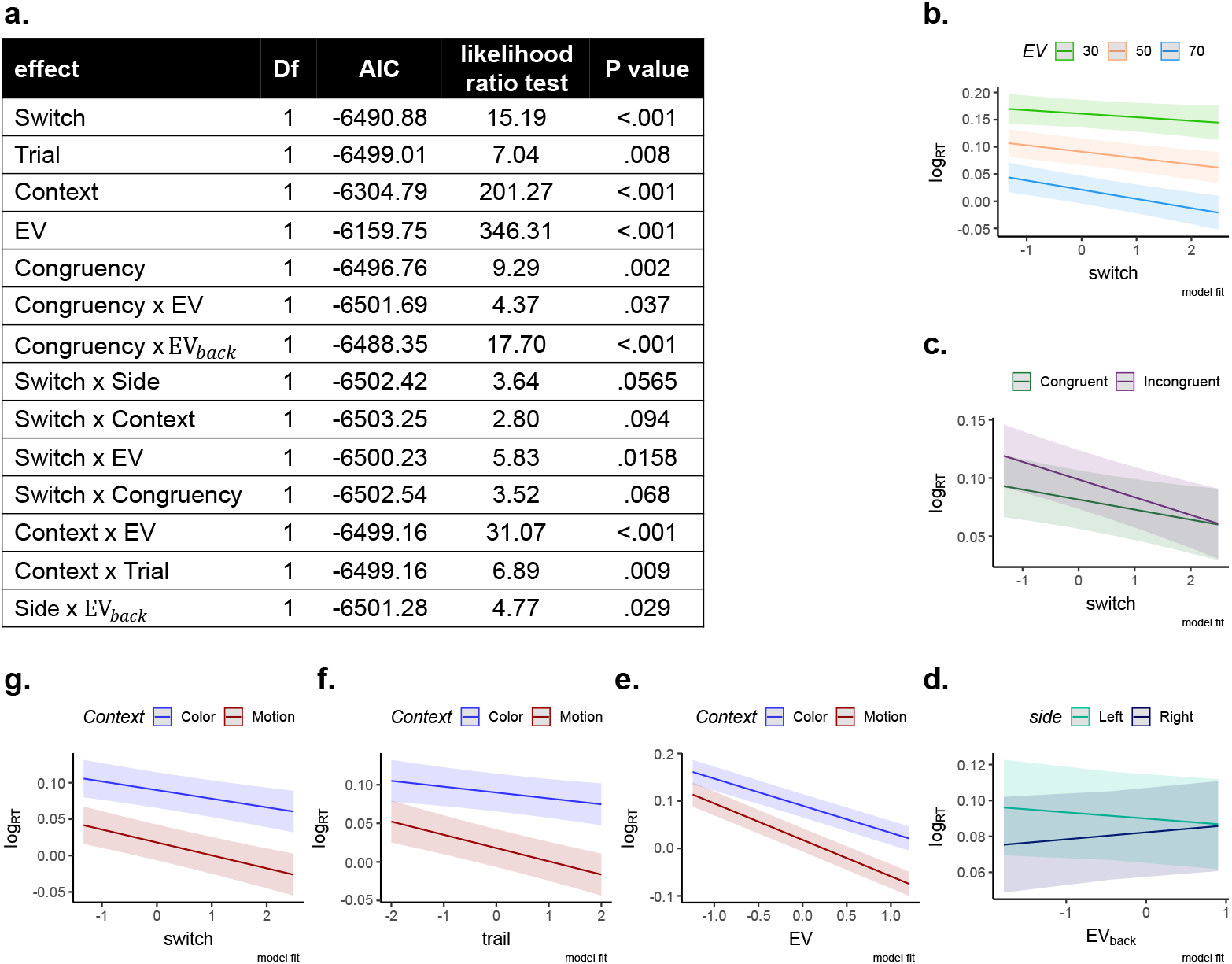
Exploratory analysis of RT model presented in Main Text, related to Fig 2. **a.** The table presents the individual contribution of terms taken from Eq. 2 and all possible two-way interactions to the model fit using the drop1 function in R [60]. In short, this exploratory analysis started with a model that included all main effects from Eq. 2 and all possible 2-way interaction between them and tested which terms contribute to the fit. If a term did not improve fit, it was dropped from the model. Presented are all effects with a p value less than *p* < .01 . **b-g.** Model fits of all effects with *p* < .1. X-axes are normalized (as in the model) and y-axes reflect RTs on a log scale (model input). Clockwise from the top: RTs became progressively faster with increasing trials since the context switch. This effect was possibly stronger for higher EV (b) and for incongruent trials (c). We note that our experiment was not designed to test the effect of the switch. (d) An interaction of Side and EV_back_ was found, for which we offer no explanation. Panels (e) to (g) reflect interaction of context with EV (e), trial (f), and switch (g). We note that due to the used perceptual color space there might be a context-specific ceiling effect in RTs due to training throughout the task which could have induced effects of context. Specifically, since dots start gray and slowly ‘gain’ the color, it might take a few frames until there is any evidence for color. However, the motion could be theoretically detected already on the second frame (since coherence was very high). This could explain why some effects that represent decrease in RT might hit a boundary for color (and not motion). Crucially, we refer the reader to supplementary Fig S2 where the main model comparison hold also when we ran the model nested within the levels of Context

**Figure S5:**
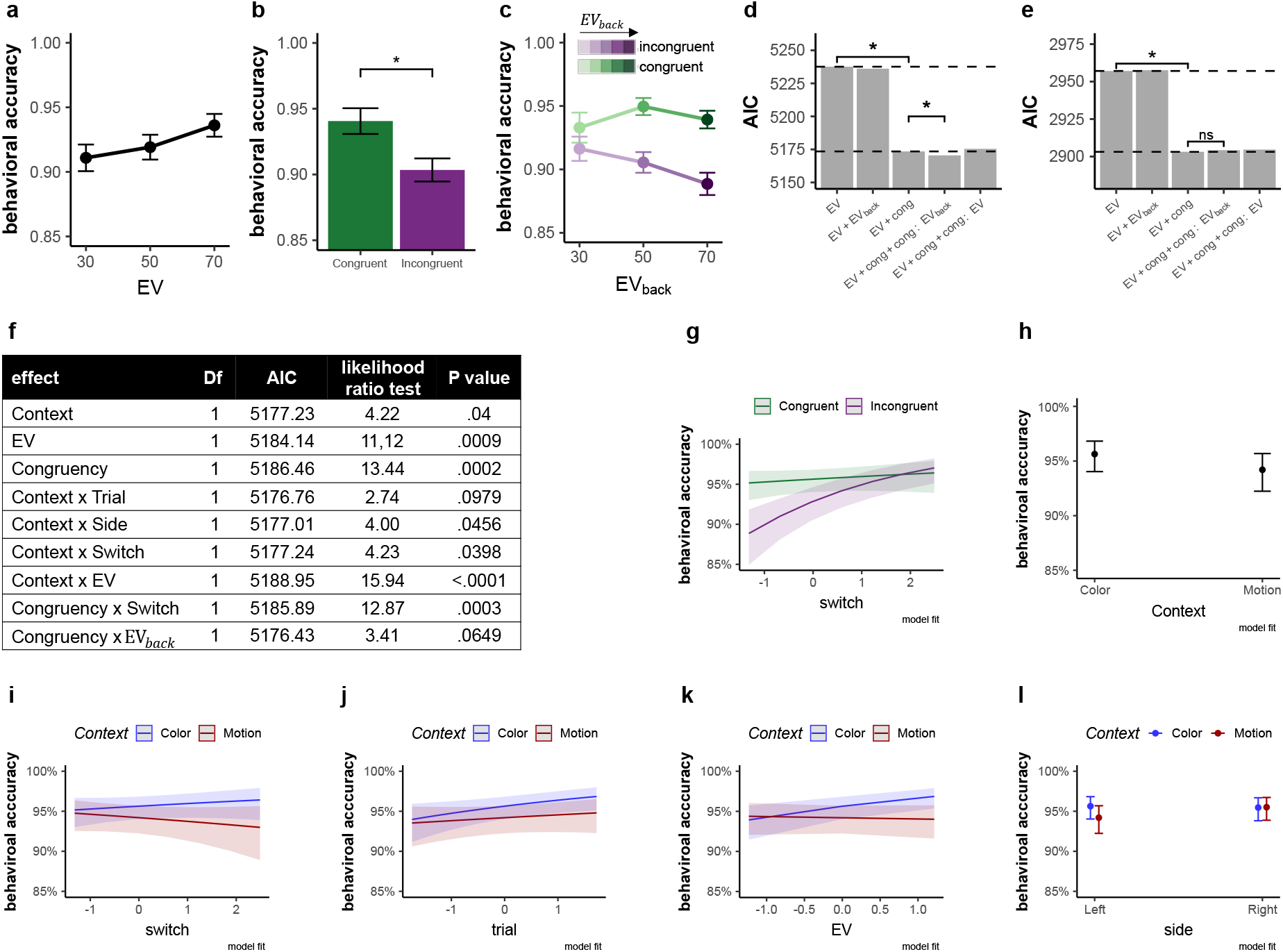
Behavioral accuracy results: related to Fig 2. **a.** Comparison of accuracy (y-axis) for each level of EV (x-axis) showed that participants were more accurate for higher EV, *p* = .001. **b.** Comparison of congruent versus incongruent trials also revealed a performance benefit of the former, *p* = .001. **c.** The effect of Congruency was modulated by EV_back_, i.e. the more participants could expect to receive from the ignored context, the less accurate they were when the contexts disagreed (x axis, shades of colours). Further investigations revealed that the modulation of EV_back_ is likely limited to Incongruent trials 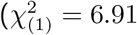, *p* = .009, when modeling only Incongruent trials), yet does not increase accuracy for Congruent trials 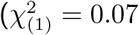, *p* = .794, when modeling only congruent trials), likely due to a ceiling effect. Error bars in panels a-c represent corrected within subject SEMs [42, 43]. **d.** Hierarchical model comparison of choice accuracy, similar to the RT model reported in the main text. These analyses showed that including Congruency improved model fit (*p* < .001). Including the additional interaction of Congruency × EV_back_ improved the fit even more (*p* = .03). **e.** We replicated the choice accuracy main effect in an independent sample of 21 participants outside of the MRI scanner, i.e. including Congruency improved model fit 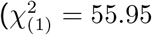, *p* < .001). We did not find a main effect of EV on accuracy in this sample 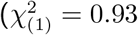, *p* = .333). The interaction term Congruency × EV_back_ did not significantly improve fit in this sample. Modeling only Incongruent trials, as above, reveled that EV_back_ had a marginal effect on accuracy 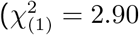, *p* = .088). Near-ceiling accuracies in Congruent trials in combination with a smaller sample might have masked the effects. **f.** The table presents the individual contribution of terms taken from Eq. 3 and all possible two-way interactions to the model fit using the drop1 function in R [60]. In short, this exploratory analysis started with a model that included all main effects from Eq. 3 and all possible 2-way interaction between them and tested which terms contribute to the fit. If a term did not improve fit, it was dropped from the model. Subsequent panels present all the effects corresponding to *p* < .01. Note that this is a non-hypothesis driven exploration of the data and that accuracy was very high in general throughout the main task. **g.** Accuracy as a function of time since switch. Akin to RTs, accuracy increased with number of trials since the last context switch, mainly for incongruent trials. **h.** Context effect on accuracy. According to the exploratory model, participants were slightly more accurate in color than in motion trials. However, a direct paired t test between average accuracy of color compared to motion was not significant (*t*_(34)_ = 0.96, *p* = .345) **i-l.** Depicted are some minor interactions of no interest with Context, according to the exploratory model.

**Figure S6:**
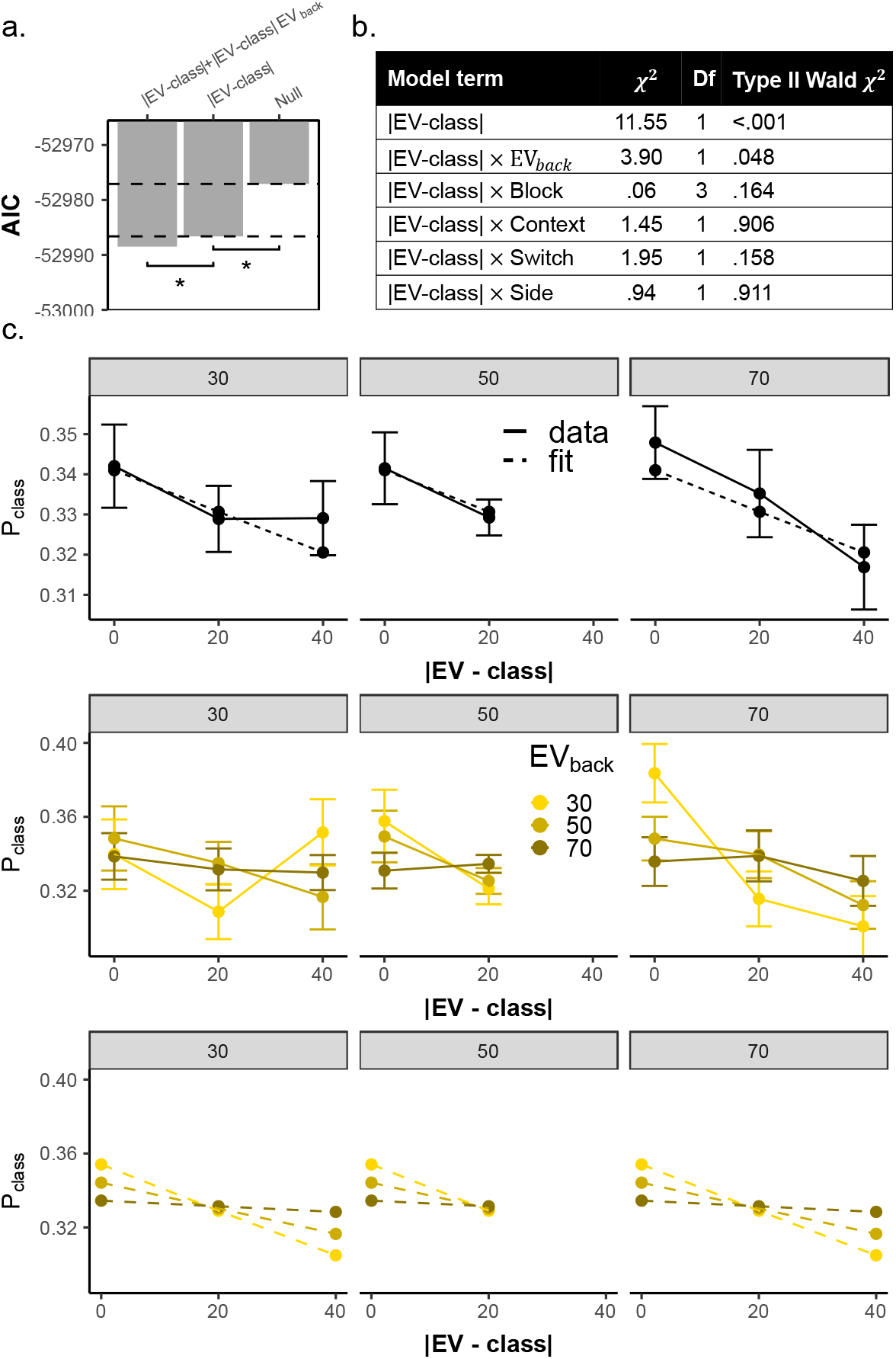
Supplementary information for Value similarity analysis: related to Fig. 4. **a.** Main value similarity model comparison replicated when fitting the models to unaveraged data. Adding a term for |EV-class| improved model fit (LR test with added term: 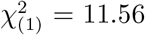, *p* < .001). Adding an additional term for |EV-class| × EV_back_ further improved the fit 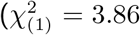, *p* = .049), as in the model reported in the main text (Fig. 4b). **b.** Effect of Nuisance regressors on unaveraged data (t, Side, Switch and Context). Same as Congruency and EV_back_, all of the nuisance regressors don’t discriminate between the classes, but rather assign the same value to all three probabilities from that trial (which sum to 1). We therefore tested if any of them modulated the value similarity effect. As can be seen in the table, none of the nuisance regressors modulated the value similarity effect. **c.** Replication of the value similarity model comparison reported in the main text, averaged across nuisance regressors and nested within the levels of EV, i.e. including EV-specific intercepts nested within each within each subject level *ζ*_0_*k*_v___, see Online Methods). As in the analysis reported in the Main Text, adding a main effect for |EV-Class| improves model fit (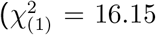, *p* < .001, first row) as well as adding an additional interaction term |EV-class| × EV_back_ (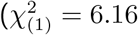, *p* = .013, middle row shows data, bottom row shows model fit. Error bars represent corrected within subject SEMs [42, 43])

**Figure S7:**
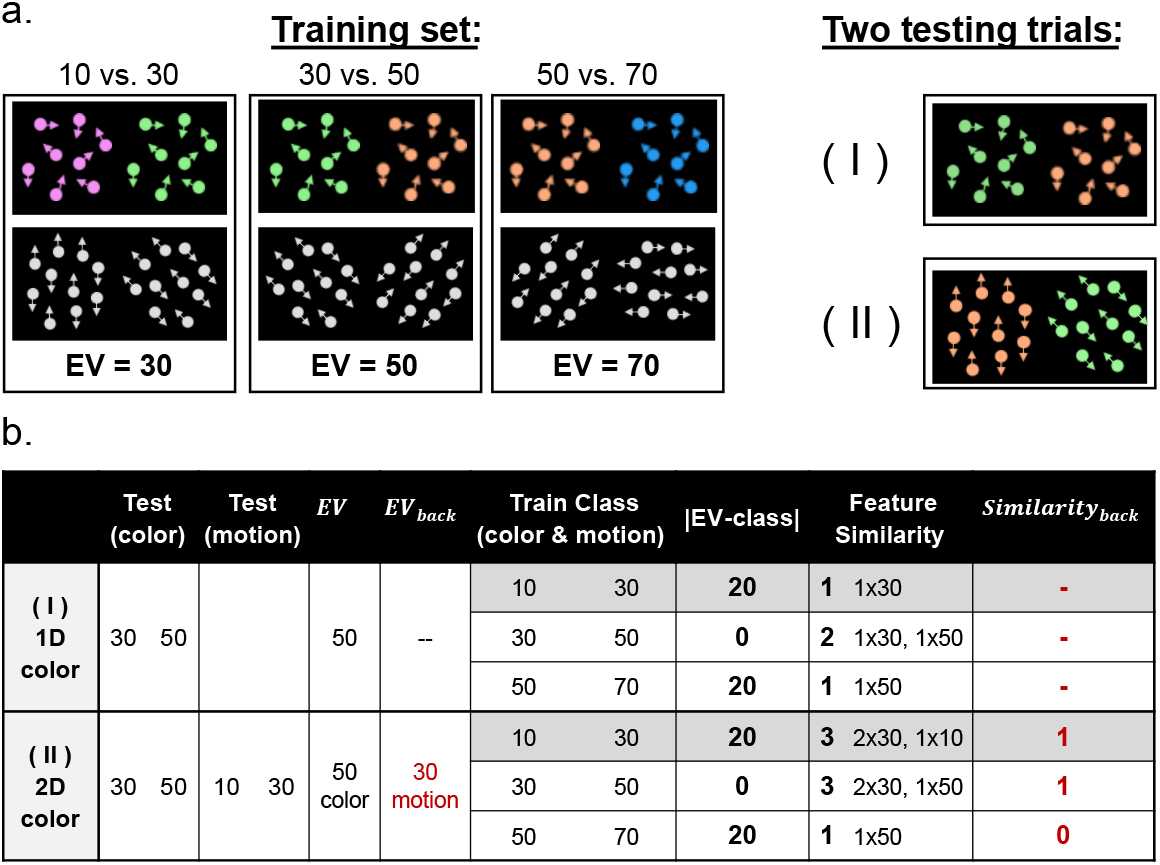
Supplementary information for perceptual similarity analysis: related to Fig. 4. To control that our EV classifier was indeed sensitive to values and not only to the perceptual features the values were associated with, we compared this value similarity model to a perceptual models that merely encodes the amount of perceptual overlap between each training class and 2D testing (irrespective of their corresponding values) and found that our model explained the data best (see 4c.). Replacing the EV_back_ with a parameter that encodes the presence of the perceptual feature corresponding to EV_back_ in the training class (Similarity_back_: 1 if the feature was preset, 0 otherwise did not provide a better AIC score (−3897.1) than including the value of EV_back_ (−3902.5). **a.** Left: training set consisting of 1D trials provided for the classifier for each class (in the experiment the sides were pseudorandomised). Note that each class had the same amount of color and motion 1D trials and that the value difference between the values was always 20. Right: two examples of 2D trials that constituted the classifier test set. **b.** The table illustrates the calculation of feature similarity between classifier test and training in two example trials in one 1D and one 2D trial. Specifically, shown are the corresponding values and features for each trial with the predicted values at each class for the parameters value similarity (|EV-class|), feature similarity and similarity_back_. Feature similarity encodes the perceptual overlap between the shown test example and the training examples underlying with each value class. The first row shows a case in which the classifier was tested on a 1D green vs. orange color trial ( 30 vs 50, EV = 50). Considering in this case for instance the predicted probability that EV=30, the table illustrates the training example underlying the EV = 30 cases (10 vs 30, dark gray shading), the |EV-class| (here: 20, because 50-30), and the feature similarity i.e. how many features from the training class appeared in the test example (here: 1). The second row shows a 2D color trial, reflecting the same value based choice between 30 and 50. The value similarity between training and test stays the same as for the 1D trial shown above. However, the feature similarity between test and training changes because of the motion features. If we take class 30 for example (which is 10 vs 30, dark gray shading), the feature 30 appeared twice (color and motion) and the feature 10 appeared once (motion), i.e. feature similarity now takes on the value 3. Similarity_back_ was used to test a perceptual-based alternative to the EV_back_ parameter. Similarity_back_ takes on 1 if the perceptual feature corresponding to the EV_back_ appeared in the training class and 0 otherwise (red text in table). As described in the main text, none of the perceptual-similarity encoding alternatives provided a better fit than the reported models that focused on the values the features represent.

**Figure S8:**
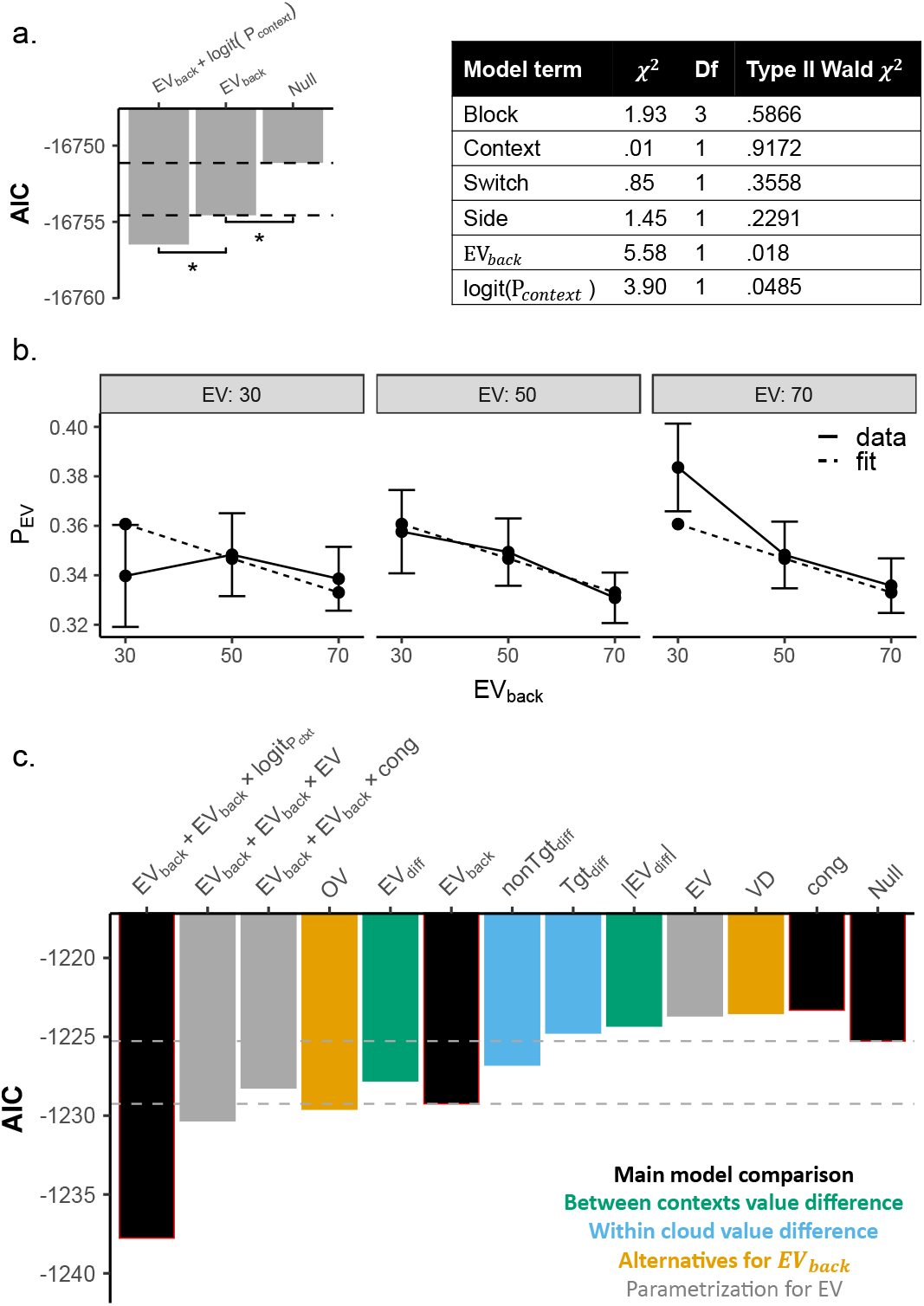
Modelling probability assigned to the EV class: related to Fig. 5. **a.** We replicated the main results using the unaveraged data. The Null model was: 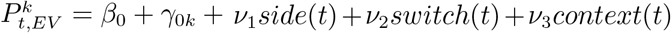, where 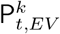 is the probability assigned to the class corresponding to the EV of trial *t* for subject *k*, *β*_0_ and *γ*_0*k*_ represent global and subject-specific intercepts. Side, Switch and Context are the same as in the RT model (Eq. 2); None of these variables had a main effect, *p* > 0.4 (see table, right). The factor *trial* could not be included due to model convergence issues. Adding a term representing EV_back_ improved model fit (LR test including term: 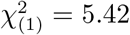, *p* = .019). Adding an additional term for context decodability further improved the fit (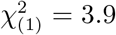, *p* = .048). The table (right) displays the Type 2 Wald *χ*^2^ test for all main effects from the model. **b.** Depicted is the effect of EV_back_ (x-axis) on the probability assignd to the EV class (P_*EV*_, y axis). Solid lines represent the data and dashed lines the model fit of a model that included random effects of subject and EV nested within subject (data averaged across nuisance regressors, adding a main effect for EV_back_ improved model fit (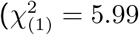, *p* = .014). Error bars represent corrected within subject SEMs [42, 43]. **c.** Similar to our analysis of alternative models of RT, we clustered models reflecting alternative explanations into three conceptual groups (see color legend; cf. Fig. S3a). All models were fitted to the probability assigned to the objective EV in accurate 2D trials, similar to Eq. 5. Each column represents the AIC (y-axis) of a different model (x-axis) where the labels on the x-axis depict all the main effects included in that specific model (i.e. added to the Null, i.e. Eq. 5 without any main effects). We found no evidence that any other parameters explain the data better than the ones we used in the main text. Specifically, only including main effect of EV_back_, Overall Value of the irrelevant values (OV) and the difference of both EVs (EV_*diff*_) provided a better AIC score than the Null model. Note that adding OV (−1229.6) only slightly surpassed EV_back_ (−1229.26). Crucially, the correlation of EV_back_ and OV is very high (*ρ* = .87, see main text). We then looked at possible interactions with the EV_back_ effect. Congruency did not seem to modulate the main effect of EV_back_ and adding an interaction term EV × EV_back_ provided a slightly better AIC (−1230.33), yet this effect was not significant (LR test: 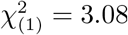, *p* = .079). Section (b) also visualizes this effect. Lastly, adding a term for the Context decodability provided the lowest (i.e. best) AIC score. This exploratory analysis revealed that our model provides the best fit for *P*_EV_ in all cases except when EV_back_ was replaced with the sum of irrelevant values (−1229.6, −1229.2, respectively, Fig. S8). In contrast, AIC scores of behavioral models’ favored EV_back_ as modulator of Congruency, over the sum of irrelevant values (−6626.6, −6619.9, respectively, Fig.S3). However, both parameters were strongly correlated (*ρ* = .87, *σ* = .004) and therefore our task was not designed to distinguish between these two alternatives.

**Figure S9:**
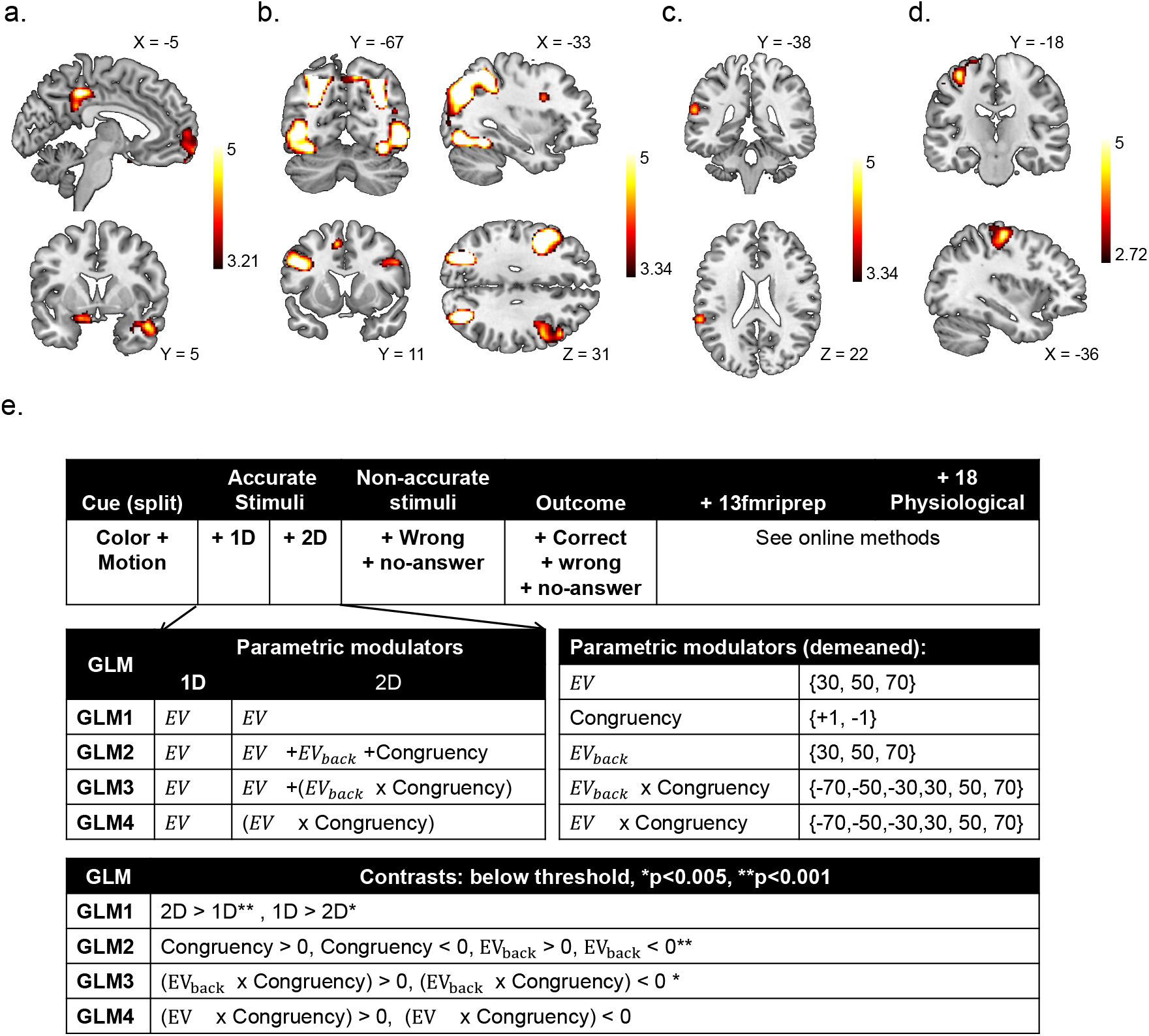
Main univariate results. The main analyses indicated that multiple value expectations are represented in parallel within vmPFC. Here, we asked whether whole-brain univariate analyses could also uncover evidence for processing of multiple value representations. In particular, we asked whether we could find evidence for a single representation that integrates the multiple value expectations into one signal. To this end, we first analyzed the fMRI data using GLMs with separate onsets and EV parametric modulators for 1D and 2D trials (see below for detailed description). **a**. The intersection of the EV parametric modulators of 1D and 2D trials (EV_1D_ > 0 ∩ EV_2D_ > 0) revealed several regions including right Amygdala, bilateral Hippocampus and Angular Gyrus, the lateral and medial OFC and overlapping vmPFC. Hence, the vmPFC signaled the expected value of the current context in both trial types as expected – even though 2D trials likely required higher attentional demands (see panel b). Voxelwise threshold *p* < .001, FDR cluster-corrected. **b** 2D trials were characterized by increased activation in an attentional network involving occipital, parietal and frontal clusters (2D > 1D, *p* < .001 FDR cluster corrected). Next, we searched for univariate evidence of processing irrelevant values by modifying the parametric modulators assigned to 2D trials in the above-mentioned GLM. Specifically, in addition to EV_2D_, we added Congruency (+1 for congruent and −1 for incongruent) and EV_back_ as additional modulators of the activity in 2D trials. This GLM revealed no evidence for a Congruency contrast anywhere in the brain (even at a liberal voxel-wise threshold of *p* < .005). **c.** An unexpected negative effect of EV_back_ was found in the Superior Temporal Gyrus (*p* < .001), i.e. the higher the EV_back_, the lower the signal in this region. *p* < .001, FDR cluster-corrected. No overlap with (b), see S10. We note that this is similar to previous reports implicating this region in modelling choices of others [93]). Notably, unlike the multivariate analysis, no effect in any frontal region was observed. Motivated by our behavioral analysis, we then turned to look for the interaction of each relevant or irrelevant value with Congruency. An analysis including only a Congruency × EV_2D_ parametric modulator revealed no cluster (even at *p* < .005). **d.** A cluster in the primary motor cortex was negatively modulated by Congruency × EV_back_, i.e. the difference between Incongruent and Congruent trials increased with higher EV_back_, similar to the RT effect and akin to a response conflict, *p* < .005, FDR cluster-corrected. No overlap with (b), see S10 Lastly, we re-ran all above analyses concerning Congruency and EV_back_ only inside the identified vmPFC ROI. No voxel survived for Congruency, EV_back_ nor the interactions, even at threshold of *p* < .005. **e.** Visualization of GLMs. The tables depict the structure of GLMs1-4 which were mainly motivated by the behavioral analysis; onset regressors are shown in the top table, parametric modulators assigned to 1D and 2D onsets (middle-left), the values they were modeled with (demeaned, middle-right) are shown below. The contrasts of interest are shown in the bottom table. The GLMs differed only in their modulations of the 2D trials: GLM1 included only modulators of the objective outcome, GLM2 included one modulator for Congruency and one for EV_back_, GLM3 included a modulator for the Congruency × EV_back_ interaction and GLM4 included instead of the EV modulator a modulator of the EV × Congruency interaction. In the contrast table (bottom) contrasts that only revealed effects at a liberal threshold of *p* < .005 are marked with one star, and contrasts significant at *p* < .001 are marked with two stars. .

**Figure S10:**
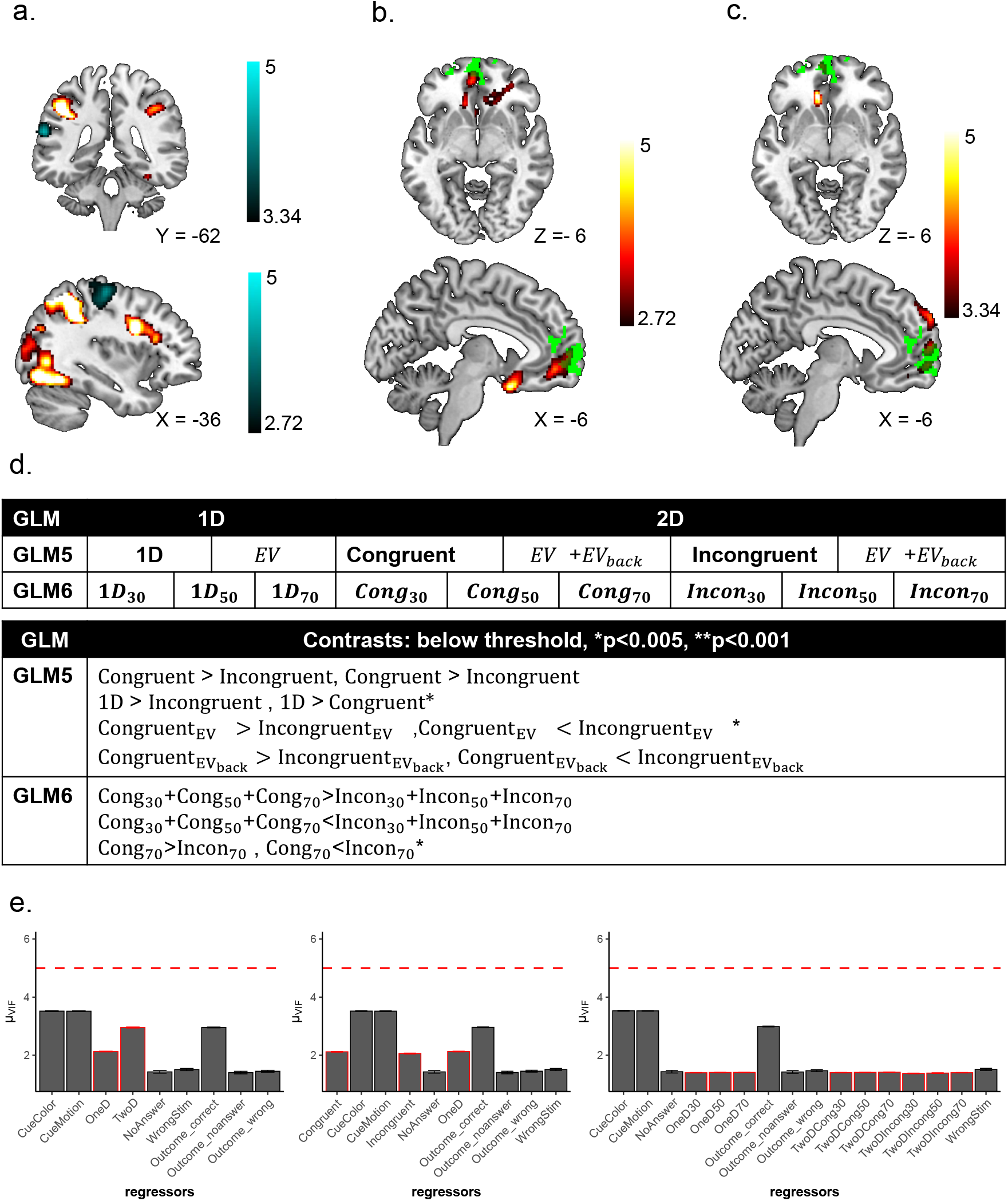
Additional univariate results. **a.** Overlap of effects of EV_back_ and trial type (2D > 1D). Main effects of EV_back_<0 (GLM2, *p* < 0.001 FDR cluster corrected, top, blue shades) and EV_back_ X Congruency < 0 (GLM3, *p* < 0.005, FDR cluster corrected, bottom, blue shades, t values) did not overlap with the 2D network (red shades in both panels, t values). **b.** Main effect of 1D > 2D. A stronger signal in vmPFC for 1D over 2D trials revealed weak activation in a PFC network (*p* < .005, red shades,t values). This included the vmPFC (our functional ROI is depicted in green). Interestingly, at a liberal threshold of *p* < .005 we found stronger activity for 1D over 2D trials in a cluster overlapping with vmPFC (1D > 2D, *p* < .005). Although this could be interpreted as a general preference for 1D trials, splitting the 2D onsets by Congruency revealed no cluster for 1D > Incongruent (also at *p* < .005) but a stronger cluster for 1D > Congruent (*p* < .001, Fig. S10). In other words, the signal in the vmPFC was *weaker* when both contexts indicate the same action, compared to when only one context is present. **c.** Stronger signal in vmPFC for 1D over congruent but not incongruent trials. When we split the onset of the 2D into Congruent and Incongruent trials (GLM5), we found no significant cluster for the 1D > Incongruent contrast, but an overlapping and stronger cluster for the 1D > Congruent contrast (*p* < .001, FDR cluster corrected, red shades, t values). We found very similar results when contrasting the onsets of 1D and Congruent in GLM6 (not presented), confirming the same results also when controlling for the number of trials for each level of EV (i.e. 1D_30_+1D_50_+1D_70_> Congruent_30_+Congruent_50_+Congruent_70_). Our functional ROI is depicted in green. **d.** Additional exploratory analyses such as contrasting the onsets of congruent and incongruent trials, confirmed the lack of Congruency modulation in any frontal region. Specifically, We constructed additional GLMs to verify the results of GLMs 1-4. In GLM5 we split the onset of 2D trials into congruent and incongruent trials and assigned a parametric modulator of EV and EV_back_ to each. As in GLM2, we found no effect of congruency; no voxel survived when contrasting the congruency onsets nor their EV_back_ modulators. Only the contrast Congruent_*EV*_ <Incongruent_*EV*_ revealed a weak cluster in the right visual cortex (peak 38,−80,16, p<0.005 not presented). In GLM6 we split the onsets of the 1D and 2D trials by levels of EV and the 2D trials further by Congruency. No Congruency main effect survived correction. Only when the onsets of Congruent and Incongruent 2D trials with EV=70 were contrasted, a cluster in the primary motor cortex was found (also at *p* < .005). Unsurprisingly, this cluster largely overlapped with the Congruency × EV_back_ effect reported in the Main Text. Except the contrast of 1D > Congruent (see Main Text) none of the other contrasts shown in the table revealed any cluster, even at *p* < .005. **e.** Variance Inflation Factor (VIF) of the different regressors in all GLMs. None of the regressors (x axis) had a mean VIF value (y axis) across blocks and participants above the threshold of 4. Regressors involved in GLMs 1-4 shown on the left (Fig. S9); GLM5 and GLM6 are shown in the middle and on the right, respectively. See Online Methods for details.

**Table S1:**
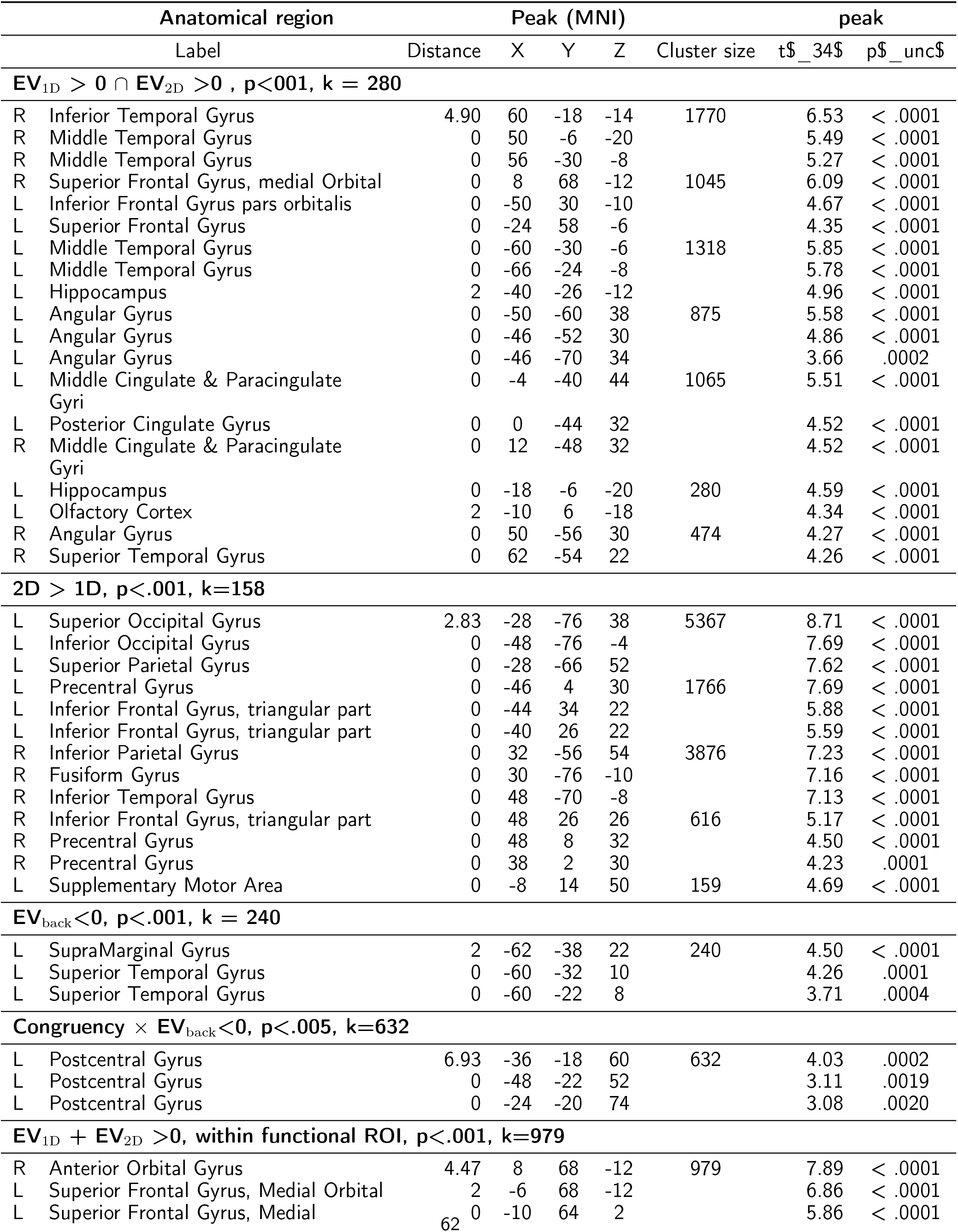
Detailed univariate results: Clusters for whole brain univariate analysis, related to Fig. S9. Presented are the closest labels to the local maxima of each cluster and each contrast using AAL3v1 [87–89]. All contrasts are FDR cluster corrected. p and k values presented for each cluster.

## Notes

### Competing Interest Statement

The authors have declared no competing interest.

## References

[1] Daniel Kahneman and Amos Tversky. Prospect theory : An analysis of decisions under risk. Econometrica, 47:278, 1979.

[2] John O’Doherty, Morten L Kringelbach, Edmund T Rolls, Julia Hornak, and Caroline Andrews. Abstract reward and punishment representations in the human orbitofrontal cortex. Nature neuroscience, 4(1):95–102, 2001.

[3] Camillo Padoa-Schioppa and John A Assad. Neurons in the orbitofrontal cortex encode economic value. Nature, 441 (7090):223–226, 2006.

[4] Oscar Bartra, Joseph T McGuire, and Joseph W Kable. The valuation system: a coordinate-based meta-analysis of bold fmri experiments examining neural correlates of subjective value. Neuroimage, 76:412–427, 2013. ISSN 1053-8119.

[5] John A Clithero and Antonio Rangel. Informatic parcellation of the network involved in the computation of subjective value. Social cognitive and affective neuroscience, 9(9):1289–1302, 2014.

[6] Hilke Plassmann, John O’doherty, and Antonio Rangel. Orbitofrontal cortex encodes willingness to pay in everyday economic transactions. Journal of neuroscience, 27(37):9984–9988, 2007.

[7] Erin L Rich and Jonathan D Wallis. Decoding subjective decisions from orbitofrontal cortex. Nature neuroscience, 19 (7):973–980, 2016.

[8] Sebastien Ballesta, Weikang Shi, Katherine E Conen, and Camillo Padoa-Schioppa. Values encoded in orbitofrontal cortex are causally related to economic choices. Nature, 2020.

[9] Maurizio Corbetta and Gordon L Shulman. Control of goal-directed and stimulus-driven attention in the brain. Nature reviews neuroscience, 3(3):201–215, 2002.

[10] Mark G Stokes, Makoto Kusunoki, Natasha Sigala, Hamed Nili, David Gaffan, and John Duncan. Dynamic coding for cognitive control in prefrontal cortex. Neuron, 78(2):364–375, 2013.

[11] Yael Niv, Reka Daniel, Andra Geana, Samuel J Gershman, Yuan Chang Leong, Angela Radulescu, and Robert C Wilson. Reinforcement learning in multidimensional environments relies on attention mechanisms. Journal of Neuroscience, 35 (21):8145–8157, 2015. ISSN 0270-6474.

[12] Yuan Chang Leong, Angela Radulescu, Reka Daniel, Vivian DeWoskin, and Yael Niv. Dynamic interaction between reinforcement learning and attention in multidimensional environments. Neuron, 93(2):451–463, 2017.

[13] Peter H Rudebeck and Elisabeth A Murray. The orbitofrontal oracle: cortical mechanisms for the prediction and evaluation of specific behavioral outcomes. Neuron, 84(6):1143–1156, 2014.

[14] Romy Frömer, Carolyn K Dean Wolf, and Amitai Shenhav. Goal congruency dominates reward value in accounting for behavioral and neural correlates of value-based decision-making. Nature communications, 10(1):1–11, 2019.

[15] G Castegnetti, M Zurita, and B De Martino. How usefulness shapes neural representations during goal-directed behavior. Science Advances, 7(15):eabd5363, 2021.

[16] Vikram S. Chib, Antonio Rangel, Shinsuke Shimojo, and John P. O’Doherty. Evidence for a common representation of decision values for dissimilar goods in human ventromedial prefrontal cortex. Journal of Neuroscience, 29(39): 12315–12320, 2009. ISSN 0270-6474. doi: 10.1523/JNEUROSCI.2575-09.2009.

[17] Daniel McNamee, Antonio Rangel, and John P O’doherty. Category-dependent and category-independent goal-value codes in human ventromedial prefrontal cortex. Nature neuroscience, 16(4):479–485, 2013. ISSN 1097-62656.

[18] Gabriel Pelletier and Lesley K Fellows. A critical role for human ventromedial frontal lobe in value comparison of complex objects based on attribute configuration. Journal of Neuroscience, 39(21):4124–4132, 2019.

[19] Thorsten Kahnt, Jakob Heinzle, Soyoung Q Park, and John-Dylan Haynes. Decoding different roles for vmpfc and dlpfc in multi-attribute decision making. Neuroimage, 56(2):709–715, 2011.

[20] Ulrike Basten, Guido Biele, Hauke R. Heekeren, and Christian J. Fiebach. How the brain integrates costs and benefits during decision making. Proceedings of the National Academy of Sciences, 107(50):21767–21772, 2010. doi: 10.1073/pnas.0908104107.

[21] Amitai Shenhav, Mark A Straccia, Sebastian Musslick, Jonathan D Cohen, and Matthew M Botvinick. Dissociable neural mechanisms track evidence accumulation for selection of attention versus action. Nature communications, 9, 2018.

[22] Nitzan Shahar, Rani Moran, Tobias U Hauser, Rogier A Kievit, Daniel McNamee, Michael Moutoussis, Raymond J Dolan, NSPN Consortium, et al. Credit assignment to state-independent task representations and its relationship with model-based decision making. Proceedings of the National Academy of Sciences, 116(32):15871–15876, 2019.

[23] Vickie Li, Elizabeth Michael, Jan Balaguer, Santiago Herce Castañón, and Christopher Summerfield. Gain control explains the effect of distraction in human perceptual, cognitive, and economic decision making. Proceedings of the National Academy of Sciences, 115(38):E8825–E8834, 2018. ISSN 0027-8424.

[24] Valerio Mante, David Sussillo, Krishna V Shenoy, and William T Newsome. Context-dependent computation by recurrent dynamics in prefrontal cortex. nature, 503(7474):78, 2013. ISSN 1476-4687.

[25] Yu Takagi, Laurence T Hunt, Mark W Woolrich, Timothy EJ Behrens, and Miriam Klein-Flugge. Projections of non-invasive human recordings into state space show unfolding of spontaneous and over-trained choice. bioRxiv, 2020.

[26] Nicolas W Schuck, Robert Gaschler, Dorit Wenke, Jakob Heinzle, Peter A Frensch, John-Dylan Haynes, and Carlo Reverberi. Medial prefrontal cortex predicts internally driven strategy shifts. Neuron, 86(1):331–340, 2015.

[27] Brian A Anderson. A value-driven mechanism of attentional selection. Journal of vision, 13(3):7–7, 2013.

[28] Marcus Grueschow, Rafael Polania, Todd A Hare, and Christian C Ruff. Automatic versus choice-dependent value representations in the human brain. Neuron, 85(4):874–885, 2015.

[29] Maël Lebreton, Soledad Jorge, Vincent Michel, Bertrand Thirion, and Mathias Pessiglione. An automatic valuation system in the human brain: Evidence from functional neuroimaging. Neuron, 64(3):431–439, 2009. ISSN 0896-6273. doi: https://doi.org/10.1016/j.neuron.2009.09.040.

[30] Nicolas W Schuck, Ming Bo Cai, Robert C Wilson, and Yael Niv. Human orbitofrontal cortex represents a cognitive map of state space. Neuron, 91(6):1402–1412, 2016.

[31] Stephanie CY Chan, Yael Niv, and Kenneth A Norman. A probability distribution over latent causes, in the orbitofrontal cortex. Journal of Neuroscience, 36(30):7817–7828, 2016.

[32] Nicolas W Schuck, Robert Wilson, and Yael Niv. A state representation for reinforcement learning and decision-making in the orbitofrontal cortex. In Goal-directed decision making, pages 259–278. Elsevier, 2018.

[33] G Elliott Wimmer and Christian Büchel. Learning of distant state predictions by the orbitofrontal cortex in humans. Nature communications, 10(1):1–11, 2019.

[34] Margaret L Schlichting, Jeanette A Mumford, and Alison R Preston. Learning-related representational changes reveal dissociable integration and separation signatures in the hippocampus and prefrontal cortex. Nature communications, 6 (1):1–10, 2015.

[35] Geoffrey Schoenbaum and Matthew Roesch. Orbitofrontal cortex, associative learning, and expectancies. Neuron, 47 (5):633–636, 2005.

[36] Michael L Mack, Alison R Preston, and Bradley C Love. Ventromedial prefrontal cortex compression during concept learning. Nature communications, 11(1):1–11, 2020.

[37] Christian F Doeller, Caswell Barry, and Neil Burgess. Evidence for grid cells in a human memory network. Nature, 463 (7281):657–661, 2010.

[38] Alexandra O Constantinescu, Jill X O’Reilly, and Timothy EJ Behrens. Organizing conceptual knowledge in humans with a gridlike code. Science, 352(6292):1464–1468, 2016.

[39] Jingfeng Zhou, Matthew PH Gardner, Thomas A Stalnaker, Seth J Ramus, Andrew M Wikenheiser, Yael Niv, and Geoffrey Schoenbaum. Rat orbitofrontal ensemble activity contains multiplexed but dissociable representations of value and task structure in an odor sequence task. Current Biology, 29(6):897–907, 2019.

[40] Anja Farovik, Ryan J Place, Sam McKenzie, Blake Porter, Catherine E Munro, and Howard Eichenbaum. Orbitofrontal cortex encodes memories within value-based schemas and represents contexts that guide memory retrieval. Journal of Neuroscience, 35(21):8333–8344, 2015.

[41] Praveen K Pilly and Aaron R Seitz. What a difference a parameter makes: A psychophysical comparison of random dot motion algorithms. Vision research, 49(13):1599–1612, 2009. ISSN 0042-6989.

[42] Denis Cousineau et al. Confidence intervals in within-subject designs: A simpler solution to loftus and masson’s method. Tutorials in quantitative methods for psychology, 1(1):42–45, 2005.

[43] Richard D Morey et al. Confidence intervals from normalized data: A correction to cousineau (2005). reason, 4(2): 61–64, 2008.

[44] Laurence T Hunt, Nils Kolling, Alireza Soltani, Mark W Woolrich, Matthew FS Rushworth, and Timothy EJ Behrens. Mechanisms underlying cortical activity during value-guided choice. Nature neuroscience, 15(3):470–476, 2012. ISSN 1097-6256.

[45] Satohiro Tajima, Jan Drugowitsch, and Alexandre Pouget. Optimal policy for value-based decision-making. Nature communications, 7(1):1–12, 2016.

[46] Ian Krajbich, Björn Bartling, Todd Hare, and Ernst Fehr. Rethinking fast and slow based on a critique of reaction-time reverse inference. Nature Communications, 6:7455, 2015. doi: 10.1038/ncomms8455. URL https://doi.org/10.1038/ncomms8455.

[47] Arni Magnusson, Hans Skaug, Anders Nielsen, Casper Berg, Kasper Kristensen, Martin Maechler, Koen van Bentham, Ben Bolker, Mollie Brooks, and Maintainer Mollie Brooks. Package ‘glmmtmb’. R Package Version 0.2. 0, 2017.

[48] Robert C Wilson, Yuji K Takahashi, Geoffrey Schoenbaum, and Yael Niv. Orbitofrontal cortex as a cognitive map of task space. Neuron, 81(2):267–279, 2014.

[49] Yael Niv. Learning task-state representations. Nature neuroscience, 22(10):1544–1553, 2019.

[50] Arno Klein, Satrajit S. Ghosh, Forrest S. Bao, Joachim Giard, Yrjö Häme, Eliezer Stavsky, Noah Lee, Brian Rossa, Martin Reuter, Elias Chaibub Neto, and Anisha Keshavan. Mindboggling morphometry of human brains. PLOS Computational Biology, 13(2):e1005350, 2017. ISSN 1553-7358. doi: 10.1371/journal.pcbi.1005350. URL http://journals.plos.org/ploscompbiol/article?id=10.1371/journal.pcbi.1005350.

[51] Anders M. Dale, Bruce Fischl, and Martin I. Sereno. Cortical surface-based analysis: I. segmentation and surface reconstruction. NeuroImage, 9(2):179–194, 1999. ISSN 1053-8119. doi: 10.1006/nimg.1998.0395. URL http://www.sciencedirect.com/science/article/pii/S1053811998903950.

[52] David H Brainard and Spatial Vision. The psychophysics toolbox. Spatial vision, 10:433–436, 1997.

[53] Denis G Pelli and Spatial Vision. The videotoolbox software for visual psychophysics: Transforming numbers into movies. Spatial vision, 10:437–442, 1997.

[54] Mario Kleiner, David Brainard, Denis Pelli, Allen Ingling, Richard Murray, Christopher Broussard, et al. What’s new in psychtoolbox-3. Perception, 36(14):1, 2007.

[55] MATLAB version 9.3.0.713579 (R2017b). The Mathworks, Inc., Natick, Massachusetts, 2017.

[56] MATLAB version (R2012b). The Mathworks, Inc., Natick, Massachusetts, 2017.

[57] Joshua T Abbott, Thomas L Griffiths, and Terry Regier. Focal colors across languages are representative members of color categories. Proceedings of the National Academy of Sciences, 113(40):11178–11183, 2016. ISSN 0027-8424.

[58] Helen C Barron, Mona M Garvert, and Timothy EJ Behrens. Repetition suppression: a means to index neural representations using bold? Philosophical Transactions of the Royal Society B: Biological Sciences, 371(1705): 20150355, 2016.

[59] Mona M Garvert, Raymond J Dolan, and Timothy EJ Behrens. A map of abstract relational knowledge in the human hippocampal–entorhinal cortex. Elife, 6:e17086, 2017.

[60] R Core Team. R: A Language and Environment for Statistical Computing. R Foundation for Statistical Computing, Vienna, Austria, 2017. URL https://www.R-project.org/.

[61] RStudio Team. RStudio: Integrated Development Environment for R. RStudio, PBC., Boston, MA, 2020. URL http://www.rstudio.com/.

[62] Douglas Bates, Martin Mächler, Ben Bolker, and Steve Walker. Fitting linear mixed-effects models using lme4. Journal of Statistical Software, 67(1):1–48, 2015. doi: 10.18637/jss.v067.i01.

[63] Nikolaus Weiskopf, Chloe Hutton, Oliver Josephs, and Ralf Deichmann. Optimal epi parameters for reduction of susceptibility-induced bold sensitivity losses: A whole-brain analysis at 3 t and 1.5 t. NeuroImage, 33(2):493–504, 2006. ISSN 1053-8119. doi: https://doi.org/10.1016/j.neuroimage.2006.07.029.

[64] Krzysztof J Gorgolewski, Tibor Auer, Vince D Calhoun, R Cameron Craddock, Samir Das, Eugene P Duff, Guillaume Flandin, Satrajit S Ghosh, Tristan Glatard, Yaroslav O Halchenko, et al. The brain imaging data structure, a format for organizing and describing outputs of neuroimaging experiments. Scientific data, 3(1):1–9, 2016.

[65] Xiangrui Li, Paul S Morgan, John Ashburner, Jolinda Smith, and Christopher Rorden. The first step for neuroimaging data analysis: Dicom to nifti conversion. Journal of neuroscience methods, 264:47–56, 2016.

[66] Oscar Esteban, Daniel Birman, Marie Schaer, Oluwasanmi O Koyejo, Russell A Poldrack, and Krzysztof J Gorgolewski. Mriqc: Advancing the automatic prediction of image quality in mri from unseen sites. PloS one, 12(9):e0184661, 2017.

[67] Oscar Esteban, Christopher Markiewicz, Ross W Blair, Craig Moodie, Ayse Ilkay Isik, Asier Erramuzpe Aliaga, James Kent, Mathias Goncalves, Elizabeth DuPre, Madeleine Snyder, Hiroyuki Oya, Satrajit Ghosh, Jessey Wright, Joke Durnez, Russell Poldrack, and Krzysztof Jacek Gorgolewski. fMRIPrep: a robust preprocessing pipeline for functional MRI. Nature Methods, 2018. doi: 10.1038/s41592-018-0235-4.

[68] Oscar Esteban, Ross Blair, Christopher J. Markiewicz, Shoshana L. Berleant, Craig Moodie, Feilong Ma, Ayse Ilkay Isik, Asier Erramuzpe, Mathias Kent, James D. andGoncalves, Elizabeth DuPre, Kevin R. Sitek, Daniel E. P. Gomez, Daniel J. Lurie, Zhifang Ye, Russell A. Poldrack, and Krzysztof J. Gorgolewski. fmriprep. Software, 2018. doi: 10.5281/zenodo.852659.

[69] K. Gorgolewski, C. D. Burns, C. Madison, D. Clark, Y. O. Halchenko, M. L. Waskom, and S. Ghosh. Nipype: a flexible, lightweight and extensible neuroimaging data processing framework in python. Frontiers in Neuroinformatics, 5:13, 2011. doi: 10.3389/fninf.2011.00013.

[70] Krzysztof J. Gorgolewski, Oscar Esteban, Christopher J. Markiewicz, Erik Ziegler, David Gage Ellis, Michael Philipp Notter, Dorota Jarecka, Hans Johnson, Christopher Burns, Alexandre Manhães-Savio, Carlo Hamalainen, Benjamin Yvernault, Taylor Salo, Kesshi Jordan, Mathias Goncalves, Michael Waskom, Daniel Clark, Jason Wong, Fred Loney, Marc Modat, Blake E Dewey, Cindee Madison, Matteo Visconti di Oleggio Castello, Michael G. Clark, Michael Dayan, Dav Clark, Anisha Keshavan, Basile Pinsard, Alexandre Gramfort, Shoshana Berleant, Dylan M. Nielson, Salma Bougacha, Gael Varoquaux, Ben Cipollini, Ross Markello, Ariel Rokem, Brendan Moloney, Yaroslav O. Halchenko, Demian Wassermann, Michael Hanke, Christian Horea, Jakub Kaczmarzyk, Gilles de Hollander, Elizabeth DuPre, Ashley Gillman, David Mordom, Colin Buchanan, Rosalia Tungaraza, Wolfgang M. Pauli, Shariq Iqbal, Sharad Sikka, Matteo Mancini, Yannick Schwartz, Ian B. Malone, Mathieu Dubois, Caroline Frohlich, David Welch, Jessica Forbes, James Kent, Aimi Watanabe, Chad Cumba, Julia M. Huntenburg, Erik Kastman, B. Nolan Nichols, Arman Eshaghi, Daniel Ginsburg, Alexander Schaefer, Benjamin Acland, Steven Giavasis, Jens Kleesiek, Drew Erickson, René Küttner, Christian Haselgrove, Carlos Correa, Ali Ghayoor, Franz Liem, Jarrod Millman, Daniel Haehn, Jeff Lai, Dale Zhou, Ross Blair, Tristan Glatard, Mandy Renfro, Siqi Liu, Ari E. Kahn, Fernando Pérez-García, William Triplett, Leonie Lampe, Jörg Stadler, Xiang-Zhen Kong, Michael Hallquist, Andrey Chetverikov, John Salvatore, Anne Park, Russell Poldrack, R. Cameron Craddock, Souheil Inati, Oliver Hinds, Gavin Cooper, L. Nathan Perkins, Ana Marina, Aaron Mattfeld, Maxime Noel, Lukas Snoek, K Matsubara, Brian Cheung, Simon Rothmei, Sebastian Urchs, Joke Durnez, Fred Mertz, Daniel Geisler, Andrew Floren, Stephan Gerhard, Paul Sharp, Miguel Molina-Romero, Alejandro Weinstein, William Broderick, Victor Saase, Sami Kristian Andberg, Robbert Harms, Kai Schlamp, Jaime Arias, Dimitri Papadopoulos Orfanos, Claire Tarbert, Arielle Tambini, Alejandro De La Vega, Thomas Nickson, Matthew Brett, Marcel Falkiewicz, Kornelius Podranski, Janosch Linkersdörfer, Guillaume Flandin, Eduard Ort, Dmitry Shachnev, Daniel McNamee, Andrew Davison, Jan Varada, Isaac Schwabacher, John Pellman, Martin Perez-Guevara, Ranjeet Khanuja, Nicolas Pannetier, Conor McDermottroe, and Satrajit Ghosh. Nipype. Software, 2018. doi: 10.5281/zenodo.596855.

[71] Alexandre Abraham, Fabian Pedregosa, Michael Eickenberg, Philippe Gervais, Andreas Mueller, Jean Kossaifi, Alexandre Gramfort, Bertrand Thirion, and Gael Varoquaux. Machine learning for neuroimaging with scikit-learn. Frontiers in Neuroinformatics, 8, 2014. ISSN 1662-5196. doi: 10.3389/fninf.2014.00014. URL https://www.frontiersin.org/articles/10.3389/fninf.2014.00014/full.

[72] N. J. Tustison, B. B. Avants, P. A. Cook, Y. Zheng, A. Egan, P. A. Yushkevich, and J. C. Gee. N4itk: Improved n3 bias correction. IEEE Transactions on Medical Imaging, 29(6):1310–1320, 2010. ISSN 0278-0062. doi: 10.1109/TMI.2010.2046908.

[73] VS Fonov, AC Evans, RC McKinstry, CR Almli, and DL Collins. Unbiased nonlinear average age-appropriate brain templates from birth to adulthood. NeuroImage, 47, Supplement 1:S102, 2009. ISSN 1053-8119. doi: 10.1016/S1053-8119(09)70884-5. URL http://www.sciencedirect.com/science/article/pii/S1053811909708845.

[74] B.B. Avants, C.L. Epstein, M. Grossman, and J.C. Gee. Symmetric diffeomorphic image registration with cross-correlation: Evaluating automated labeling of elderly and neurodegenerative brain. Medical Image Analysis, 12(1): 26–41, 2008. ISSN 1361-8415. doi: 10.1016/j.media.2007.06.004. URL http://www.sciencedirect.com/science/article/pii/S1361841507000606.

[75] Y. Zhang, M. Brady, and S. Smith. Segmentation of brain MR images through a hidden markov random field model and the expectation-maximization algorithm. IEEE Transactions on Medical Imaging, 20(1):45–57, 2001. ISSN 0278-0062. doi: 10.1109/42.906424.

[76] Robert W. Cox and James S. Hyde. Software tools for analysis and visualization of fmri data. NMR in Biomedicine, 10(4-5):171–178, 1997. doi: 10.1002/(SICI)1099-1492(199706/08)10:4/5<171::AID-NBM453>3.0.CO;2-L.

[77] Douglas N Greve and Bruce Fischl. Accurate and robust brain image alignment using boundary-based registration. NeuroImage, 48(1):63–72, 2009. ISSN 1095-9572. doi: 10.1016/j.neuroimage.2009.06.060.

[78] Mark Jenkinson, Peter Bannister, Michael Brady, and Stephen Smith. Improved optimization for the robust and accurate linear registration and motion correction of brain images. NeuroImage, 17(2):825–841, 2002. ISSN 1053-8119. doi: 10.1006/nimg.2002.1132. URL http://www.sciencedirect.com/science/article/pii/S1053811902911328.

[79] Jonathan D. Power, Anish Mitra, Timothy O. Laumann, Abraham Z. Snyder, Bradley L. Schlaggar, and Steven E. Petersen. Methods to detect, characterize, and remove motion artifact in resting state fmri. NeuroImage, 84(Supplement C):320–341, 2014. ISSN 1053-8119. doi: 10.1016/j.neuroimage.2013.08.048. URL http://www.sciencedirect.com/science/article/pii/S1053811913009117.

[80] Yashar Behzadi, Khaled Restom, Joy Liau, and Thomas T. Liu. A component based noise correction method (CompCor) for BOLD and perfusion based fmri. NeuroImage, 37(1):90–101, 2007. ISSN 1053-8119. doi: 10.1016/j.neuroimage.2007.04.042. URL http://www.sciencedirect.com/science/article/pii/S1053811907003837.

[81] C. Lanczos. Evaluation of noisy data. Journal of the Society for Industrial and Applied Mathematics Series B Numerical Analysis, 1(1):76–85, 1964. ISSN 0887-459X. doi: 10.1137/0701007. URL http://epubs.siam.org/doi/10.1137/0701007.

[82] Gary H Glover, Tie-Qiang Li, and David Ress. Image-based method for retrospective correction of physiological motion effects in fmri: Retroicor. Magnetic Resonance in Medicine: An Official Journal of the International Society for Magnetic Resonance in Medicine, 44(1):162–167, 2000.

[83] Chloe Hutton, Oliver Josephs, Jörg Stadler, Eric Featherstone, Alphonso Reid, Oliver Speck, Johannes Bernarding, and Nikolaus Weiskopf. The impact of physiological noise correction on fmri at 7 t. Neuroimage, 57(1):101–112, 2011.

[84] Ann K Harvey, Kyle TS Pattinson, Jonathan CW Brooks, Stephen D Mayhew, Mark Jenkinson, and Richard G Wise. Brainstem functional magnetic resonance imaging: disentangling signal from physiological noise. Journal of Magnetic Resonance Imaging: An Official Journal of the International Society for Magnetic Resonance in Medicine, 28(6): 1337–1344, 2008.

[85] Lars Kasper, Steffen Bollmann, Andreea O Diaconescu, Chloe Hutton, Jakob Heinzle, Sandra Iglesias, Tobias U Hauser, Miriam Sebold, Zina-Mary Manjaly, Klaas P Pruessmann, et al. The physio toolbox for modeling physiological noise in fmri data. Journal of neuroscience methods, 276:56–72, 2017.

[86] William D Penny, Karl J Friston, John T Ashburner, Stefan J Kiebel, and Thomas E Nichols. Statistical parametric mapping: the analysis of functional brain images. Elsevier, 2011.

[87] Nathalie Tzourio-Mazoyer, Brigitte Landeau, Dimitri Papathanassiou, Fabrice Crivello, Olivier Etard, Nicolas Delcroix, Bernard Mazoyer, and Marc Joliot. Automated anatomical labeling of activations in spm using a macroscopic anatomical parcellation of the mni mri single-subject brain. Neuroimage, 15(1):273–289, 2002.

[88] Edmund T Rolls, Marc Joliot, and Nathalie Tzourio-Mazoyer. Implementation of a new parcellation of the orbitofrontal cortex in the automated anatomical labeling atlas. Neuroimage, 122:1–5, 2015.

[89] Edmund T Rolls, Chu-Chung Huang, Ching-Po Lin, Jianfeng Feng, and Marc Joliot. Automated anatomical labelling atlas 3. NeuroImage, 206:116189, 2020.

[90] Jeanette A Mumford, Jean-Baptiste Poline, and Russell A Poldrack. Orthogonalization of regressors in fmri models. PloS one, 10(4):e0126255, 2015.

[91] F. Pedregosa, G. Varoquaux, A. Gramfort, V. Michel, B. Thirion, O. Grisel, M. Blondel, P. Prettenhofer, R. Weiss, V. Dubourg, J. Vanderplas, A. Passos, D. Cournapeau, M. Brucher, M. Perrot, and E. Duchesnay. Scikit-learn: Machine learning in Python. Journal of Machine Learning Research, 12:2825–2830, 2011.

[92] Mollie E. Brooks, Kasper Kristensen, Koen J. van Benthem, Arni Magnusson, Casper W. Berg, Anders Nielsen, Hans J. Skaug, Martin Maechler, and Benjamin M. Bolker. glmmTMB balances speed and flexibility among packages for zero-inflated generalized linear mixed modeling. The R Journal, 9(2):378–400, 2017. URL https://journal.r-project.org/archive/2017/RJ-2017-066/index.html.

[93] Antoinette Nicolle, Miriam C Klein-Flügge, Laurence T Hunt, Ivo Vlaev, Raymond J Dolan, and Timothy EJ Behrens. An agent independent axis for executed and modeled choice in medial prefrontal cortex. Neuron, 75(6):1114–1121, 2012.

